# DNA-PK facilitates HIV transcription by regulating the activity of RNA polymerase II and the recruitment of transcription machinery at HIV LTR

**DOI:** 10.1101/174573

**Authors:** Sonia Zicari, Geetaram Sahu, Larisa Dubrovsky, Lin Sun, Han Yue, Tejaswi Jada, Alex Ochem, Michael Bukrinsky, Gary Simon, Mudit Tyagi

**Affiliations:** Division of Infectious Diseases, Department of Medicine, George Washington University, Washington, DC 20037, USA; Section of Intercellular Interactions, Eunice-Kennedy National Institute of Child Health and Human Development, National Institutes of Health, Bethesda, MD, USA; Department of Microbiology, Immunology and Tropical Medicine, George Washington University, Washington, DC 20037, USA; International Centre for Genetic Engineering and Biotechnology (ICGEB), Wernher and Beit Building (South), Anzio Road, Observatory 7925, Cape Town, South Africa

**Author notes:** First Co-authors (Sonia Zicari, Geetaram Sahu). **Corresponding author:** Mudit Tyagi, School of Medicine and Health Sciences, Ross Hall, Room 731, 2300 Eye Street, N.W., Washington, DC 20037.

**Keywords:** HIV, DNA-PK, Transcription, Replication, DNA-PK inhibitors, Latency reactivation

## Abstract

Despite the use of highly effective antiretroviral therapy (HAART), the presence of latent or transcriptionally silent proviruses prevents cure and eradication of HIV infection. These transcriptionally silent proviruses are well protected from both the immune system and HAART regimens. Thus, in order to tackle the problem of latent HIV reservoirs, it is a prerequisite to define all the pathways that regulate HIV transcription. We have previously reported that DNA-PK facilitates HIV transcription by interacting with the RNA polymerase II (RNAP II) complex recruited at HIV LTR. To extend those studies further, here we demonstrate that DNA-PK promotes HIV transcription by supporting it at several stages, including initiation, pause-release and elongation. We discovered that DNA-PK increases phosphorylation of RNAP II C-terminal domain (CTD) at serine 5 (Ser5) and serine 2 (Ser2) by both directly catalyzing and by augmenting the recruitment of P-TEFb at HIV LTR. We found that DNA-PK facilitates the establishment of euchromatin structure at HIV LTR, which further supports HIV gene expression. DNA-PK inhibition or knockdown leads to the severe impairment of HIV gene expression and conversion of euchromatin to heterochromatin at HIV LTR. It also profoundly restricts HIV replication and reactivation of latent provirus. DNA-PK promotes the recruitment of TRIM28 at LTR and facilitates the release of paused RNAP II through TRIM28 phosphorylation. The results were reproduced in cell lines belonging to both lymphoid and myeloid lineages and were confirmed in primary CD4^+^ T cells and peripheral blood mononuclear cells (PBMCs) from HIV-infected patients.

**IMPORTANCE:** Our results reveal the important role of DNA-PK in supporting HIV transcription, replication and latent proviral reactivation. Intriguingly, this study sheds light on an important pathway that affects HIV gene expression. These findings provide strong rationale for developing and using transcriptional inhibitors, such as DNA-PK inhibitors, as supplement to HAART regimens in order to further enhance their effectiveness and to suppress toxicity due to HIV proteins.

## INTRODUCTION

The present combination anti-retroviral therapy (cART), commonly known as highly active antiretroviral therapy (HAART), is able to suppress replication of human immunodeficiency virus (HIV) quite effectively, oftentimes with a single daily dose. However, when HAART is interrupted, even after several years of undetectable viremia, the virus resurrects out of latent reservoirs and quickly re-emerges in circulation (11, 14). Thus, the presence of latent or transcriptionally silent HIV provirus is a major hurdle to HIV eradication. The persistence of HIV in patients, despite prolonged treatment, has renewed interest in understanding the molecular mechanisms that control HIV life cycle. It is well accepted that the state of HIV transcription dictates the prevalence of either productively replicating or latent HIV proviruses in the cells (27, 38, 39, 53, 58, 59). The bigger pool of latent proviruses is incompetent in generating fully functional HIV progeny (4, 25). However, it has been documented that defective proviruses can still transcribe their genes and produce some of the viral proteins, which contribute to HIV-mediated cytotoxicity (46). Therefore, thorough understanding of the mechanisms that regulate HIV transcription is a prerequisite for effectively applying any strategy for HIV eradication or a functional cure.

Analogous to host gene transcription, HIV transcription is regulated by the well-controlled phosphorylation events of the carboxyl-terminal domain (CTD) of the largest subunit of the RNA polymerase II (RNAP II) (50). The mammalian RNAP II CTD consists of 52 tandem repeats of a consensus sequence Tyr1-Ser2-Pro3-Thr4-Ser5-Pro6-Ser7 (13, 42). More than half a dozen kinases are known to phosphorylate CTD of RNAP II. Among these, the importance of cyclin-dependent kinases (mainly CDK7 for phosphorylation of Serine 5 (Ser5), and CDK9 for phosphorylation of Serine 2 (Ser2)) during HIV transcription has already been established (13, 26, 50). We recently discovered that the DNA-dependent protein kinase (DNA-PK) is another kinase that phosphorylates the CTD of RNAP II and plays an important role during HIV transcription (61). We have previously documented the parallel presence of DNA-PK along with RNAP II throughout HIV genome, during HIV transcription. By performing *in vitro* kinase assays, we have shown that DNA-PK is able to phosphorylate all three serine residues (Ser2, Ser5 and Ser7) of the CTD region of RNAP II. We found that the transactivator of transcription (Tat) protein, which is vital for HIV transcription, is a potential substrate of DNA-PK. The finding that cellular activation enhances nuclear translocation of DNA-PK and its activation further supports our observation of greater DNA-PK recruitment at HIV long terminal repeat (LTR) following cellular activation (41, 61).

The human DNA-PK is a nuclear kinase that specifically requires association with DNA for its activity (2, 10, 16, 31). DNA-PK holoenzyme consists of two components: a 450 kDa catalytic subunit (DNA-PKcs) (23), which is a serine/threonine kinase, and a regulatory component known as Ku (20). Ku is a heterodimer comprised of two subunits, one 70 kDa (48) and another 80 kDa (68). The 70 kDa subunit possesses ATPase and DNA helicase activities. The vital role of DNA-PK in the non-homologous end joining (NHEJ) DNA-repair pathway is well-recognized (32, 55).

HIV transcription pauses after transcribing around first 60 bp (45, 49). RNAP II pausing is mainly attributed to the binding of negative elongation factor (NELF) and DRB sensitivity-inducing factor (DSIF) to HIV LTR (45, 47). Later, the Tat protein, by recruiting positive transcription elongation factor b (P-TEFb), relieves RNAP II pausing (5, 18). The CDK9 subunit of P-TEFb phosphorylates the NELF and DSIF subunits, which either converts them to a positive elongation factor or removes them from LTR (27). Transcriptional elongation needs the sequential specific phosphorylation events at RNAP II CTD in order to transform RNAP II to an elongating or processive enzyme.

Phosphorylation of Ser5 residue of the RNAP II CTD is linked to the initiation phase of transcription (28, 67), whereas phosphorylation of Ser2 is found to be correlated with the elongation phase of transcription, also during HIV gene expression (29, 45, 52). In addition to DSIF and NELF, recently another factor, the tripartite motif-containing 28 (known as TRIM28, KAP1, TIF1β), has been shown to support RNAP II pausing at certain cellular genes (6-8). Similar to the SPT5 subunit of DSIF (3), the phosphorylation of TRIM28 converts it from a pausing or negative elongation factor to a positive elongation factor (8, 33). DNA-PK is the principal kinase which directly interacts with TRIM28 and catalyzes the phosphorylation of TRIM28 at serine 824 residue converting it to an elongation factor (8). Pertaining HIV transcription, the role of TRIM28 is still not clear. However, the presence of TRIM28 bound with 7SK snRNP complex at HIV LTR has been documented (40). In addition to ours (61), other studies have also noted the interaction between RNAP II and DNA-PK (37). Moreover, we have shown that DNA-PK is a component of RNAP II holoenzyme, recruited at HIV LTR and it rides along RNAP II, throughout HIV genome (61). Recently, the interaction of TRIM28 with RNAP II and the continuous presence of TRIM28 with RNAP II along cellular genes’ body have been documented (7, 8). In this investigation, by attenuating the activity or cellular levels of DNA-PK, we have established the role of DNA-PK not only in activating TRIM28 through phosphorylation, but also in the recruitment of TRIM28 and phosphorylated TRIM28 (p-TRIM28, S824) at HIV LTR.

Several studies focusing on cancer therapy have targeted DNA-PK with small molecule inhibitors (15, 34, 69) in efforts to kill cancerous cells through accumulation of unrepaired DNA breaks induced by ionizing radiation (65, 72). For this purpose, several DNA-PK inhibitors have been developed, including LY294002, which binds to the kinase domain of DNA-PK (54). Subsequently, LY294002 became the template for the development of other more specific DNA-PK inhibitors, such as Nu7026 (2-(morpholin-4-yl)-benzo-H-chromen-4-one) that was shown to be 50-fold more specific for DNA-PK as compared to other kinases (44, 66). Another related compound, Nu7441 (2-N-(morpholino-8-benzothiophenyl)-chromen-4-one), was found to be even more selective for DNA-PK than Nu7026 (12, 15, 22, 30). Another class of DNA-PK inhibitors, such as 1-(2-hydroxy-4-morpholin-4-yl-phenyl)-ethanone (known as IC86621), differs structurally from other DNA-PK inhibitors. These highly specific, non-toxic competitive DNA-PK inhibitors are known to induce phenotype similar to that of DNA-PK deficient cell line and DNA-PK knockout mice (severe combined immunodeficiency, SCID, mice) (12, 15).

In prior publication we suggested an important role of DNA-PK during HIV transcription, as we found the continuous presence of DNA-PK with RNAP II at HIV LTR (61). In the current study, we sought to evaluate the mechanism through which DNA-PK promotes HIV transcription, replication and latent proviral reactivation. We explored the impact on HIV gene expression after inhibition or depletion of endogenous DNA-PK, by treating cells with either specific inhibitors or specific shRNA, respectively. We assessed the impact of DNA-PK on the phosphorylation state of RNAP II CTD and on the regulation of epigenetic changes at HIV LTR. The results were confirmed in various cell types belonging to different lineages, including in physiologically relevant primary T cells and cells from HIV-infected patients. We found that DNA-PK plays a major role during HIV transcription by supporting it at multiple steps. Consequently, DNA-PK inhibition or removal drastically impairs HIV transcription, replication and reactivation of latent provirus. In latently infected cells, where HIV provirus is transcriptionally silent, we noted highly reduced nuclear levels of DNA-PK. We have shown that DNA-PK promotes the HIV transcriptional initiation by catalyzing the phosphorylation of RNAP II CTD at Ser5; it relieves the pausing of RNAP II by catalyzing the phosphorylation of TRIM28 at S824. Finally, DNA-PK increases the elongation phase of HIV transcription by augmenting CTD phosphorylation, both by directly catalyzing and through co-recruiting P-TEFb at HIV LTR. We also evaluated the DNA-PK inhibitors as potential therapeutic agents in restricting HIV transcription, replication and proviral reactivation.

## Results

### DNA-PK inhibitors are strong repressors of HIV transcription, replication and reactivation of latent HIV provirus

We have shown previously that DNA-PK plays a significant role during HIV transcription (61). To extend those findings and in our quest to define the molecular mechanisms involved, we first evaluated the role of DNA-PK in supporting proviral reactivation from latency. Our hypothesis was that DNA-PK inhibition should impair HIV transcription, and consequently latent proviral reactivation. For proving this hypothesis, we assessed the reactivation of proviruses in latently infected cell lines and primary T cells in the presence of highly specific and well-established DNA-PK inhibitors. The latently infected THP-1, a human myeloid/monocytic cell line, Jurkat, a human lymphoid/T cell line, and primary CD4+ T cells were generated using our established methodologies (59, 60). The impact of DNA-PK inhibitors on the reactivation of integrated latent HIV provirus (pHR′P-Luc) expressing *luciferase* reporter gene under the control of the HIV LTR promoter (**Fig.1A**) was evaluated through luciferase assays. The cells were incubated overnight with various concentrations of different DNA-PK inhibitors or DMSO, as solvent control. The cells were stimulated with either Tumor Necrosis Factor alpha (TNF-α) or T cell receptor (TCR). A clear dose-dependent inhibition of HIV proviral reactivation by DNA-PK inhibitors was observed in all cell types, indicated by the reduced luciferase counts in cells treated with the inhibitors compared to the control **(Figs. 1B to D)**. The more specific DNA-PK inhibitors, Nu7441 and Nu7026, demonstrated better effect in restricting HIV gene expression than the less specific inhibitor IC86621. Since IC86621 was comparatively less efficient in restricting HIV transcription, in successive experiments we primarily used Nu7441 and Nu7026.

**Fig. 1:**
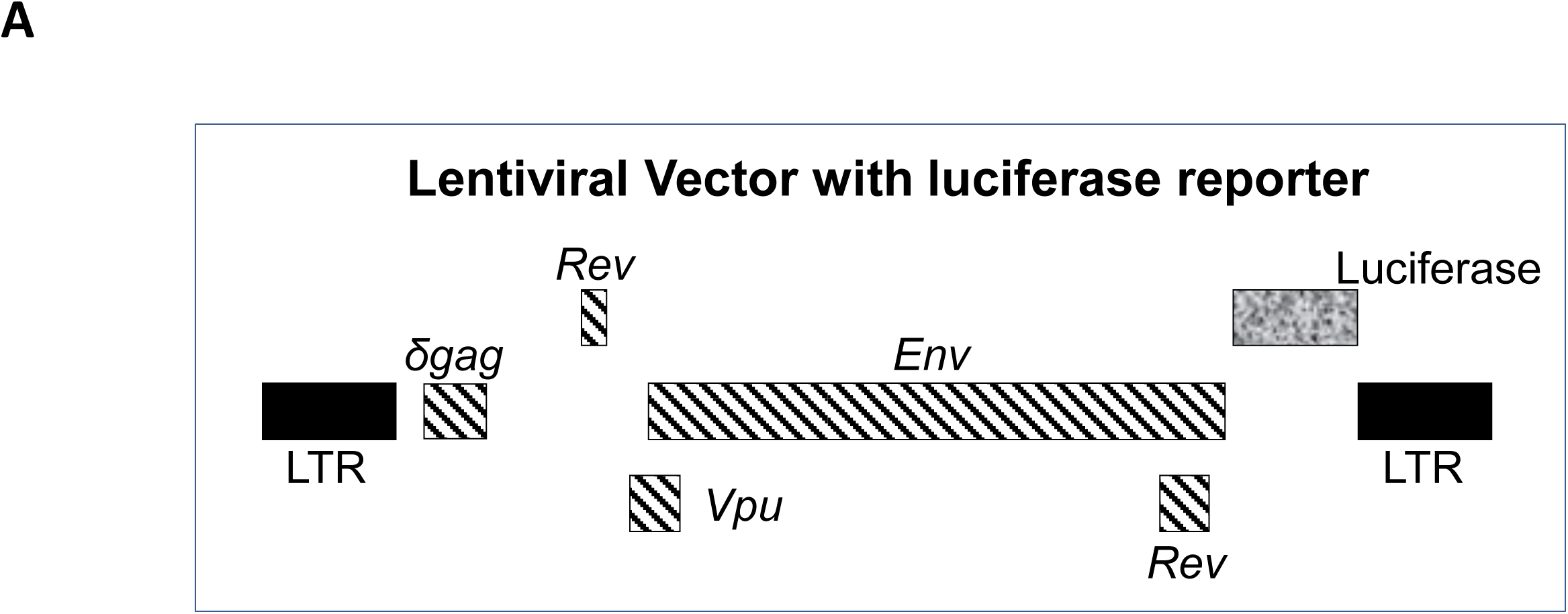
DNA-PK inhibitors repress HIV transcription without showing cytotoxicity and drastically impair latent proviral reactivation and HIV replication. (**A**) HIV-based lentiviral vector, pHR’P-Luc, which express *luciferase* reporter gene through HIV LTR promoter.

**Fig. 1:**
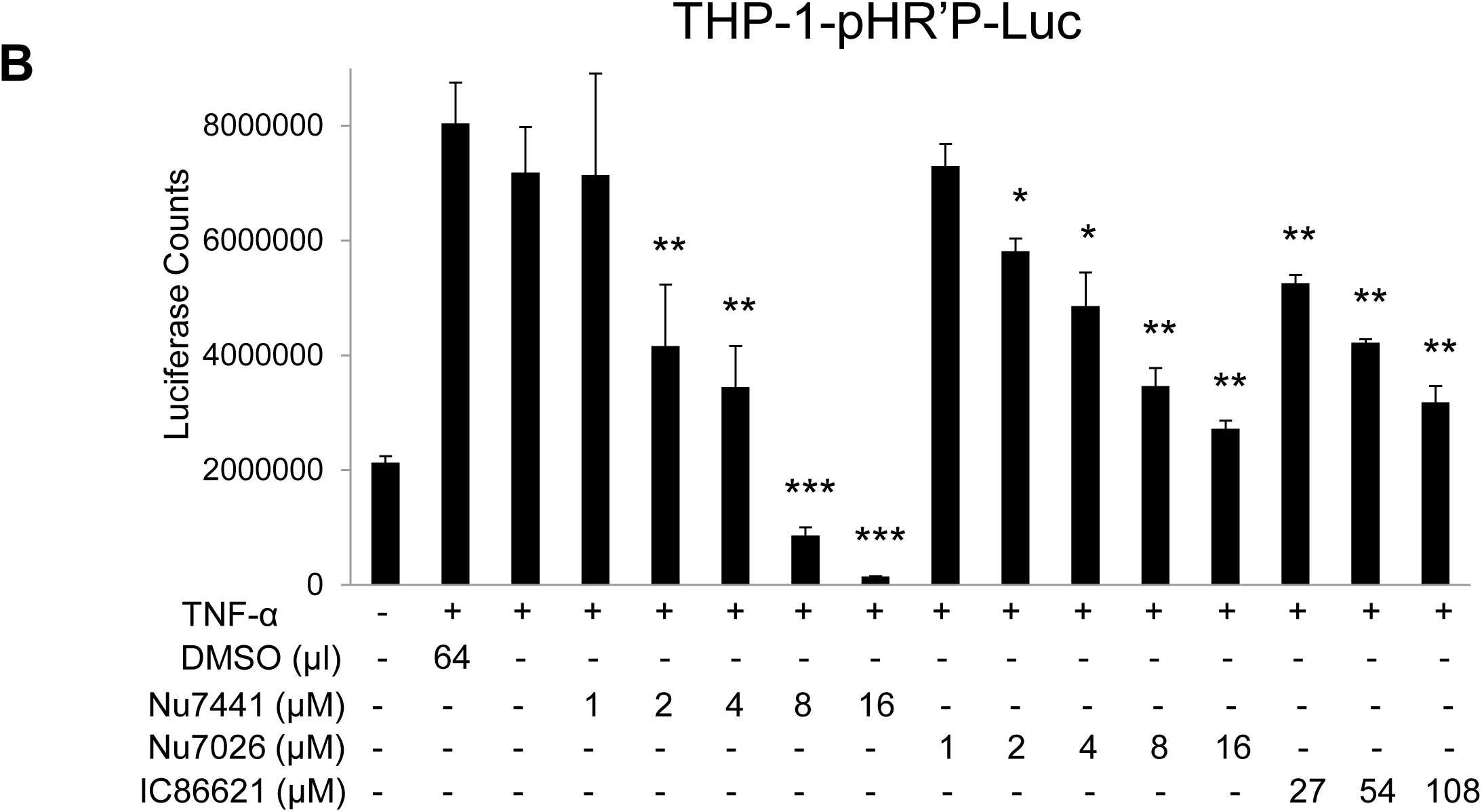
DNA-PK inhibitors repress HIV transcription without showing cytotoxicity and drastically impair latent proviral reactivation and HIV replication. The (**B**) THP-1, a monocytic cell line carrying integrated latent pHR’P-Luc provirus in their genome were treated with indicated amounts of the DNA-PK inhibitors Nu7441, Nu7026, IC86621 or DMSO solvent-control along with or without 10 ng/ml of TNF-α. After 48 hours cells were lysed and luciferase assays were performed using equal amounts of protein. The results represent the Mean ± SD of three different independent assays. The p value of statistical significance was set at either; p < 0.05 (*), 0.01 (**) or 0.001 (***).

**Fig. 1:**
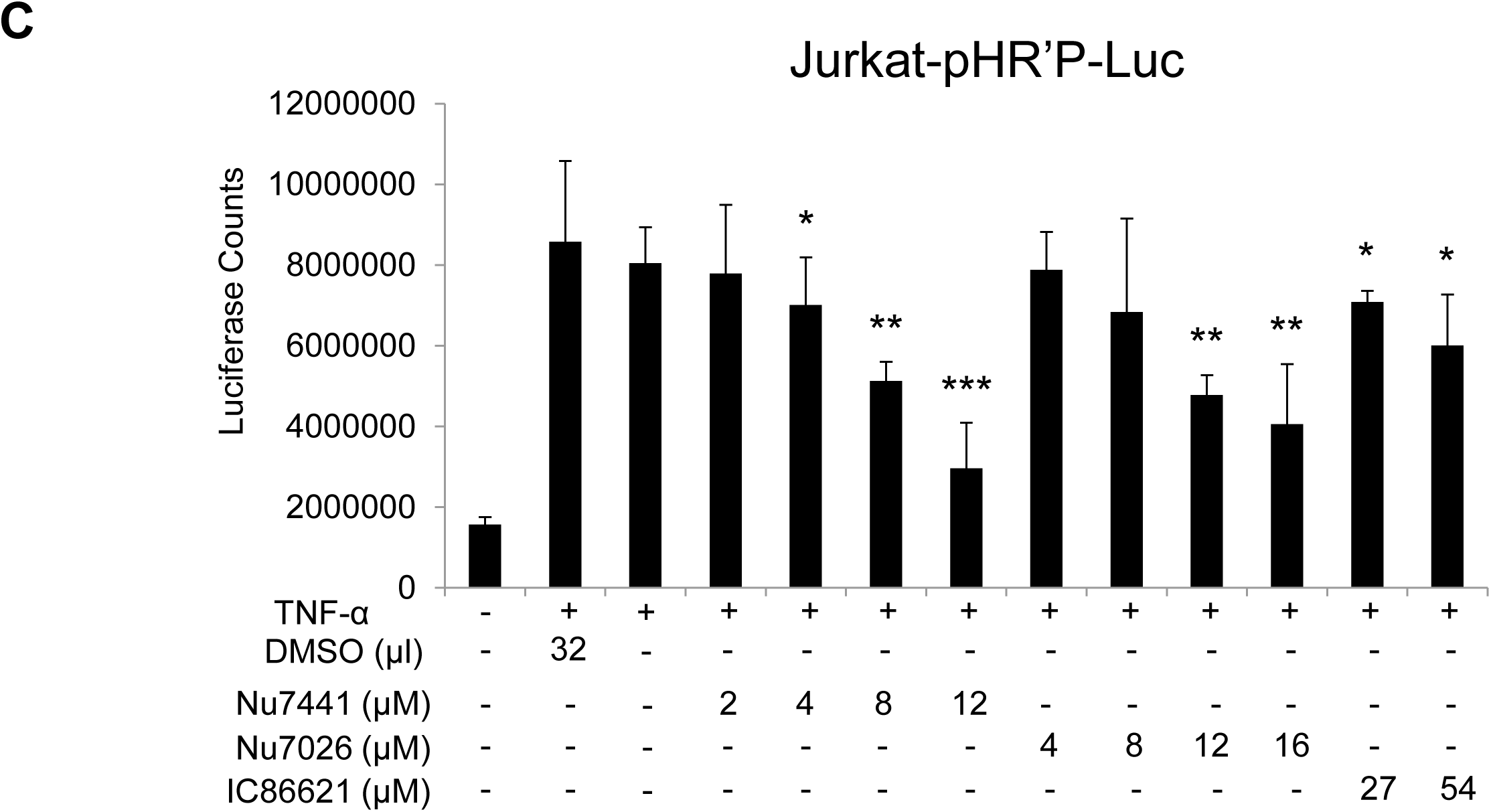
DNA-PK inhibitors repress HIV transcription without showing cytotoxicity and drastically impair latent proviral reactivation and HIV replication. The (**C**) Jurkat, a T cell line carrying integrated latent pHR’P-Luc provirus in their genome were treated with indicated amounts of the DNA-PK inhibitors Nu7441, Nu7026, IC86621 or DMSO solvent-control along with or without 10 ng/ml of TNF-α. After 48 hours cells were lysed and luciferase assays were performed. The results represent the Mean ± SD of three independent assays. The p value of statistical significance was set at either; p < 0.05 (*), 0.01 (**) or 0.001 (***).

**Fig. 1:**
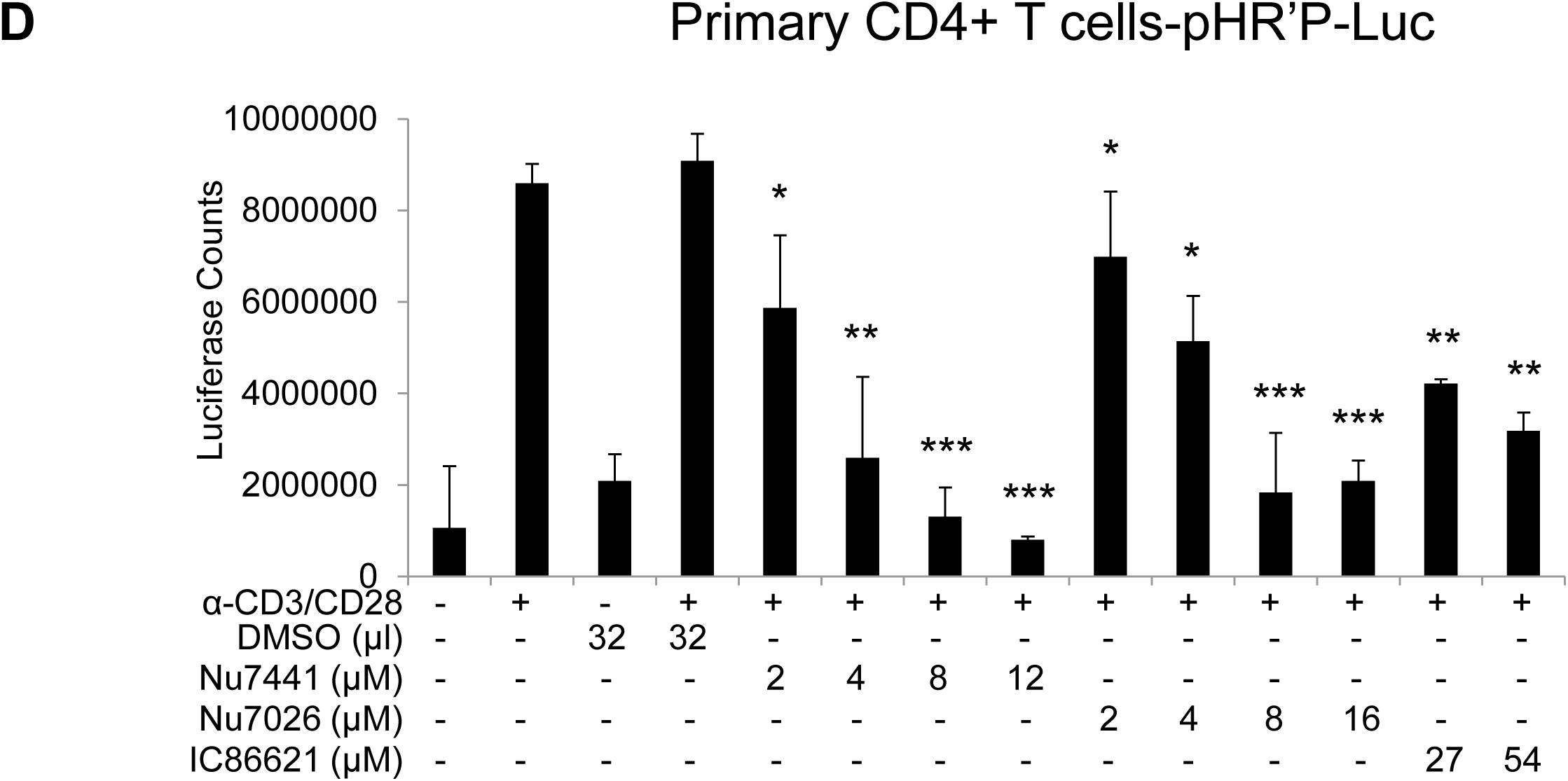
DNA-PK inhibitors repress HIV transcription without showing cytotoxicity and drastically impair latent proviral reactivation and HIV replication. The (**D**) primary CD4^+^ T cells carrying integrated latent pHR’P-Luc provirus in their genome were treated with indicated amounts of the DNA-PK inhibitors Nu7441, Nu7026, IC86621 or DMSO solvent-control along with or without 10 ng/ml of TNF-α. After 48 hours cells were lysed and luciferase assays were performed. The results represent the Mean ± SD of three independent assays. The p value of statistical significance was set at either; p < 0.05 (*), 0.01 (**), or 0.001 (***).

**Fig. 1:**
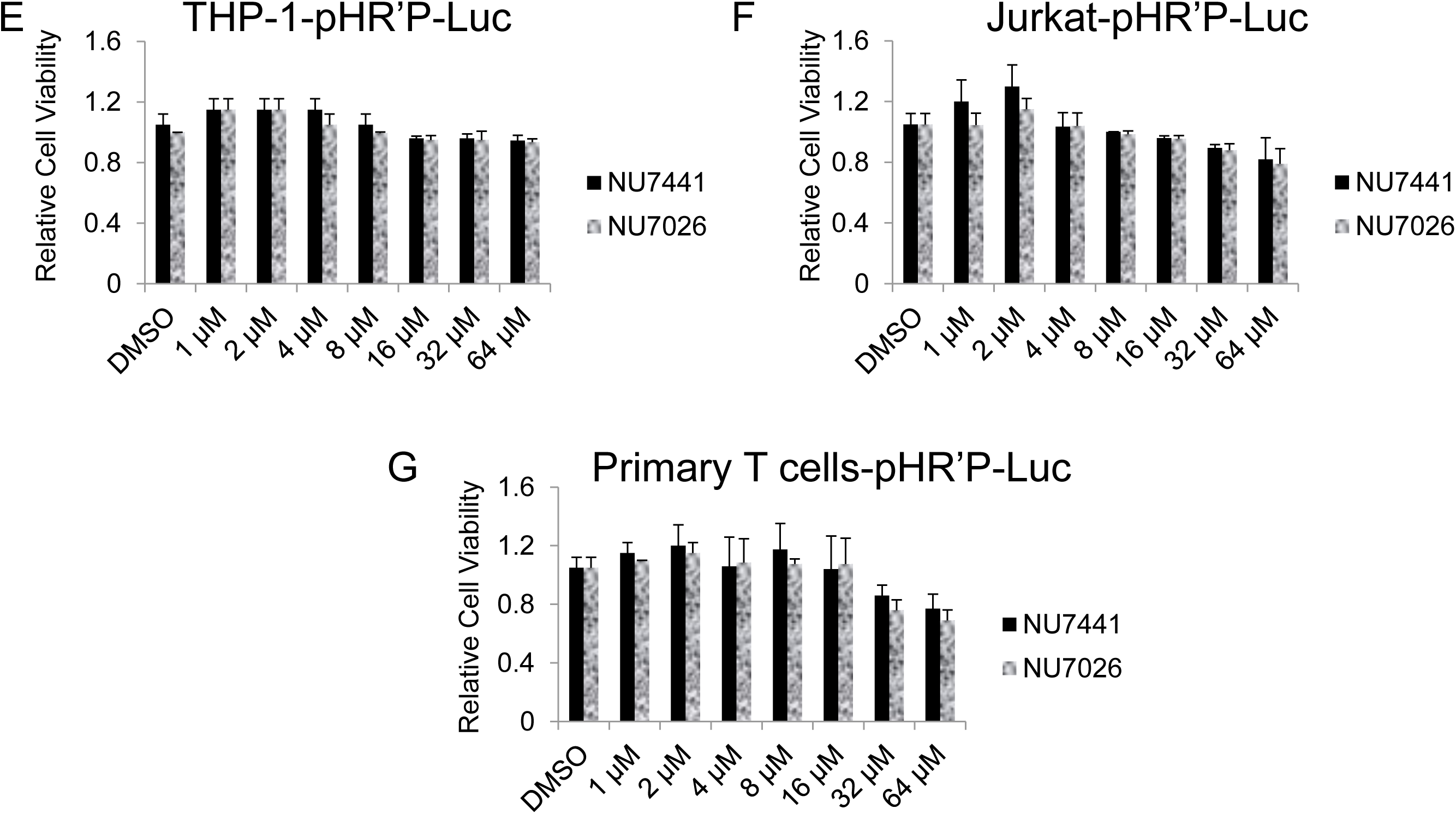
DNA-PK inhibitors repress HIV transcription without showing cytotoxicity and drastically impair latent proviral reactivation and HIV replication. MTS cell viability assays were performed after 5 days of culture in (**E**) THP-1-pHR’P-Luc, (**F**) Jurkat-pHR’fP-Luc and (**G**) primary CD4^+^ T cells-pHR’P-Luc. The results represent the Mean ± SD of three independent assays.

**Fig. 1:**
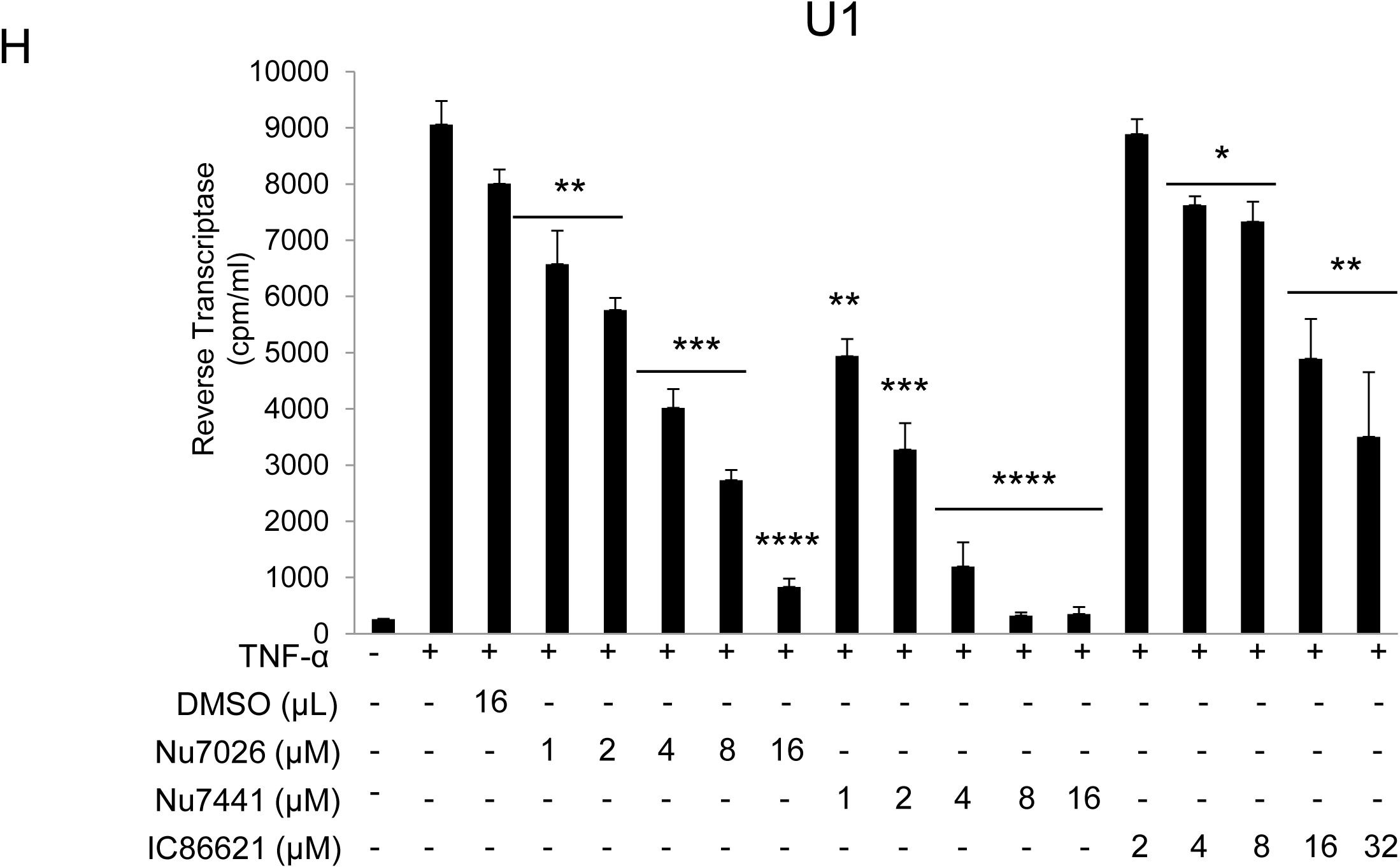
DNA-PK inhibitors repress HIV transcription without showing cytotoxicity and drastically impair latent proviral reactivation and HIV replication. For assessing the impact of DNA-PK inhibitors on the latent proviral reactivation and replication, the cells were pre-incubated with indicated amounts of DNA-PK inhibitors or DMSO control, overnight. Next day, cells were supplied with fresh media containing inhibitors and/or 10ng/ml TNF-α, accordingly, and further incubated for 48 hours. The cell supernatants were collected and used for the Reverse Transcriptase (RT) assays performed in (**H**) U1 (latently infected U937 based monocytic cell line).The results represent the Mean ± SD of three different independent assays. The p value of statistical significance was set at either; p < 0.05 (*), 0.01 (**), 0.001 (***), or 0.0001 (****).

**Fig. 1:**
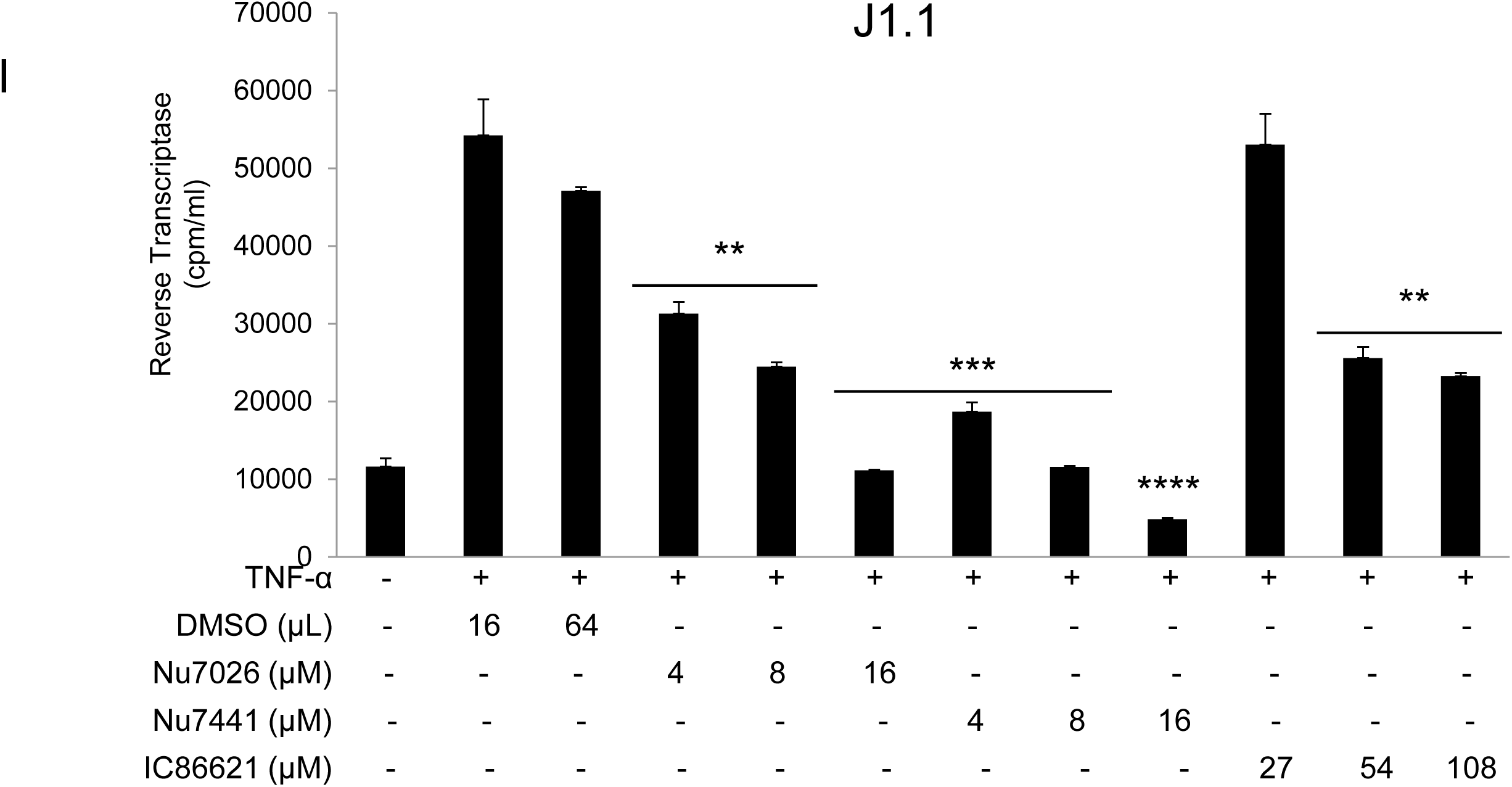
DNA-PK inhibitors repress HIV transcription without showing cytotoxicity and drastically impair latent proviral reactivation and HIV replication. (For assessing the impact of DNA-PK inhibitors on the latent proviral reactivation and replication, the cells were pre-incubated with indicated amounts of DNA-PK inhibitors or DMSO control, overnight. Next day, cells were supplied with fresh media containing inhibitors and/or 10ng/ml TNF-α, accordingly, and further incubated for 48 hours. The cell supernatants were collected and used for the Reverse Transcriptase (RT) assays performed in (**I**) J1.1 cell line (latently infected Jurkat based T cell line). The results represent the Mean ± SD of three different independent assays. The p value of statistical significance was set at either; p < 0.05 (*), 0.01 (**), 0.001 (***), or 0.0001 (****).

**Fig. 1:**
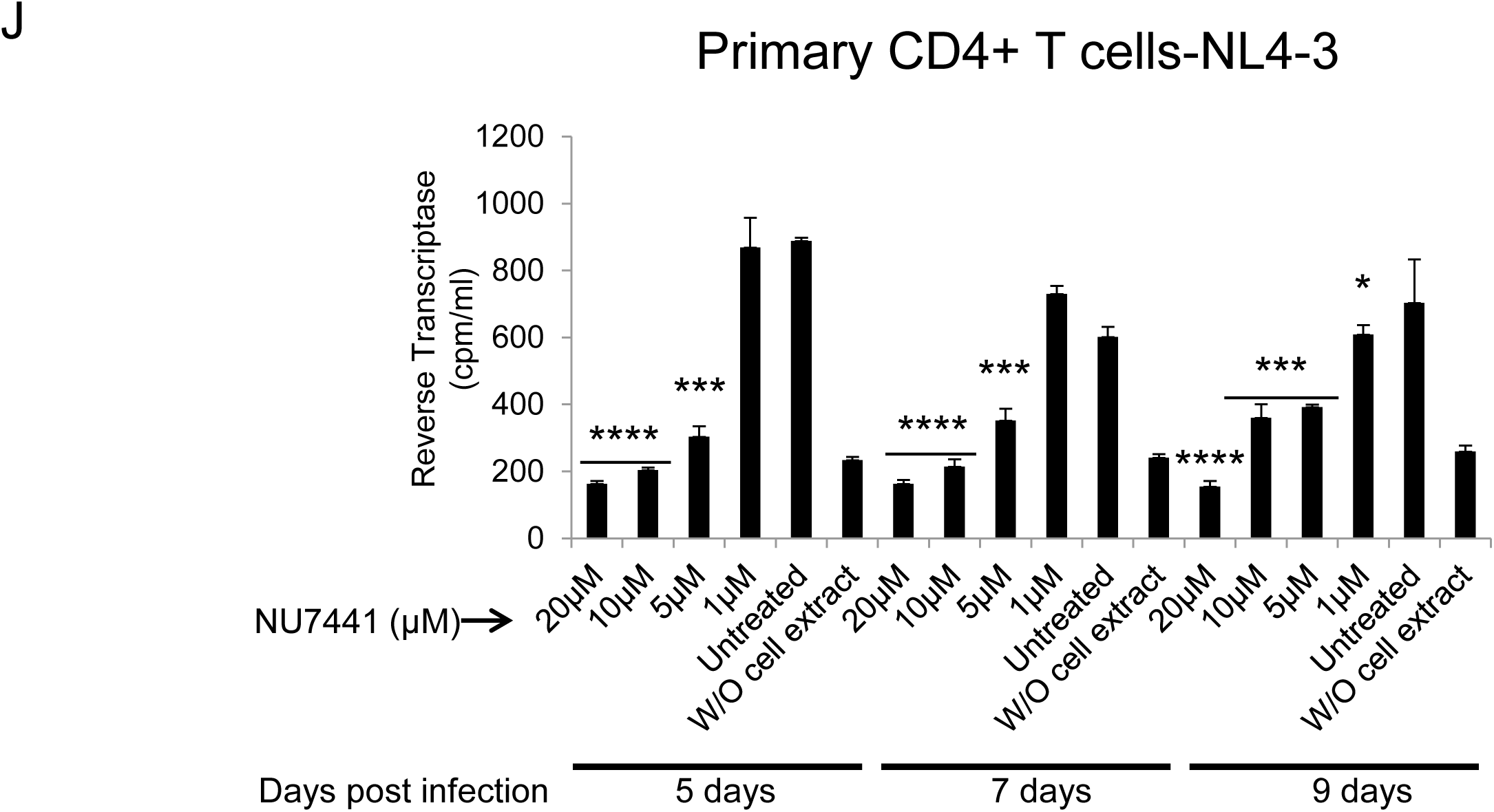
DNA-PK inhibitors repress HIV transcription without showing cytotoxicity and drastically impair latent proviral reactivation and HIV replication. For assessing the impact of DNA-PK inhibitors on the latent proviral reactivation and replication, the cells were pre-incubated with indicated amounts of DNA-PK inhibitors or DMSO control, overnight. Next day, cells were supplied with fresh media containing inhibitors and/or 10ng/ml TNF-α, accordingly, and further incubated for 48 hours. In (**J**), we assessed the impact of Nu7441 on HIV replication in freshly infected primary CD4^+^ T cells. The results represent the Mean ± SD of three different independent assays. The p value of statistical significance was set at either; p < 0.05 (*), 0.01 (**), 0.001 (***), or 0.0001 (****).

In order to exclude the possibility that the loss of luciferase activity was due to non-specific cell cytotoxicity of DNA-PK inhibitors, the cell viability was assessed using the MTS assay, after treating the cells with Nu7441 and Nu7026. We did not find any significant cell cytotoxicity in Jurkat and THP-1 cells up to 64 μM even after 5 days of culture (**Figs. 1E and F**). However, in case of primary CD4+ T cells, DNA-PK inhibitors showed some toxicity at concentrations around 32 μM (**Fig. 1G**). Therefore, in our successive analyses involving primary T cells, we used concentrations of inhibitors below 32 μM.

Taken together, our data suggest that DNA-PK inhibitors efficiently restrict proviral reactivation from latency by down-regulating HIV gene expression. The stronger suppression of HIV gene expression by the more specific DNA-PK inhibitors indicates target-specific inhibition and confirms the vital role of DNA-PK during HIV transcription.

We further used U1 (**Fig. 1H**) and J1.1 (**Fig. 1I**) cells to investigate the effect of DNA-PK inhibitors on latent proviral reactivation and HIV replication. The U1 is a latently infected monocytic cell line, while the J1.1 is a latently infected Jurkat T cell line; both carry an integrated replication-competent latent HIV provirus. The cells were treated with different amounts of DNA-PK inhibitors or DMSO control overnight (∼11 hours). Next day, cells were supplied with fresh media containing inhibitors and/or 10ng/ml TNF-α for another 48 hours, and the activity of HIV reverse transcriptase (RT) was analyzed in cell supernatants. We noticed a steady suppression of HIV replication by DNA-PK inhibitors in a dose-dependent manner (**Figs. 1H and I**). Similar to the results obtained in the transcription assays, the more specific inhibitors Nu7441 and Nu7026 were more effective than IC86621 in restricting HIV replication in both cell types. Comparable results from both J1.1 and U1 cells confirmed the vital role of DNA-PK in facilitating HIV gene expression and replication irrespective of cell lineage. These data also suggest the therapeutic potential of DNA-PK inhibitors in restricting the reactivation of latent HIV proviruses and subsequent replication.

For providing the physiological relevance to the anti-HIV effect of DNA-PK inhibitors, we examined the impact of DNA-PK inhibitors on HIV replication in primary CD4^+^ T cells, isolated from human peripheral blood mononuclear cells (PBMCs) (**Fig. 1J**). CD4^+^ T cells were activated with PHA and IL-2, infected with a CXCR4 tropic HIV virus (NL4-3), and treated with different concentrations of the most effective DNA-PK inhibitor, Nu7441. The culture medium containing Nu7441 was replaced every 3 days. The results of the RT assays performed at 5, 7 and 9 days post-infection showed that Nu7441 effectively repressed HIV replication in primary CD4^+^ T cells in a dose-dependent fashion. Taken together, our results confirm the pivotal role of DNA-PK during HIV transcription, latent proviral reactivation and replication.

### DNA-PK promotes HIV transcription by enhancing the phosphorylation of C-terminal domain (CTD) of RNA polymerase II (RNAP II)

In our prior publication, by performing *in vitro* kinase assays, we have shown the phosphorylation of RNAP II CTD by DNA-PK, a crucial post-translational modification, is required to generate complete functional transcripts, as it makes RNAP II processive (61). In order to understand if the observed inhibition of the HIV gene expression by DNA-PK inhibitors is the result of insufficient phosphorylation of RNAP II CTD, we assessed the impact of DNA-PK inhibitors on RNAP II CTD phosphorylation. Jurkat (**Fig. 2A**) and U937 (**Fig. 2C**), were treated with the DNA-PK inhibitor Nu7441 at different concentrations for 3 hours. The nuclear extracts were run on an acrylamide gel, transferred to a nitrocellulose membrane and the membrane was developed using antibodies that recognize specific epitopes present on the CTD of phosphorylated RNAP II. The H5 antibody recognizes the RNAP II with CTDs carrying Ser2 phosphorylation, while the H14 antibody recognizes epitopes of CTD carrying Ser5 phosphorylation. A dose-dependent effect of Nu7441 on the phosphorylation of Ser2 and Ser5 of RNAP II CTD in both Jurkat (**Figs. 2A and B**) and U937 cells (**Figs. 2C and D**) was observed. However, total RNAP II levels were not much affected. The parallel impairment of HIV gene expression (**Fig. 1**) and CTD phosphorylation (**Fig. 2**), upon DNA-PK inhibition, suggests an important role for DNA-PK-mediated CTD phosphorylation in generating processive RNAP II, a requirement for efficient gene expression. In addition, the present study validates our prior *in vitro* findings and confirms that RNAP II CTD is indeed the physiologic substrate for DNA-PK. The impact of DNA-PK inhibitors on both types of CTD phosphorylation modifications indicates that DNA-PK plays a role in both the initiation and elongation phases of transcription. Notably, the reduction in Ser5 phosphorylation is less pronounced than the reduction in Ser2 phosphorylation following inhibition of DNA-PK, indicating that DNA-PK facilitates HIV transcription predominantly by supporting the elongation phase.

**Fig. 2:**
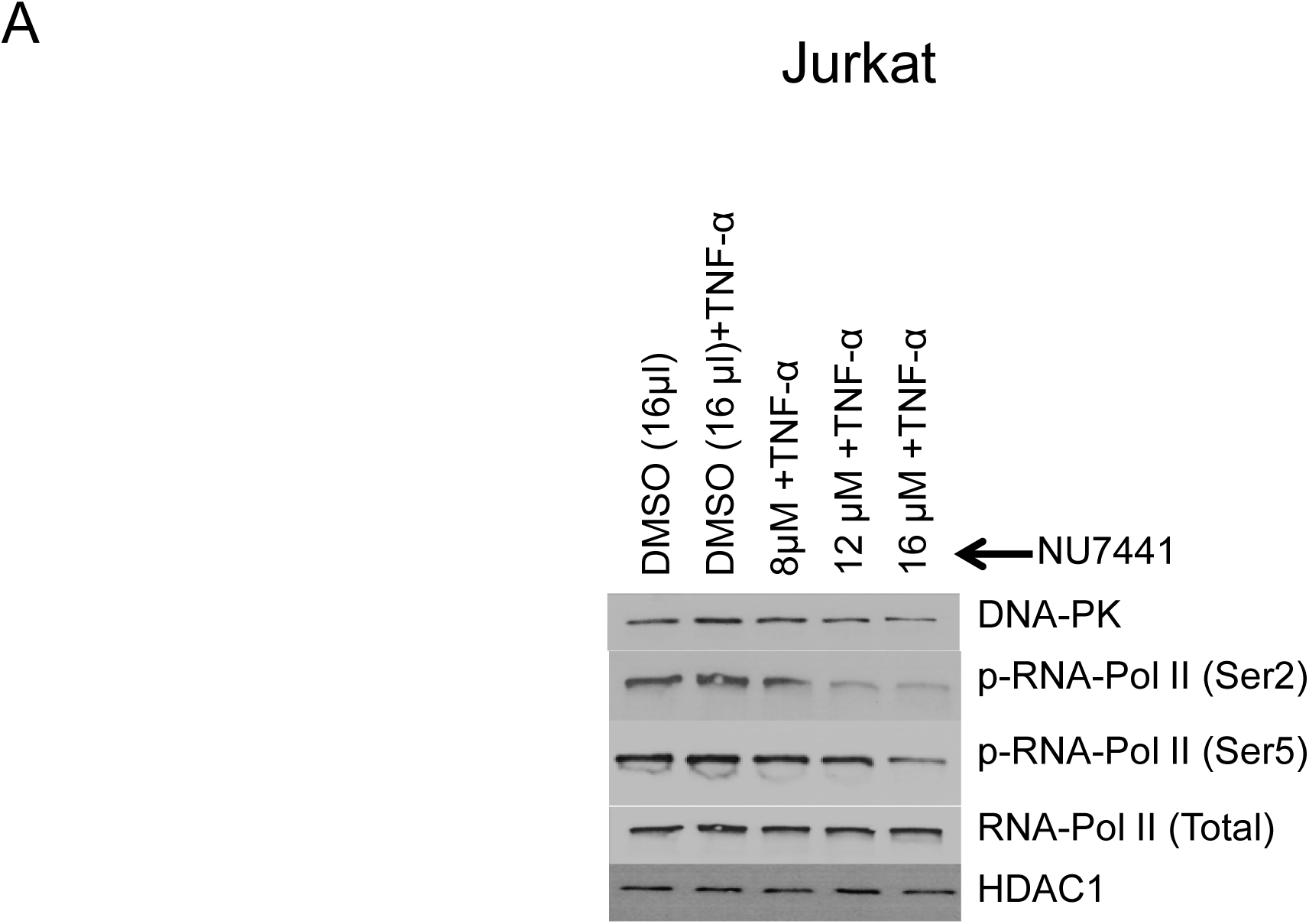
DNA-PK inhibitors repress HIV transcription by restricting RNAP II CTD phosphorylation. (**A**) Jurkat cells were treated with increasing concentrations of the DNA-PK inhibitor Nu7441 and then activated with TNF-α (10 ng/ml) for 3 hours. The cells were harvested and nuclear extracts were assessed through western blot. The phosphorylation state of carboxyl terminal domain (CTD) of RNA polymerase II (RNAP II) was assessed through western blot using antibodies that are specific for RNAP II and its phosphorylated forms (Ser2 and Ser5). HDAC1 was used as loading control. The presented data is one representative western blot analysis out of three.

**Fig. 2:**
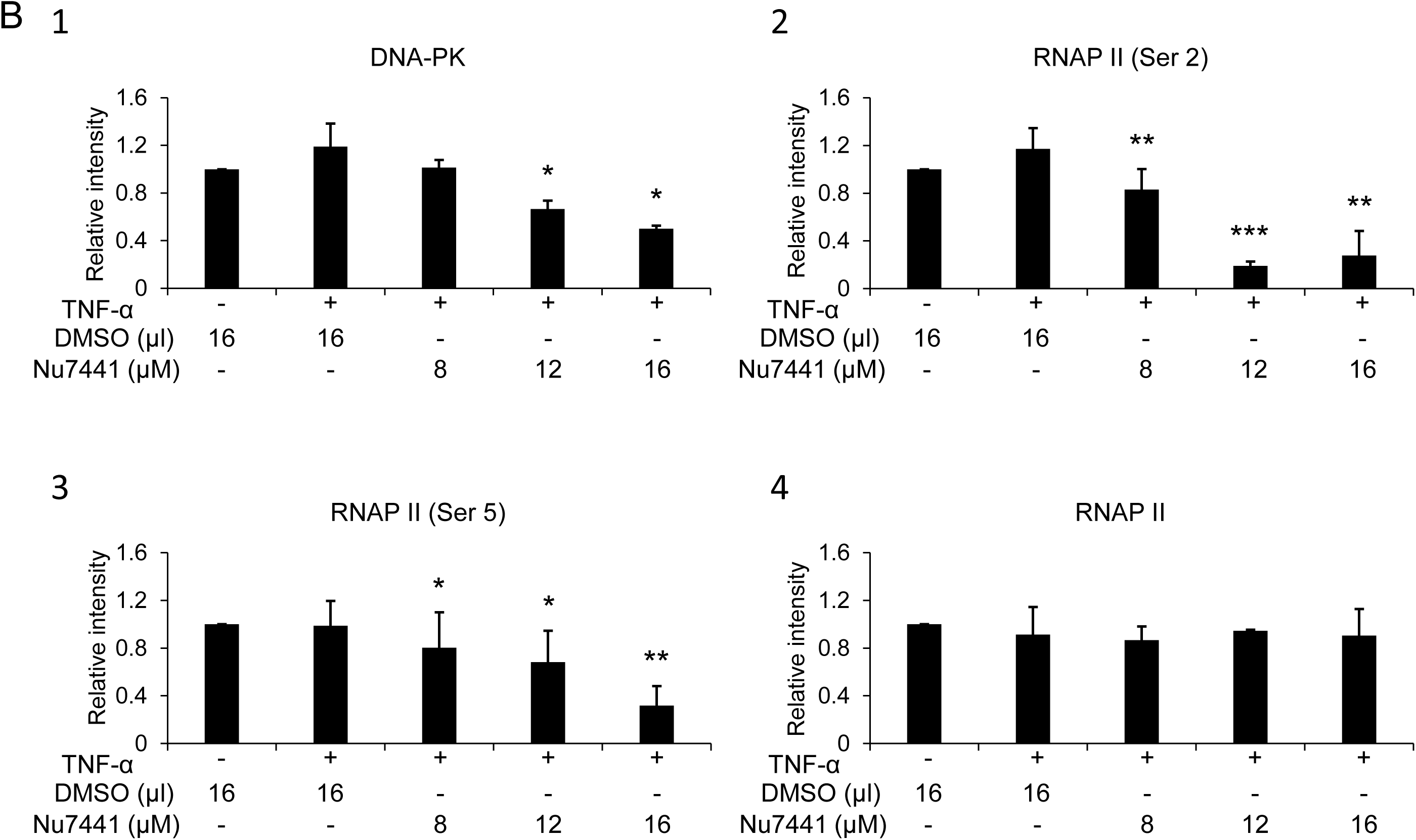
DNA-PK inhibitors repress HIV transcription by restricting RNAP II CTD phosphorylation. Densitometric analyses were performed on western blot bands using ImageJ software for (**B**) Jurkat cells gels. The results represent the Mean ± SD of three different independent assays. Statistical significance is set as p < 0.05 (*), 0.01 (**) or 0.001 (***).

**Fig. 2:**
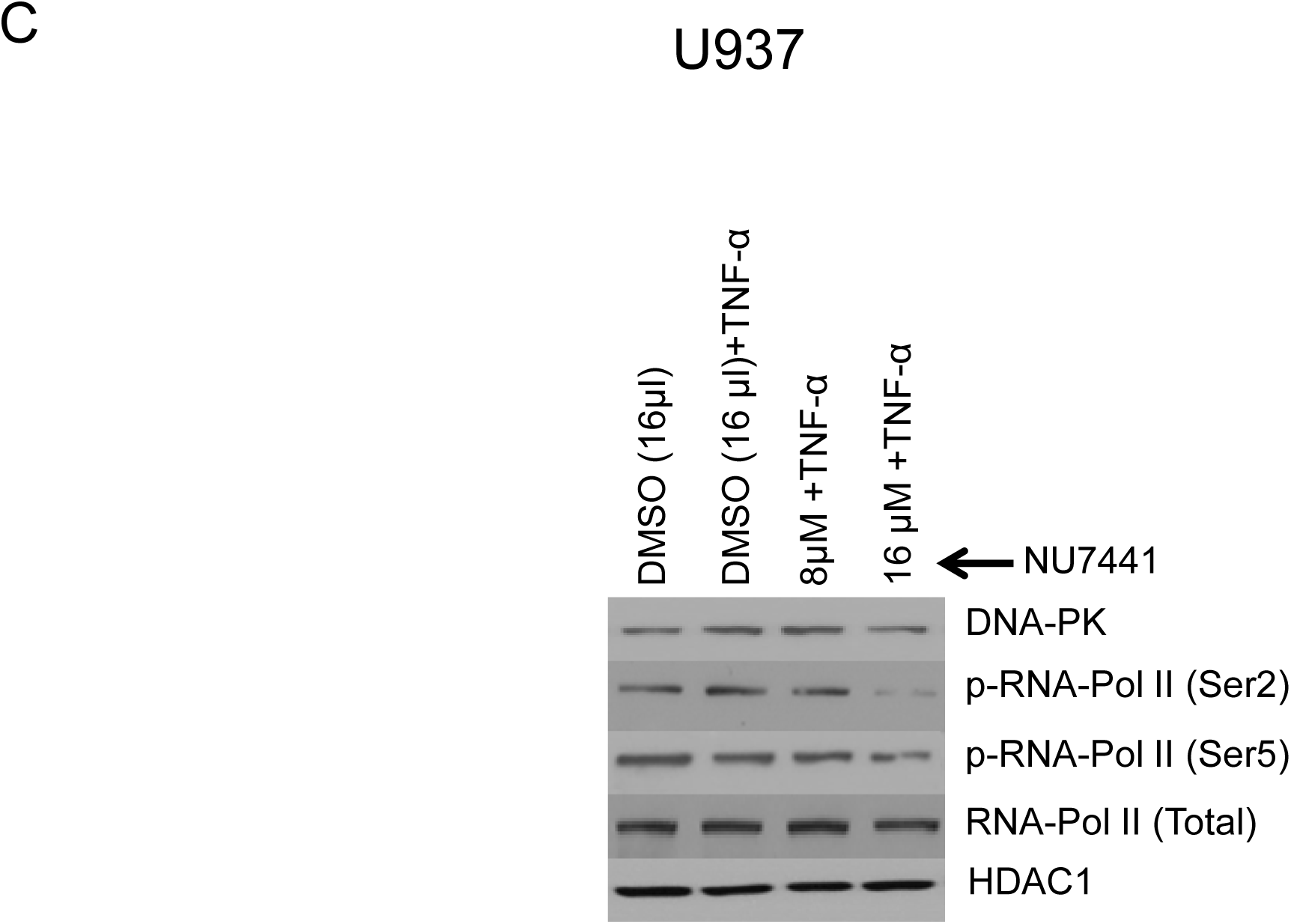
DNA-PK inhibitors repress HIV transcription by restricting RNAP II CTD phosphorylation. (**C**) U937 cells were treated with increasing concentrations of the DNA-PK inhibitor Nu7441 and then activated with TNF-α (10 ng/ml) for 3 hours. The cells were harvested and nuclear extracts were assessed through western blot. The phosphorylation state of carboxyl terminal domain (CTD) of RNA polymerase II (RNAP II) was assessed through western blot using antibodies that are specific for RNAP II and its phosphorylated forms (Ser2 and Ser5). HDAC1 was used as loading control. The presented data is one representative western blot analysis out of three.

**Fig. 2:**
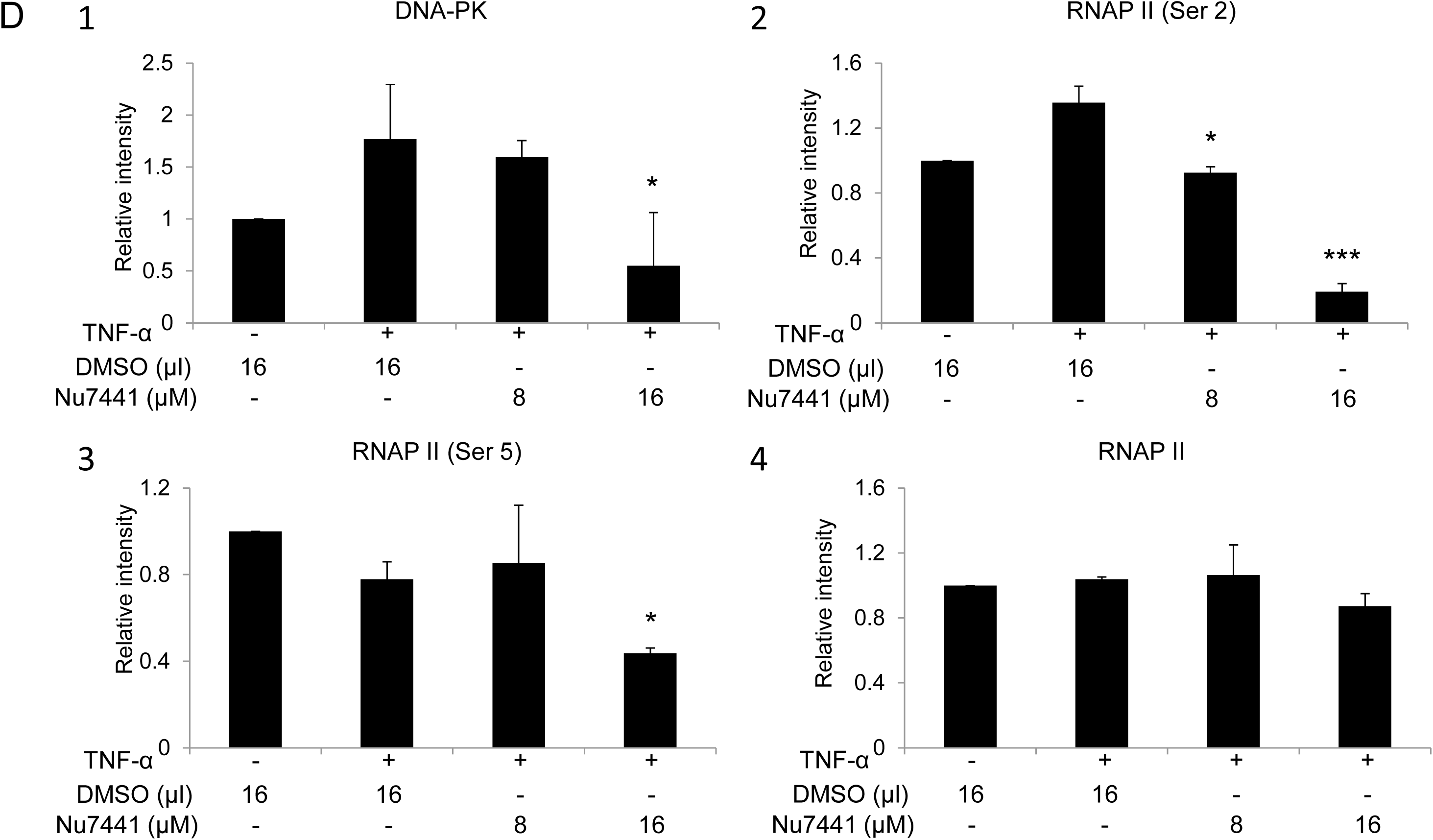
DNA-PK inhibitors repress HIV transcription by restricting RNAP II CTD phosphorylation. Densitometric analyses were performed on western blot bands using ImageJ software for (**D**) U937 cells gels. The results represent the Mean ± SD of three different independent assays. Statistical significance is set as p < 0.05 (*), 0.01 (**) or 0.001 (***).

Overall, these results confirm that DNA-PK catalyzes the phosphorylation of RNAP II CTD, and suggest that the observed reduction in HIV transcription and replication following treatment with DNA-PK inhibitors is a direct consequence of reduced CTD phosphorylation of RNAP II.

### Knockdown of endogenous DNA-PK restricts RNAP II CTD phosphorylation

In order to confirm that the effects we observed after DNA-PK inhibition are certainly due to the impairment of DNA-PK activity, the knockdown of endogenous DNA-PK was performed. To knockdown endogenous DNA-PK, the U937 and Jurkat cells were infected with lentiviral vectors expressing shRNAs either against the catalytic subunit of DNA-PK (DNA-PKc) or control scrambled shRNA. The neutrality of scrambled shRNA towards HIV and cellular genomes was confirmed by using the NCBI program Blast (59). Depending on the degree of DNA-PK knockdown in different clones, we selected one clone each for U937 (clone U3) (**Figs. 3A and B**), and Jurkat (clone J3) (**Figs. 3C and D**) for further analysis. In case of U937 clone U3, we were able to obtain more than 85% knockdown (**Fig. 3A**), whereas in case of Jurkat clone J3 the knockdown was 70% (**Fig. 3C**). We also noted profound cell death in Jurkat cells with over 70% depletion of endogenous DNA-PK. In both knockdown clones, we noted loss of the CTD phosphorylation, at both Ser2 and Ser5. We also observed some reduction of the total RNAP II levels in U937 cells, likely due to high level of DNA-PK depletion. Given the fact that RNAP II is also required for RNAP II gene expression, severe defects to the functionality of cellular RNAP II hamper its own expression, an effect also observed upon using strong inhibitors of P-TEFb (data not shown).

**Fig. 3:**
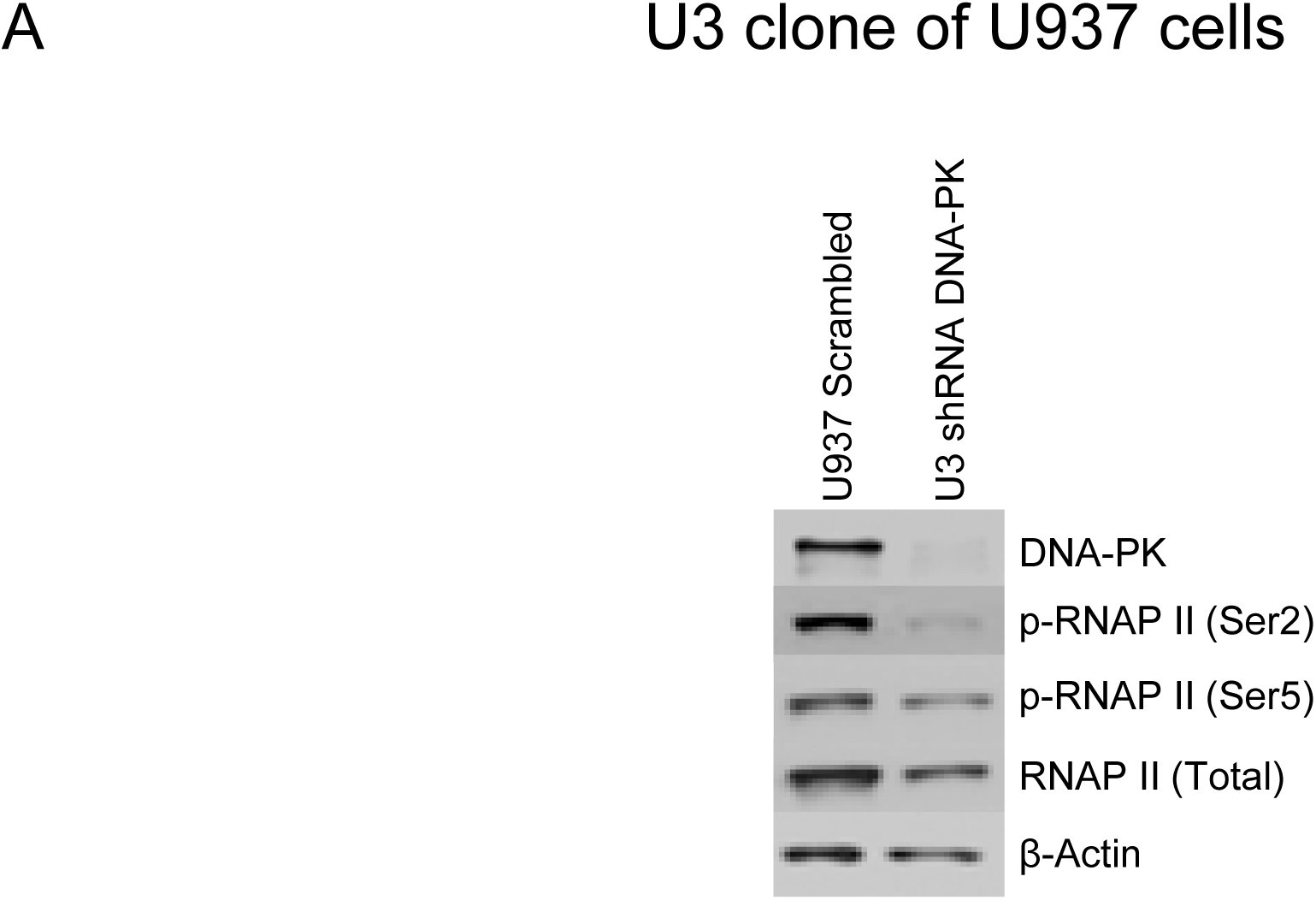
Knockdown of DNA-PK restricts RNAP II CTD phosphorylation. The endogenous DNA-PK was knocked down using lentiviral vectors expressing shRNA sequences specific for DNA-PK. The vector expressing non-targeting scrambled shRNA sequence was used as control. Western blot assays showing nuclear levels of DNA-PK, total RNAP II, RNAP II (Ser2) and RNAP II (Ser5) in (**A**) U937 cells and corresponding DNA-PK knockdown clone U3.

**Fig. 3:**
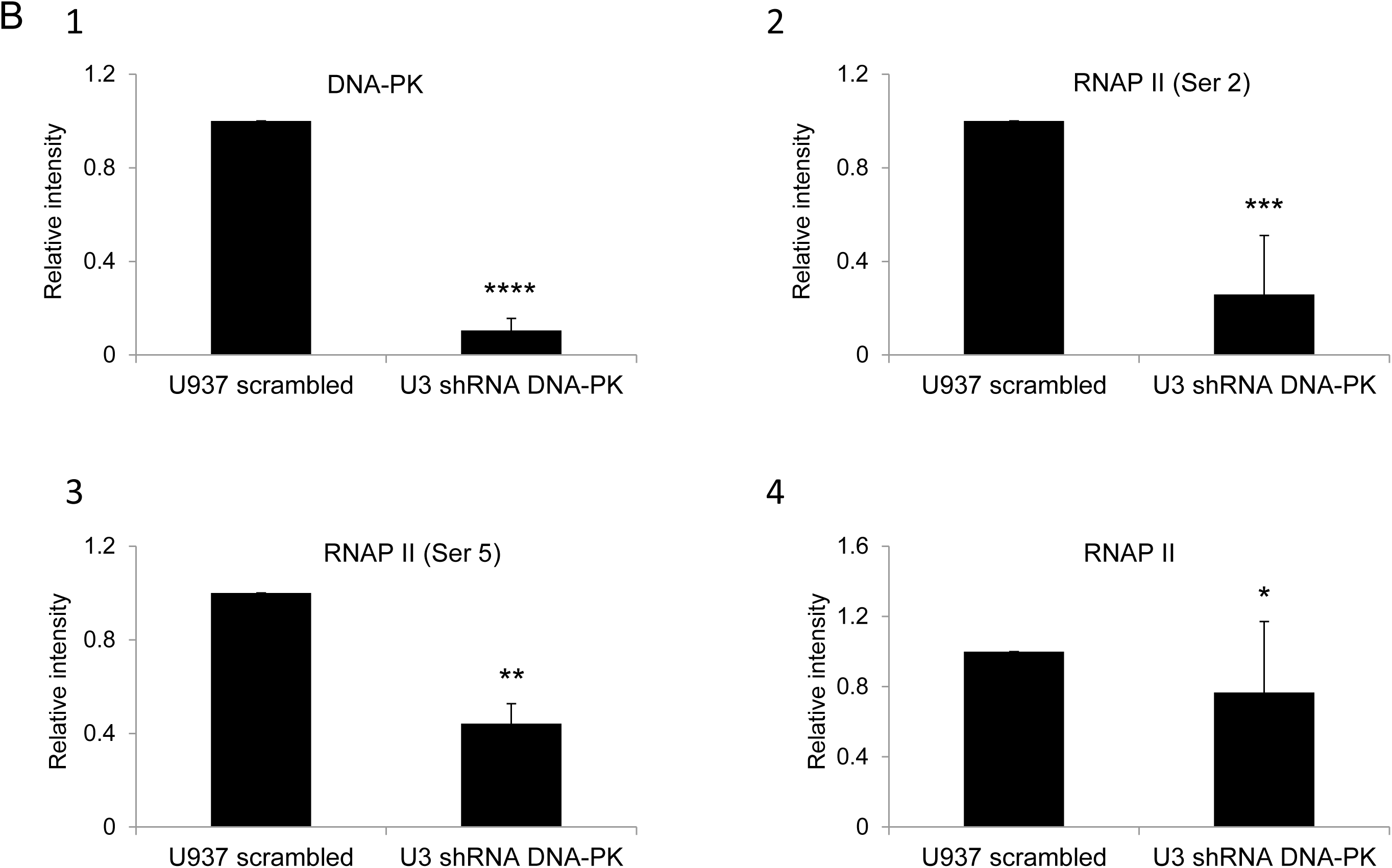
Knockdown of DNA-PK restricts RNAP II CTD phosphorylation. Densitometric analyses were performed on (**B**) U937 cells western blot’s bands. Error bars represent the SD of three separate experiments. The statistical significance was set at either; p < 0.05 (*), 0.01 (**), 0.001 (***), or 0.0001 (****).

**Fig. 3:**
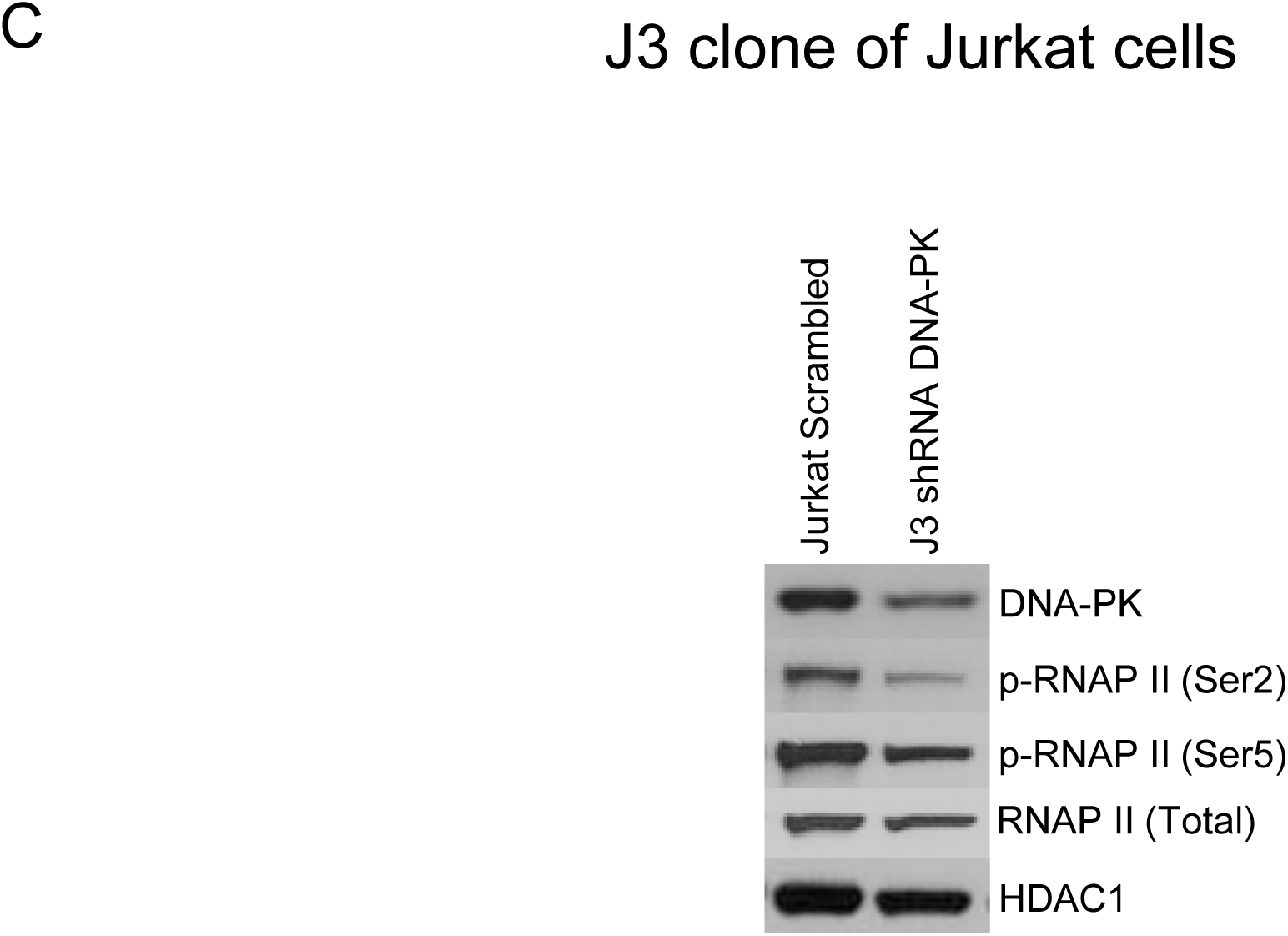
Knockdown of DNA-PK restricts RNAP II CTD phosphorylation. The endogenous DNA-PK was knocked down using lentiviral vectors expressing shRNA sequences specific for DNA-PK. The vector expressing non-targeting scrambled shRNA sequence was used as control. Western blot assays showing nuclear levels of DNA-PK, total RNAP II, RNAP II (Ser2) and RNAP II (Ser5) in Jurkat cells and DNA-PK knockdown clone J3. β-Actin and HDAC1 were used as loading controls.

**Fig. 3:**
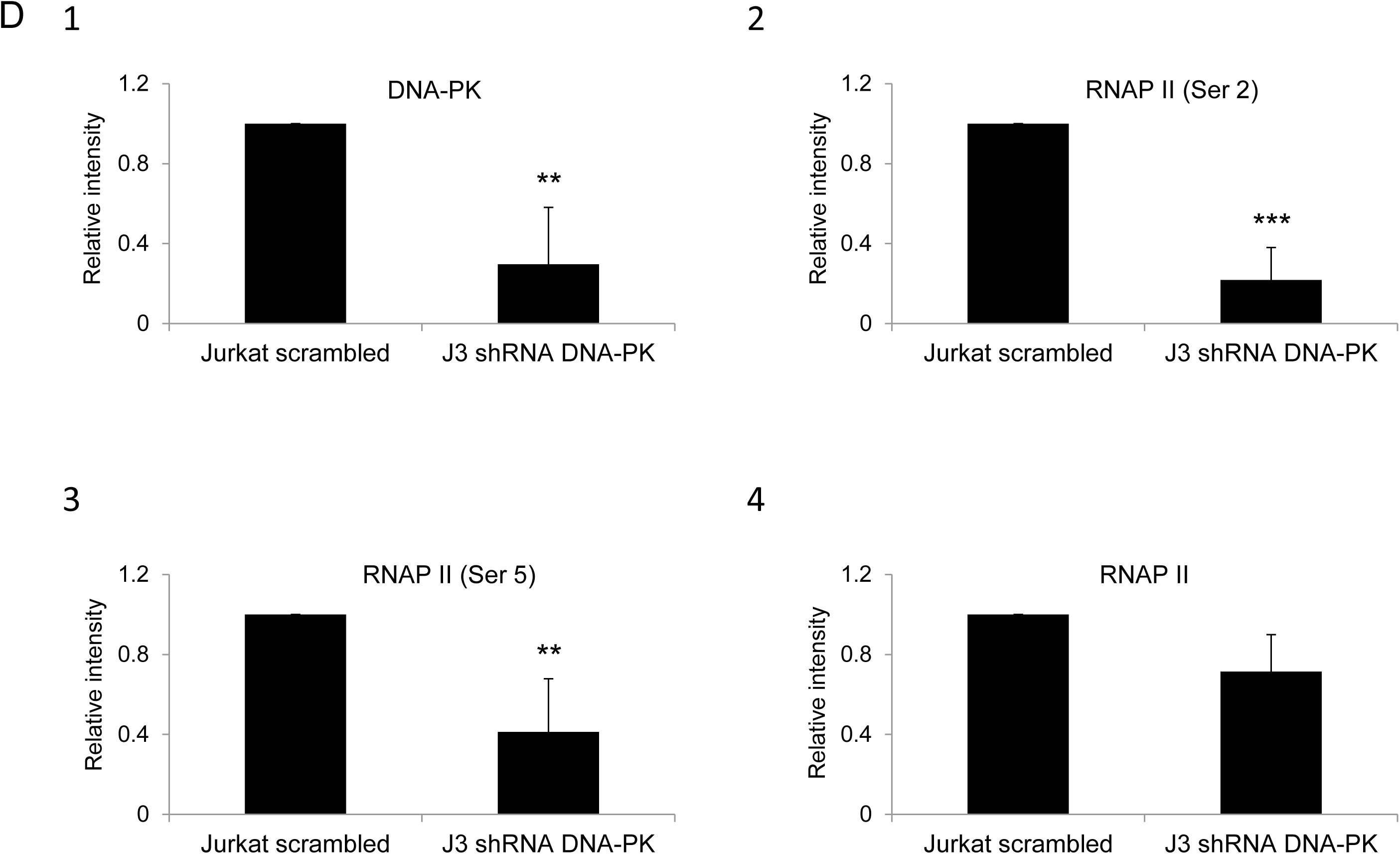
Knockdown of DNA-PK restricts RNAP II CTD phosphorylation. Densitometric analyses were performed on (**D**) Jurkat cells western blot’s bands. Error bars represent the SD of three separate experiments. Statistical significance is set as p < 0.05 (*), 0.01 (**) or 0.001 (***).

These results, along with those obtained using DNA-PK inhibitors (**Fig. 2**), confirm that DNA-PK catalyzes RNAP II CTD phosphorylation, which promotes HIV transcription.

### DNA-PK promotes HIV transcription by supporting the recruitment of P-TEFb at HIV LTR

The interaction between DNA-PK and P-TEFb has been previously documented (1, 36). CDK9 subunit of P-TEFb is the major kinase that catalyzes the phosphorylation of RNAP II CTD. This observation suggests that in addition to directly catalyzing the phosphorylation of CTD of RNAP II, DNA-PK may enhance CTD phosphorylation by increasing the recruitment of P-TEFb at HIV LTR. Therefore, we examined the recruitment profile of the main subunits of P-TEFb, Cyclin T1 and CDK9, at HIV LTR by performing chromatin immunoprecipitation (ChIP) assays in wild type and DNA-PK knockdown cells. Given that T cells are the main reservoirs of latent proviruses, we performed experiments using Jurkat T cells. The recruitment kinetics of P-TEFb were analyzed in knockdown (J3) and control Jurkat clones harboring latent HIV provirus, pHR’-PNL-wildTat-d2GFP (Jurkat-HIV-GFP) (**Fig. 4A**), before and after activation with TNF-α (10 ng/ml) for 30 min.

**Fig. 4:**
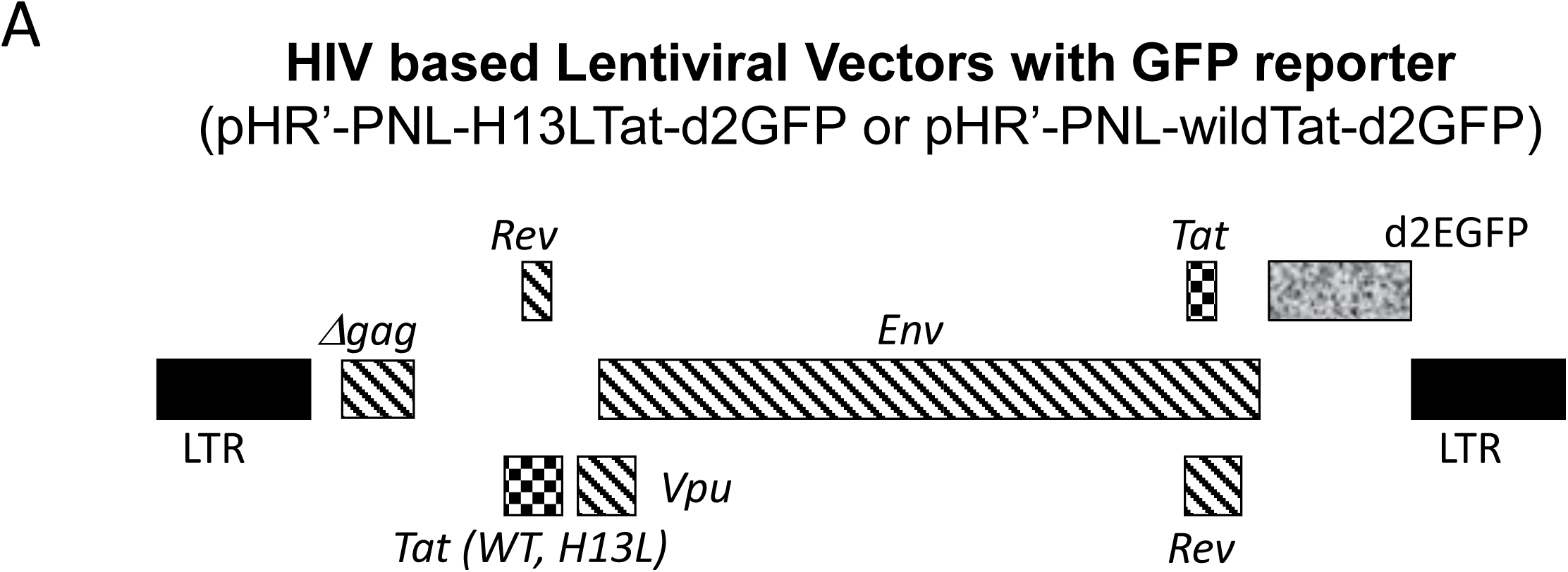
DNA-PK facilitates the recruitment of P-TEFb at HIV LTR. (**A**) Schematic diagram of HIV based lentiviral vector, which carry either mutated Tat (H13L) or wild type Tat and express GFP as reporter.

**Fig. 4:**
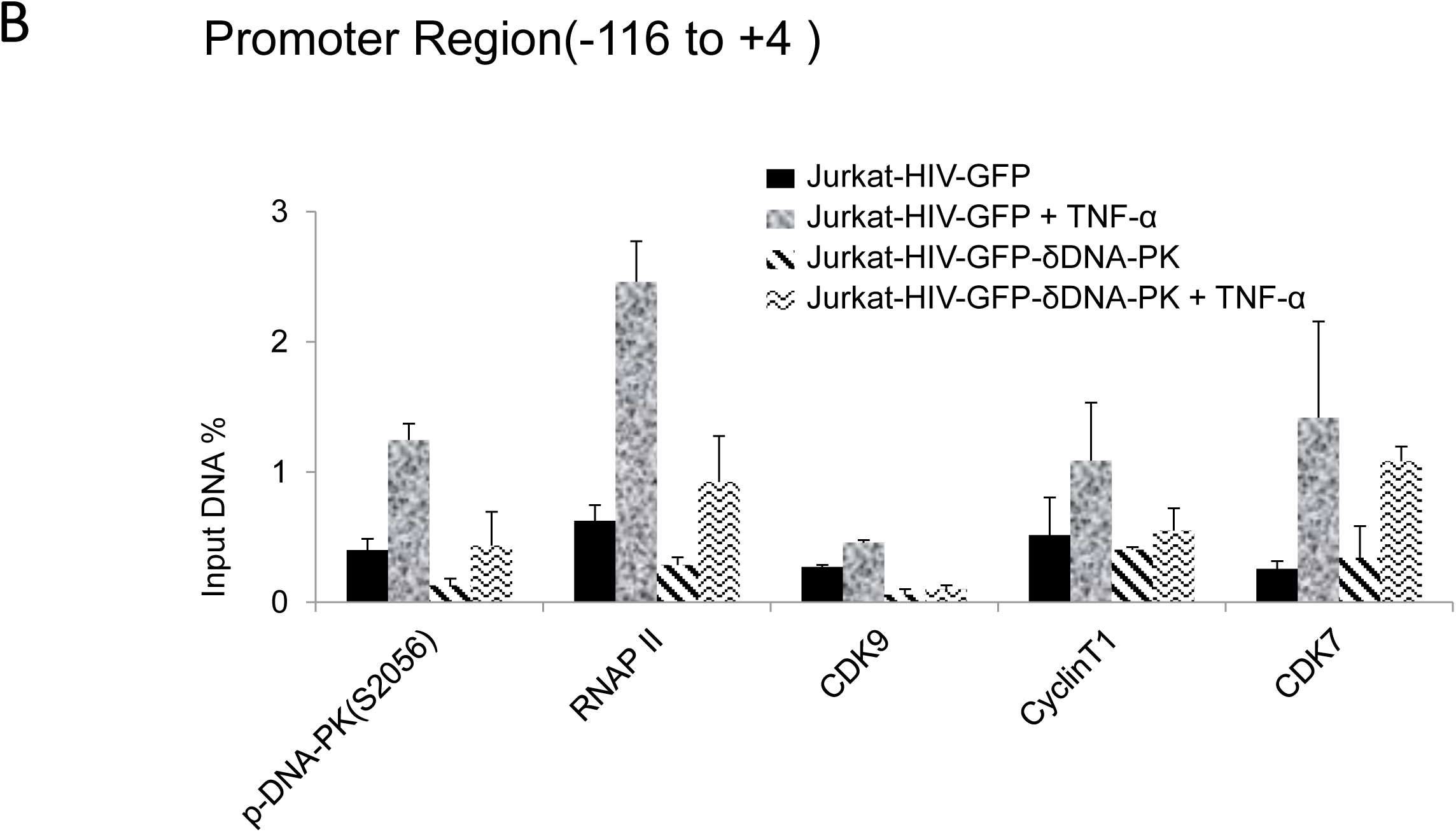
DNA-PK facilitates the recruitment of P-TEFb at HIV LTR. In these experiments VSV-G pseudotyped HIV, harboring wild type Tat gene (HIV-GFP), was used. The DNA-PK knockdown (J3) or control Jurkat cells with scrambled shRNA were infected with HIV-GFP and the provirus was allowed to become latent. The ChIP assays were performed before and after reactivating the latent provirus with TNF-α (10 ng/ml) for 30 minutes. ChIP assays were done using indicated antibodies and two transcriptionally relevant regions of LTR were amplified using specific primer sets: (**B**) promoter region of the LTR. Data are presented as percentage of input. Error bars represent the Mean ± SD of two independent experiments and three separate real-time qPCR measurements.

**Fig. 4:**
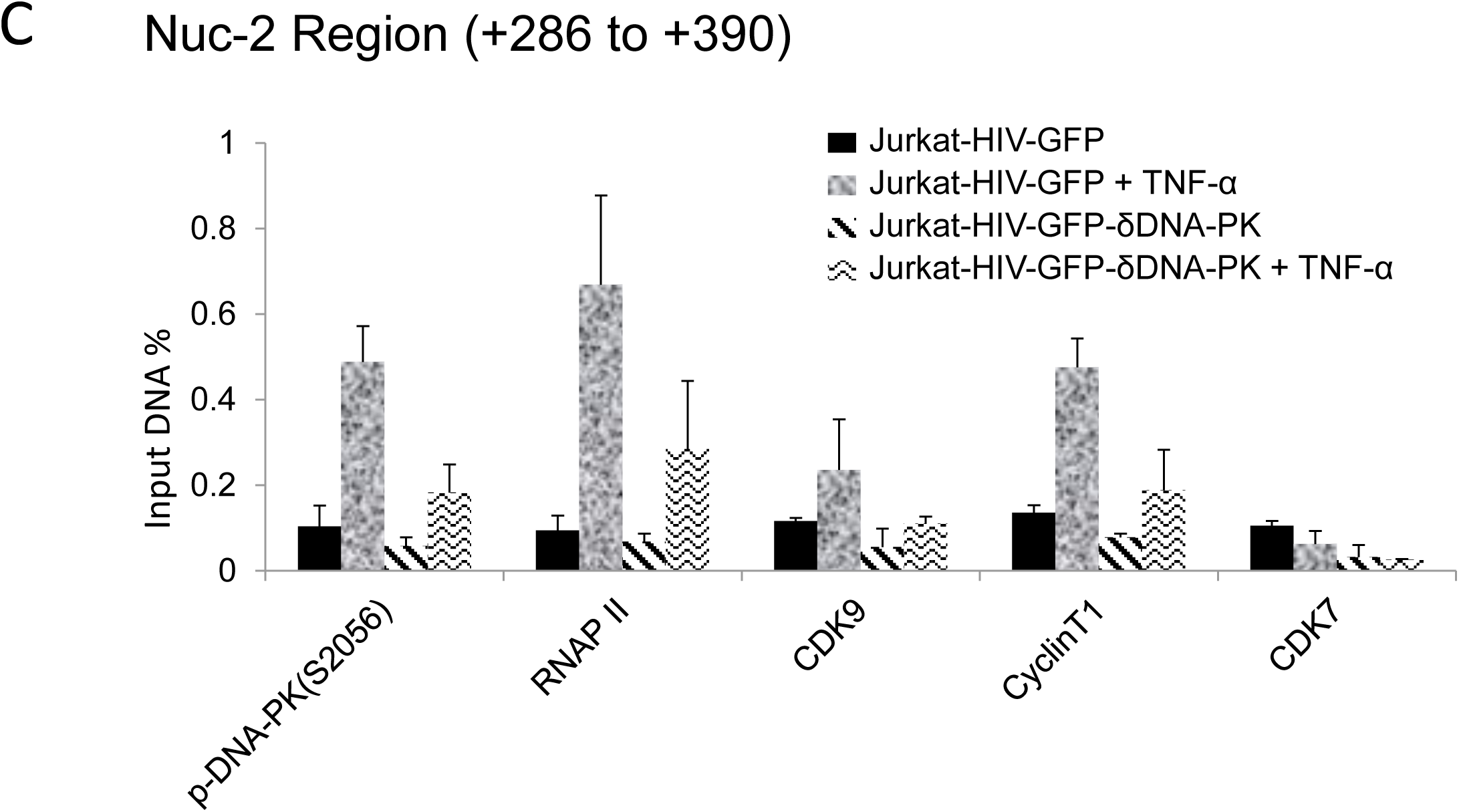
DNA-PK facilitates the recruitment of P-TEFb at HIV LTR. In these experiments VSV-G pseudotyped HIV, harboring wild type *Tat* gene (HIV-GFP), was used. The DNA-PK knockdown (J3) or control Jurkat cells with scrambled shRNA were infected with HIV-GFP and the provirus was allowed to become latent. The ChIP assays were performed before and after reactivating the latent provirus with TNF-α (10 ng/ml) for 30 minutes. ChIP assays were done using indicated antibodies and two transcriptionally relevant regions of LTR were amplified using specific primer sets: (**C**) Nuc-2 region of the LTR. HIV DNA levels were calculated as percentages of the reaction input. Error bars represent the Mean ± SD of two independent experiments and three separate real-time qPCR measurements.

The ChIP assays were performed using antibodies against Cyclin T1 and CDK9. We also analyzed CDK7, a component of the TFIIH transcription factor, which catalyzes the phosphorylation of RNAP II CTD primarily at Ser5 residue, hence playing vital role during the initiation phase of HIV transcription (29). As control, we assessed the recruitment of RNAP II, a marker of ongoing transcription, and an auto-phosphorylated form of DNA-PK, specifically at S2056, p-DNA-PK (S2056), which marks the functionally active form of DNA-PK. The immunoprecipitated DNA was examined by using two primer sets, one amplifying the promoter region of the LTR (-116 to +4 with respect to the transcription start site), which provides a measure of the factors involved in transcriptional initiation (**Fig. 4B**), the other one amplifying the downstream nucleosome-2 region (Nuc-2) of the LTR (+286 to +390 with respect to the transcription start site), which quantifies the factors involved in the elongation phase of HIV transcription (**Fig. 4C**). The IgG control was subtracted from all samples as background. The results were normalized with the housekeeping GAPDH gene (-145 to +21), a constitutively expressing cellular gene. In figure 4B, the low level of RNAP II at the promoter region of latent provirus marks reduced ongoing transcription. Interestingly, we also noted highly reduced presence of p-DNA-PK(S2056), suggesting that due to reduced level of DNA-PK in latently infected cells, HIV provirus was unable to initiate its transcription. However, upon induction with TNF-α, the latent provirus gets reactivated, as indicated by the rapid enrichment (more than three folds) of RNAP II at the promoter. Parallel to the RNAP II accumulation, we found enhanced recruitment of DNA-PK at the promoter, validating the pivotal role of DNA-PK in HIV transcription. The enhanced nuclear-levels of DNA-PK upon cell stimulation were also validated in other experiments (**Figs. 2 and 7**). As anticipated, we noted relatively less recruitment of P-TEFb subunits at the promoter region. However, at a distal Nuc-2 region of LTR, we noticed significantly higher enrichment of P-TEFb upon activation, validating main role of P-TEFb during the elongation phase of transcription. On the other hand, after activation, CDK7 was enriched as expected at LTR promoter, but not at Nuc-2 region, again validating its requirement especially during the initiation phase of HIV transcription. As shown in our previous publication (61), we noted better enrichment of DNA-PK at the downstream Nuc-2 region of LTR after activation, validating the main role of DNA-PK during transcriptional elongation. However, in DNA-PK deficient cells, we observed highly reduced recruitment of almost all the factors, further validating the highly restricted ongoing HIV transcription. Interestingly, even after stimulation with TNF-α, we found highly impaired enrichment of P-TEFb at LTR, confirming a clear role of DNA-PK in P-TEFb recruitment. Notably, we observed a relatively smaller impact of DNA-PK knockdown on CDK7 recruitment at promoter region, suggesting that CDK7 recruitment at LTR does not depend on the DNA-PK.

Together, our results demonstrated that DNA-PK promotes HIV transcription, not only via directly catalyzing the phosphorylation of CTD of RNAP II (61), but also by enhancing the recruitment of P-TEFb at HIV LTR, which further increases CTD phosphorylation.

### DNA-PK depletion in cells severely impairs HIV gene expression

To confirm the direct involvement of DNA-PK in promoting HIV transcription, we assessed the effect of DNA-PK knockdown on HIV gene expression. We superinfected DNA-PK knockdown cells or control cells that carry scrambled shRNA, with pHR’-PNL-H13LTat-d2GFP or pHR’-PNL-wildTat-d2GFP. These non-replicating HIV have similar backbones and both express GFP reporter gene under the control of HIV LTR promoter (**Fig. 4A**). The only difference between them is that one expresses wild-type Tat and the other mutated Tat (H13L), a partially defective Tat mutant highly prevalent in latent proviruses (43, 59, 70). Interestingly, for both Jurkat and U937 cells, upon superinfection with pHR’-PNL-H13LTat-d2GFP in DNA-PK depleted cells, we noted more than 4-fold reduction of GFP positive cells in comparison to control cells carrying scrambled shRNA (**Figs. 5A and B**). However, in cells infected with HIV expressing wild-type *Tat* gene, pHR’-PNL-wildTat-d2GFP, we observed relatively less impaired HIV gene expression, but still more than 2-fold reduction of GFP positive cells in DNA-PK depleted cells. Given the fact that the expression of *GFP* gene is controlled by the HIV LTR promoter in these HIV constructs, the GFP expression marks HIV gene expression and LTR activation. These results demonstrate that DNA-PK plays crucial role in promoting HIV gene expression. Accordingly, similar to GFP expression, we noticed reduced levels of Tat in DNA-PK knockdown U2 and U3 clones infected with pHR’-PNL-wildTat-d2GFP (**Fig. 5C**), suggesting that DNA-PK-mediated transcriptional boost is essential for generating optimal Tat levels as well.

**Fig. 5:**
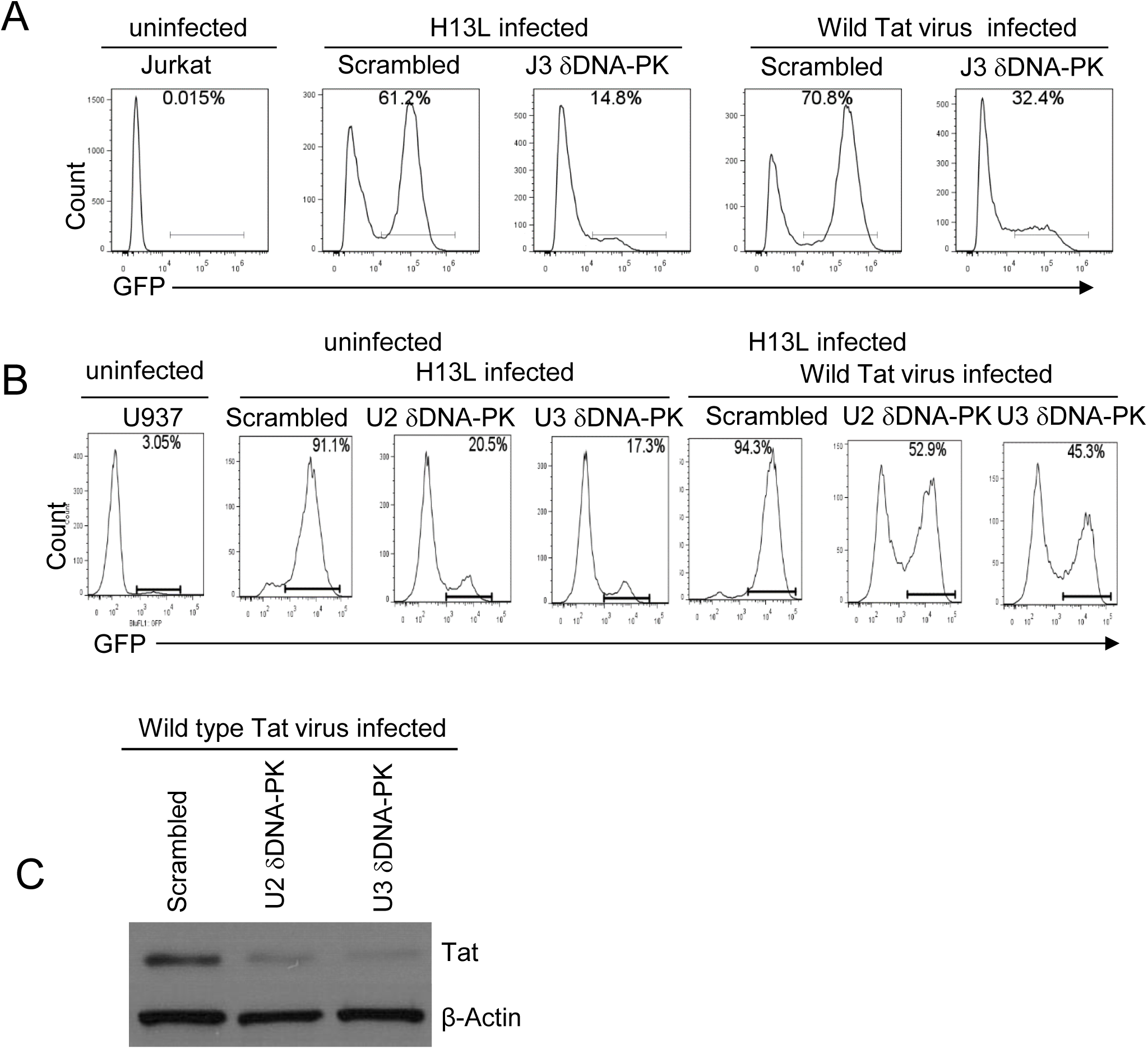
DNA-PK plays a vital role in HIV gene expression. Flow cytometric analyses of freshly-infected DNA-PK knockdown or control cells with either pHR’-PNL-H13LTat-d2GFP or pHR’-PNL-wildTat-d2GFP. (**A**) DNA-PK knockdown Jurkat clone J3 and control Jurkat with scrambled shRNA.(**B**) DNA-PK knockdown U937 cell clones, U2 and U3, and control cells with scrambled shRNA. After 48 hours post-infection, GFP positive cells were analyzed using a flow cytometer. The uninfected Jurkat and U937 were used as background control. (**C**) Tat protein levels in DNA-PK knockdown U2 and U3 clones infected with pHR’-PNL-wildTat-d2GFP analyzed via western blot.

**Fig. 5:**
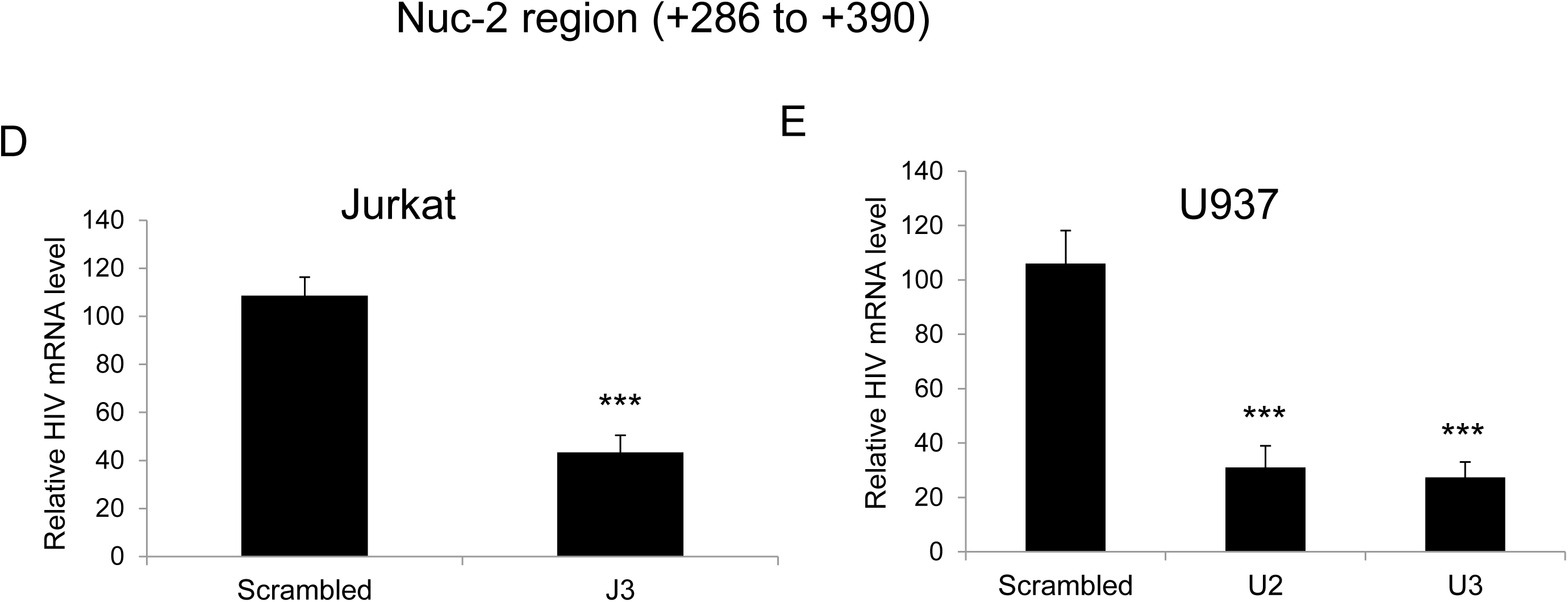
DNA-PK plays a vital role in HIV gene expression. (**D** and **E**), The relative expression levels of HIV mRNA in knockdown and control cells evaluated via RT-qPCR. The qPCR results represent the Mean ± SD of three independent assays. Statistical significance is set as p < 0.05 (*), 0.01 (**) or 0.001 (***).

The results were also confirmed at the mRNA level. The cellular RNAs were extracted from control and DNA-PK knockdown cell clones infected with pHR’-PNL-wildTat-d2GFP; cDNAs were synthesized and equal amount of cDNAs were quantified via real-time PCR analysis (qPCR) using primer set for the downstream Nuc-2 region of HIV LTR. Analogously to GFP levels, we observed a significantly reduced expression of HIV mRNA in DNA-PK knockdown clones compared to cells that express normal levels of DNA-PK (**Figs. 5D and E**). These data confirm that GFP expression is impaired at transcriptional level in DNA-PK deficient cells.

### DNA-PK enhances HIV transcription by facilitating the recruitment of transcriptional machinery and generating transcriptionally active euchromatin structures at HIV LTR

It is well established that the type of chromatin structure and transcription factors present at the gene promoter and nearby regions, regulate its transcription (60, 63). We examined the recruitment of the main transcription factors and induced chromatin structures at and around the HIV LTR promoter. Heterochromatin or closed chromatin structures repress transcription, whereas, euchromatin or open chromatin structures facilitate transcription by allowing access to the transcription machinery at the promoter (27, 53, 58). Hence, for characterizing the underlying molecular mechanisms through which DNA-PK promotes HIV gene expression and replication, we evaluated the changes to chromatin structures at HIV LTR before and after inhibiting DNA-PK.

To examine whether DNA-PK inhibitors instigate the generation of transcriptionally repressive heterochromatin structures at HIV LTR, ChIP assays were performed. The most specific inhibitor of DNA-PK, Nu7441 (12 μM), was used to assess the impact of DNA-PK inhibition in Jurkat-pHR’P-Luc cell line. ChIP assays were performed before and after activating the latent provirus with TNF-α in the absence or presence of the DNA-PK inhibitor **(Fig. 6A and B)**. The immunoprecipitated DNA was analyzed using two different primer sets. One primer set amplified the promoter region of the LTR (−116 to +4 with respect to the transcription start site) (**Fig. 6A**) and the other the nucleosome-1 (Nuc-1) region of the LTR (+30 to +134 with respect to the transcription start site) (**Fig. 6B**), an adjacent critical region, where slight epigenetic changes affect overall HIV gene expression (62, 64). The data obtained through immunoprecipitation using control IgG was subtracted from all samples as background and results were normalized with housekeeping GAPDH gene expression (-145 to +21) level. As shown in Fig. 6, at both promoter and Nuc-1 regions the latent proviruses had low levels of associated RNAP II, which increased by several folds following TNF-α treatment. However, in the presence of the DNA-PK inhibitor, TNF-α stimulation was significantly weak. The limited amount of RNAP II at HIV LTR in the presence of DNA-PK inhibitors is an indication of impaired HIV transcription. In parallel, we noted reduced recruitment of DNA-PK and P-TEFb (cyclinT1) in Nu7441 treated cells (**Figs. 6A and B**). Additionally, since pHR’P-Luc does not carry Tat gene, our data validate that Tat is not required for DNA-PK recruitment at LTR.

**Fig. 6:**
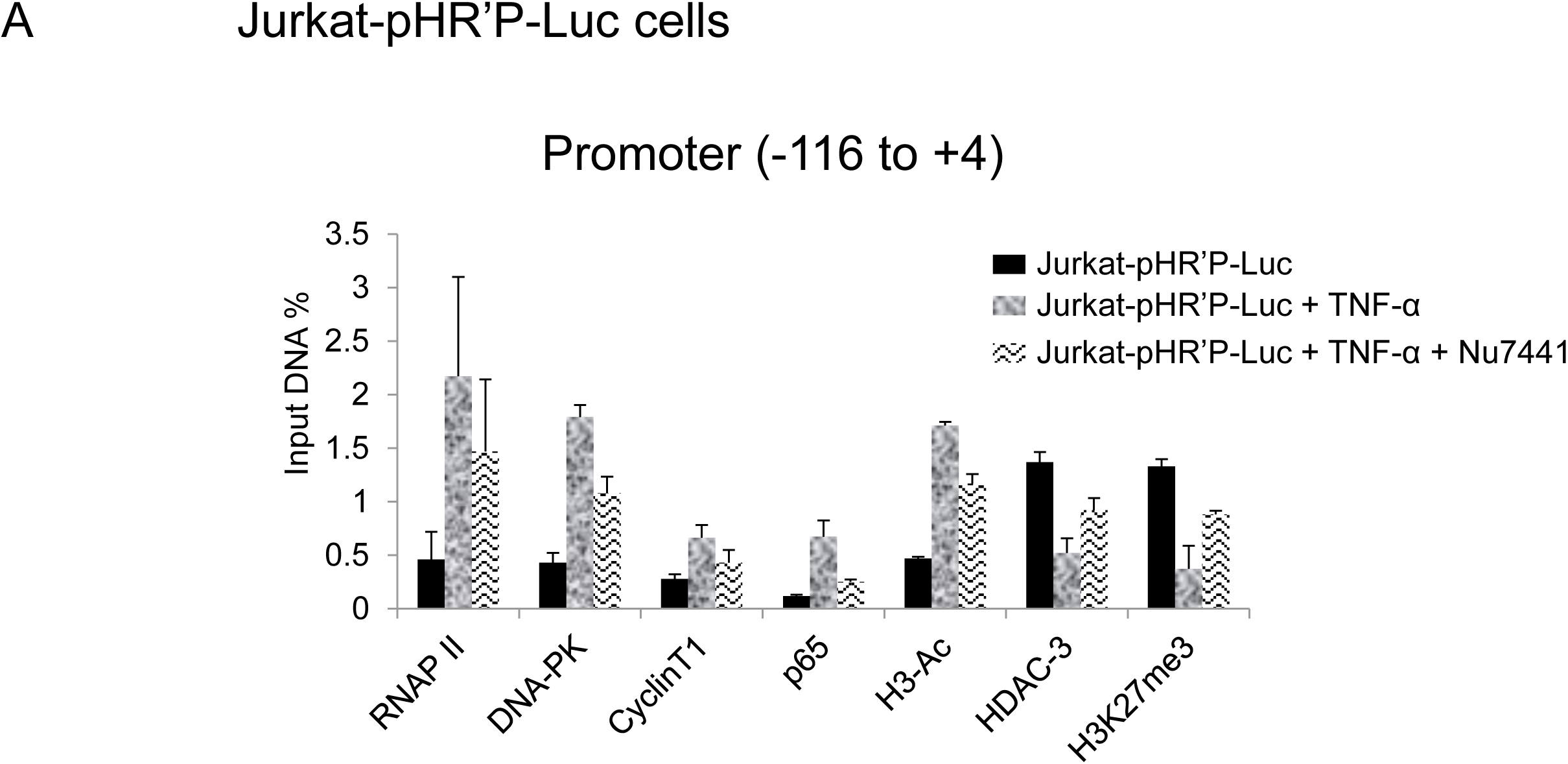
DNA-PK facilitates both the recruitment of transcription factors and establishment of euchromatin structures at HIV LTR. ChIP analysis in Jurkat-pHR’P-Luc before and after activating latent HIV provirus with TNF-α in the absence or presence of DNA-PK inhibitor Nu7441. The cellular chromatin was immunoprecipitated using indicated antibodies. The recovered DNA was analyzed using primer sets amplifying (**A**) the promoter region of the LTR. Data is represented as percentages of the input nuclear extract. Error bars represent the Mean ± SD of two independent experiments and three separate real-time qPCR measurements.

**Fig. 6:**
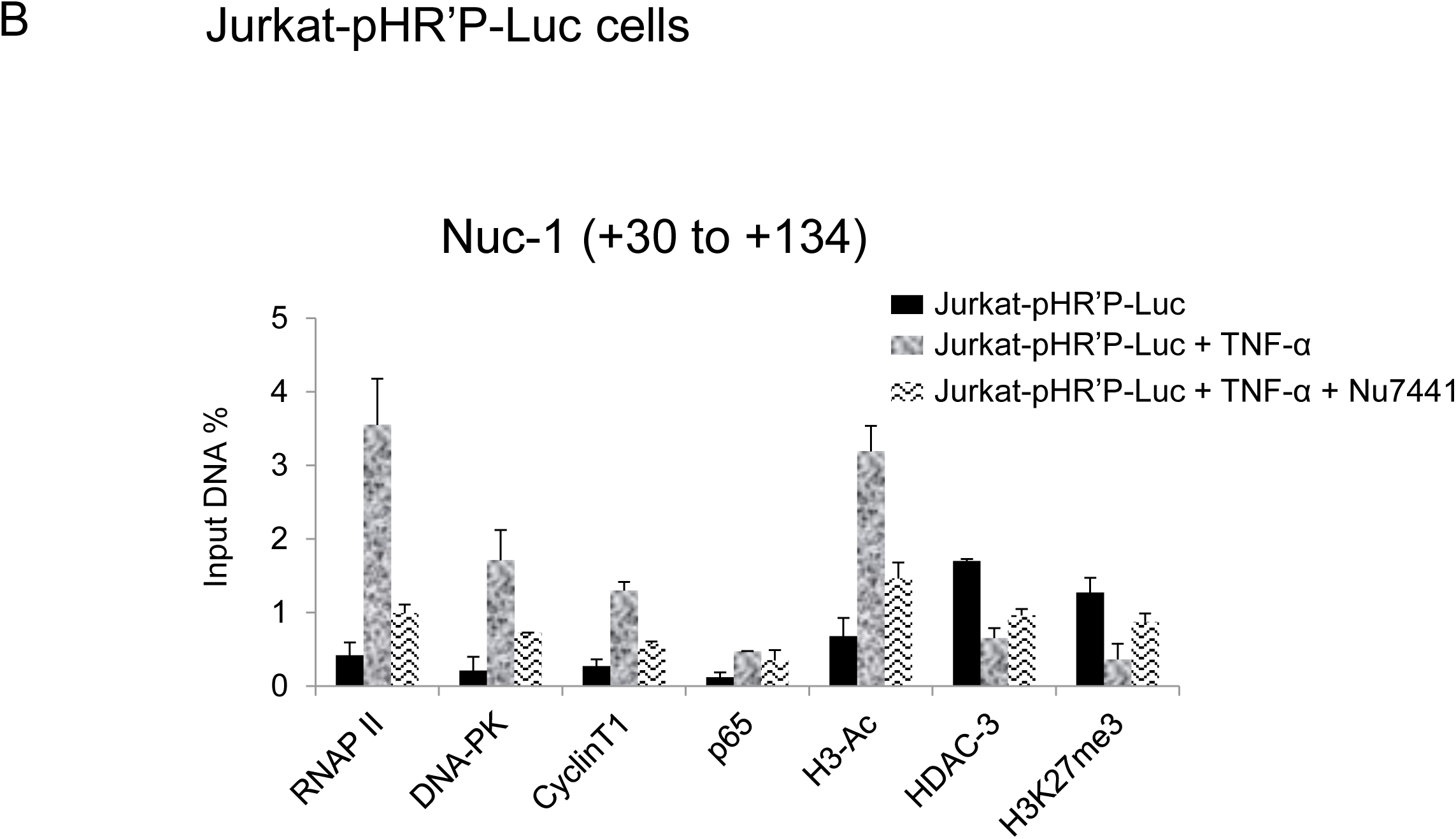
DNA-PK facilitates both the recruitment of transcription factors and establishment of euchromatin structures at HIV LTR. ChIP analysis in Jurkat-pHR’P-Luc before and after activating latent HIV provirus with TNF-α in the absence or presence of DNA-PK inhibitor Nu7441. The cellular chromatin was immunoprecipitated using indicated antibodies. The recovered DNA was analyzed using primer sets amplifying (**B**) the nucleosome-1 (Nuc-1) region of the LTR. Data is represented as percentages of the input nuclear extract. Error bars represent the Mean ± SD of two independent experiments and three separate real-time qPCR measurements.

TNF-α activates (nuclear translocation) NF-κB, and the greater recruitment of p65, a main subunit of NF-κB, marks the enhanced NF-κB binding at LTR after activation. Consistent with the fact that NF-κB binding sites resides in the promoter area of LTR, we found higher recruitment of p65 in promoter area after stimulation with TNF-α (**Fig. 6A**). Notably, we also observed some recruitment of p65 in the neighboring Nuc-1 region after stimulation with TNF-α. Given the ChIP resolution limit of ∼500 bp, an overlap of signals between adjacent regions, such as promoter and Nuc-1, was expected. After binding to the promoter, NF-κB is known to recruit histone acetyltransferases (HATs) at HIV LTR, which in turn acetylate the core histones (51). The augmented levels of acetylated histone H3 (H3-Ac), especially in the Nuc-1 region, marks the presence of higher levels of acetylated core histones at HIV LTR. We also observed the concomitant loss of histone deacetylases, HDACs, (HDAC3) from LTR further suggests that not only the recruitment of HATs, but dissociation of HDACs from LTR is responsible for establishing euchromatin structures at LTR. Following the treatment with Nu7441, the euchromatic marker H3-Ac went down; instead, the heterochromatic marker H3K27me3 got established at HIV LTR (**Fig. 6**). These heterochromatic markers indicate the presence of transcriptionally repressive heterochromatin structure at HIV LTR, resulting in reduced levels of RNAP II and DNA-PK at LTR, which eventually translate to reduced HIV transcription, gene expression and replication.

Consequently, it can be deduced from these results that DNA-PK enhances HIV gene expression by facilitating the establishment of transcriptionally active chromatin structures at HIV LTR. DNA-PK inhibitors restrict HIV transcription by reversing those changes and inducing the establishment of transcriptionally repressive heterochromatin structures at LTR.

### DNA-PK facilitates RNAP II pause release by phosphorylating and recruiting TRIM28 at LTR

DNA-PK has been shown to interact and catalyze the TRIM28 phosphorylation at serine 824 residue, p-TRIM28-(S824) (6, 8). This modification converts TRIM28 from a transcriptionally repressive factor to a transcriptionally active factor (6-8). We hypothesized that if DNA-PK plays a major role in TRIM28 activation by catalyzing its phosphorylation at serine 824, then DNA-PK inhibition or depletion in cells should reduce TRIM28 phosphorylation and thus contribute to the suppression of HIV gene repression, observed upon ablation of DNA-PK. We examined the effect of DNA-PK inhibitors on the TRIM28 phosphorylation at the serine 824 residue by western blot, using an antibody that detects specifically p-TRIM28-(S824). Jurkat cells were treated overnight with the DNA-PK inhibitor NU7441; next day cell media was replaced with fresh media containing Nu7441 along with TNF-α. After 30 minutes, nuclear-extracts were prepared and the levels of TRIM28 and p-TRIM28-(S824) were assessed. The results showed that DNA-PK inhibitor effectively reduced TNF-α-induced TRIM28-(S824) phosphorylation (**Figs. 7A and B**). However, the total TRIM28 protein levels were almost unaffected upon DNA-PK inhibition.

**Fig. 7:**
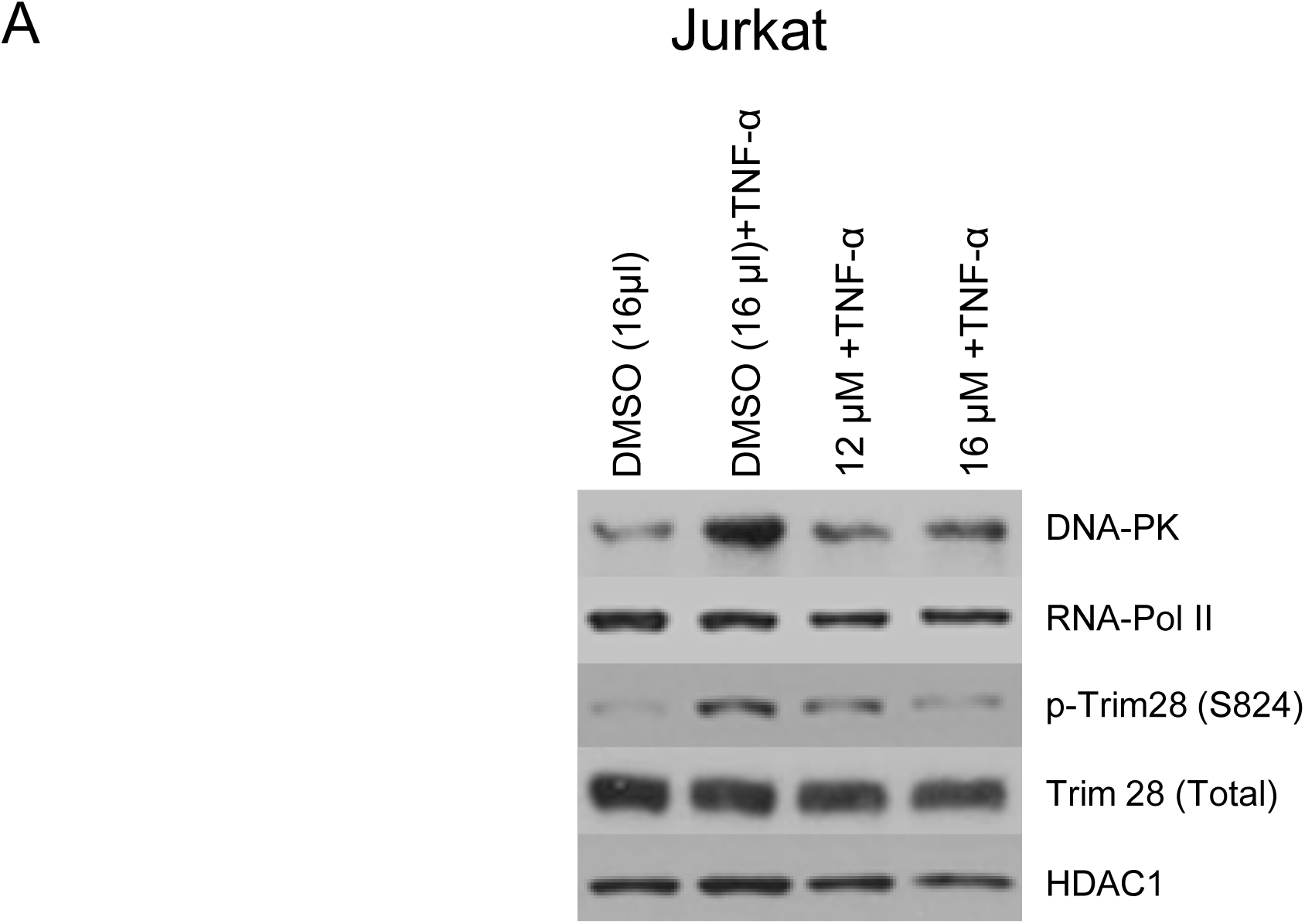
DNA-PK catalyzes the phosphorylation of TRIM28 and supports its recruitment at HIV LTR. Western blots were performed to evaluate the expression levels of Trim28 and phospho-Trim28 [p-TRIM28 (S824)] in (**A**) Jurkat cells treated with increasing concentration of the DNA-PK inhibitor Nu7441 with or without TNF-α.

**Fig. 7:**
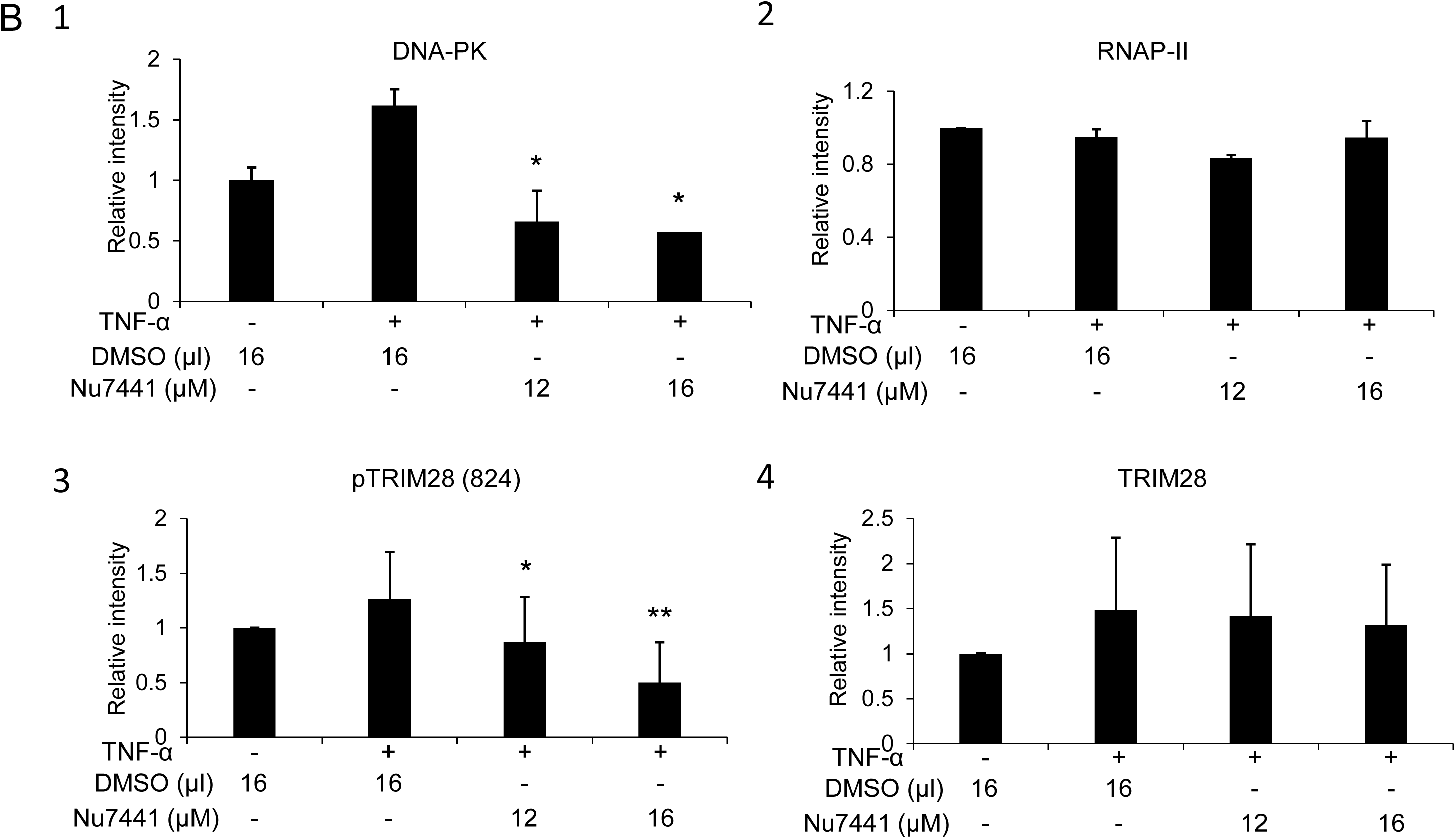
DNA-PK catalyzes the phosphorylation of TRIM28 and supports its recruitment at HIV LTR. Densitometric analyses of (**B**) Jurkat cells were performed using ImageJ software. Results represent the Mean ± SD of three independent analyses. Statistical significance is set as p < 0.05 (*) or 0.01 (**).

**Fig. 7:**
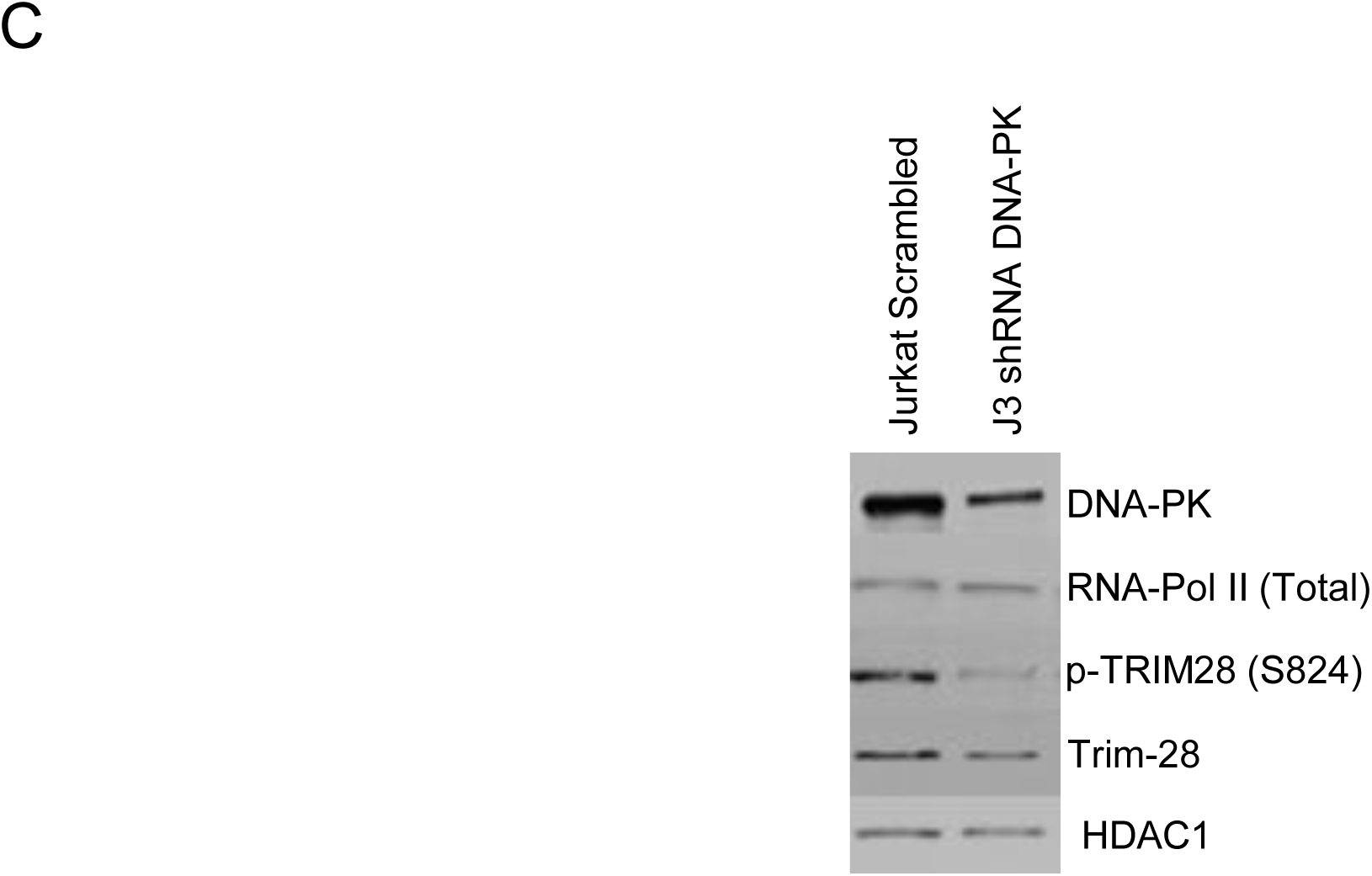
DNA-PK catalyzes the phosphorylation of TRIM28 and supports its recruitment at HIV LTR. Western blots were performed to evaluate the expression levels of Trim28 and phospho-Trim28 [p-TRIM28 (S824)] in (**C**) DNA-PK knockdown clone J3 and scrambled shRNA control.

**Fig. 7:**
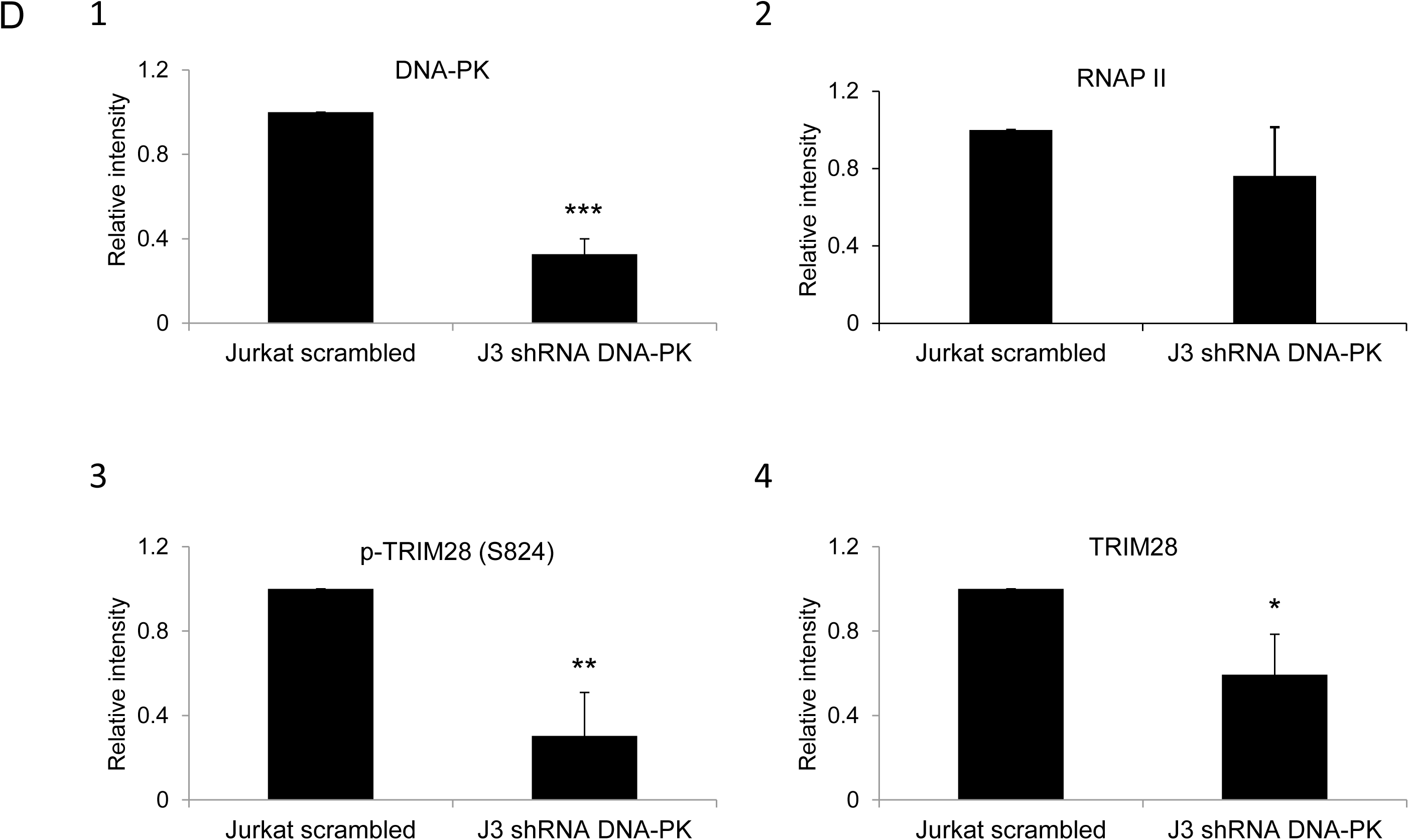
DNA-PK catalyzes the phosphorylation of TRIM28 and supports its recruitment at HIV LTR. Densitometric analysis of (**D**) DNA-PK knockdown J3 clone were performed using ImageJ software. Results represent the Mean ± SD of three independent analyses. Statistical significance is set as p < 0.05 (*), 0.01 (**) or 0.001 (***).

**Fig. 7:**
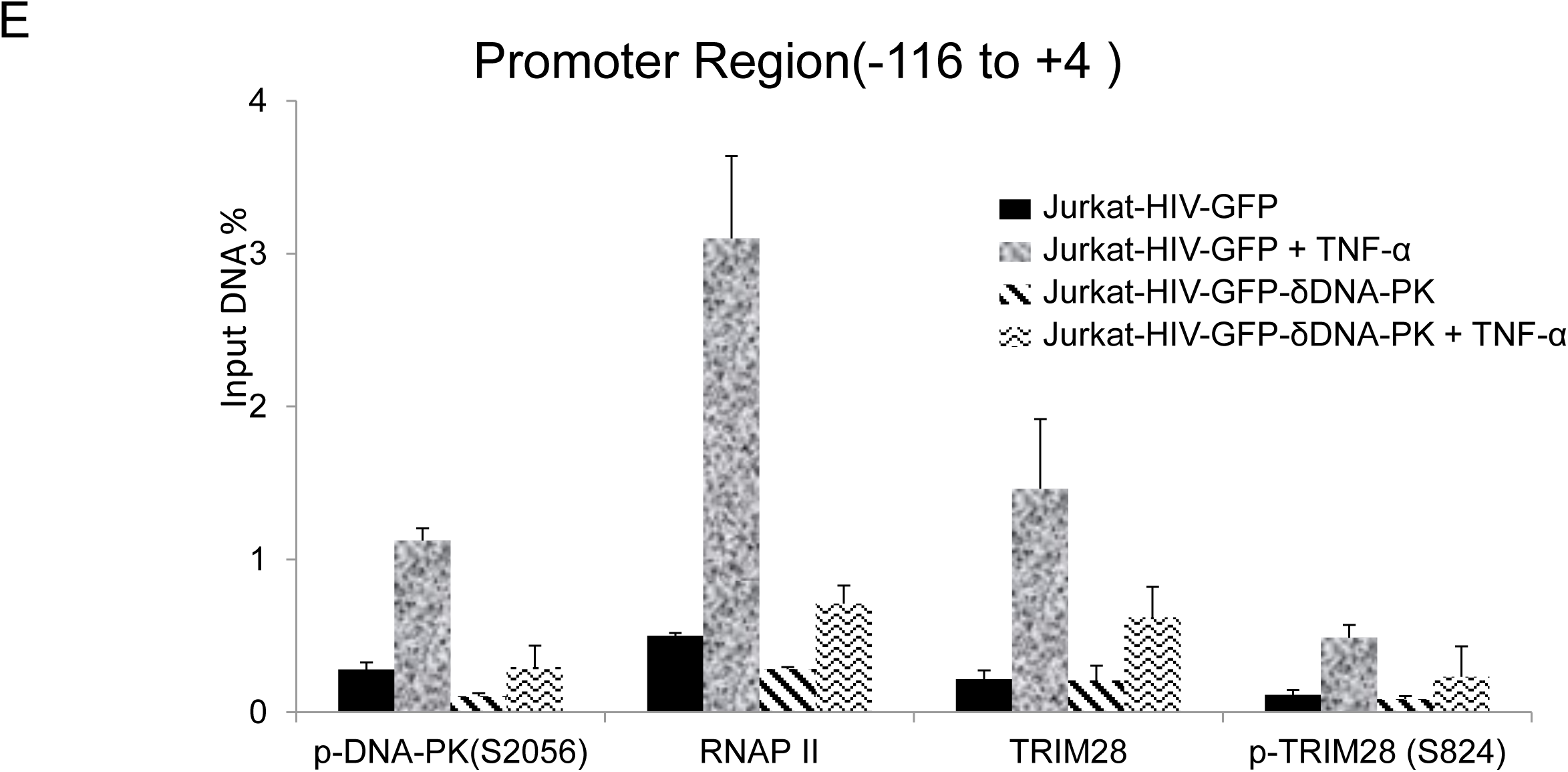
DNA-PK catalyzes the phosphorylation of TRIM28 and supports its recruitment at HIV LTR. Recruitment kinetics of TRIM28 and p-TRIM28-(S824) were assessed using ChIP assays in J3 and scrambled control, and also before and after reactivating latent HIV provirus. (**E**) Primer set amplifying the promoter region of the LTR. Data represent the percentages of the input. Error bars represent the Mean ± SD of three separate real-time qPCR measurements.

**Fig. 7:**
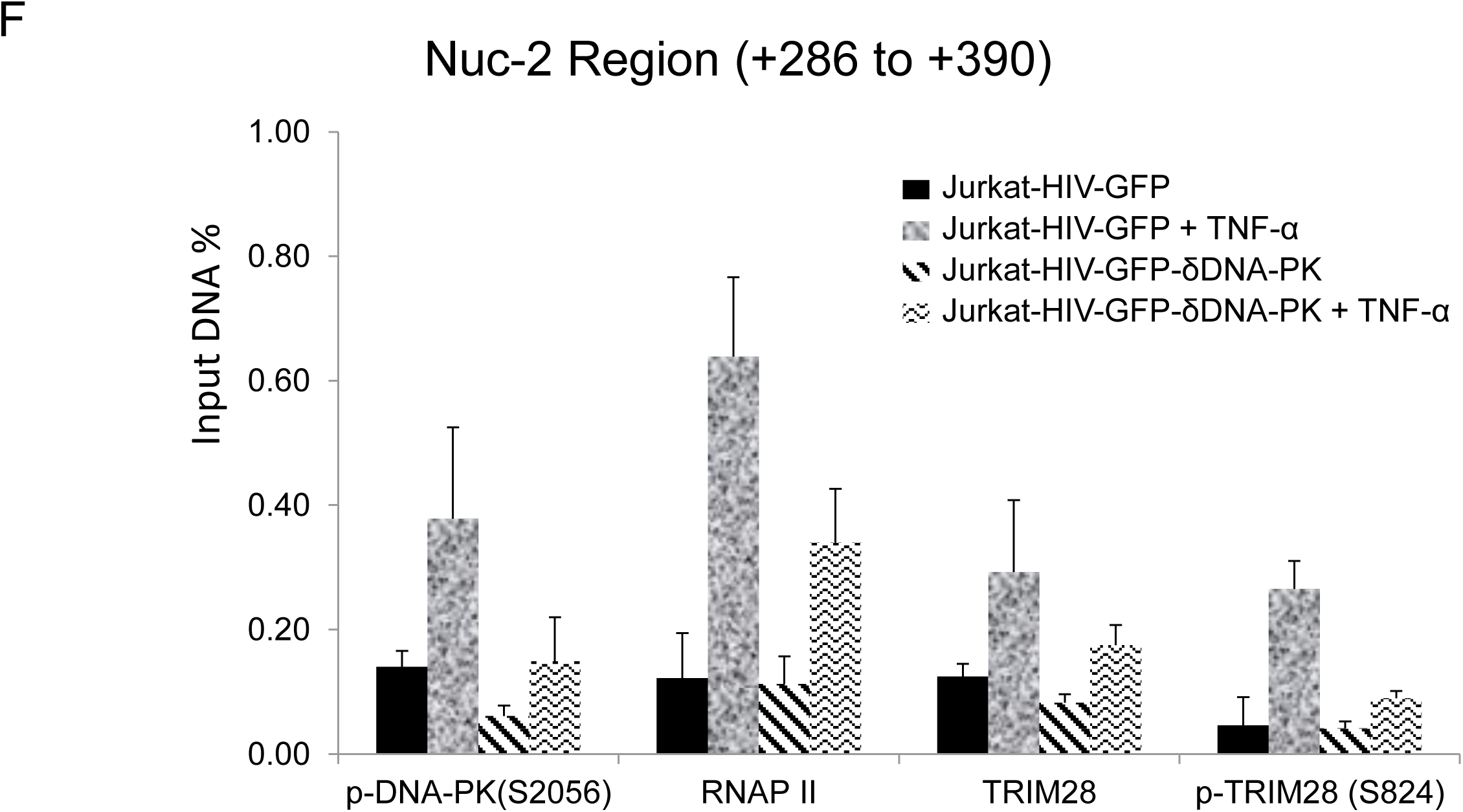
DNA-PK catalyzes the phosphorylation of TRIM28 and supports its recruitment at HIV LTR. Recruitment kinetics of TRIM28 and p-TRIM28-(S824) were assessed using ChIP assays in J3 and scrambled control, and also before and after reactivating latent HIV provirus. (**F**) primers amplifying the downstream Nuc-2 region of the LTR. Data represent the percentages of the input. Error bars represent the Mean ± SD of three separate real-time qPCR measurements.

We further evaluated the impact of DNA-PK knockdown on TRIM28 and p-TRIM28-(S824) in J3 clone of Jurkat cells. Following DNA-PK depletion, we observed clear reduction in p-TRIM28-(S824) levels (**Figs. 7C and D**). Interestingly, we also noted some reduction in the total TRIM28 levels. Overall data suggest that DNA-PK-mediated activation (phosphorylation) of TRIM28 may contribute to DNA-PK induced HIV transcription by relieving paused RNAP II. For confirming this notion, we next analyzed the recruitment of TRIM28 and p-TRIM28-(S824) in the presence and absence (depletion) of DNA-PK at HIV LTR.

The DNA-PK knocked-down clone J3 and control (scrambled shRNA) Jurkat cells carrying latent HIV provirus, pHR’-PNL-wildTat-d2GFP (Jurkat-HIV-GFP), in their genome were used in this analysis. The ChIP assays were performed before and after activating the cells with TNF-α (10 ng/ml) for 30 min. The immunoprecipitated DNA was analyzed using two primer sets, one amplifying the promoter region of the LTR (-116 to +4 with respect to the transcription start site, **Fig. 7E**) and another downstream Nuc-2 region of the LTR (+286 to +390 with respect to the transcription start site, **Fig. 7F**). In case of control wild type cells (scrambled shRNA), upon TNF-α treatment the latent provirus got reactivated, indicated by the higher recruitment of RNAP II. Interestingly, in parallel to the recruitment profile of DNA-PK after activation, we observed enhanced recruitment of TRIM28 at both promoter and Nuc-2 regions. However, we noticed relatively more enrichment of p-TRIM28-(S824) at Nuc-2 region after activation. In case of DNA-PK knockdown cells, we noticed impaired recruitment of TRIM28 and p-TRIM28-(S824) at HIV LTR. However, upon stimulation with TNF-α, we found some, but still highly reduced recruitment of TRIM28 and p-TRIM28-(S824). The decrease in the recruitment of TRIM28 and p-TRIM28-(S824) at HIV LTR following DNA-PK depletion, clearly suggests a contribution of DNA-PK in TRIM28 recruitment at HIV LTR. Moreover, restricted RNAP II levels at both promoter and Nuc-2 region of provirus in DNA-PK knockdown cells, further validates the highly reduced ongoing HIV gene expression, which again validates the importance of DNA-PK during the reactivation of latent HIV provirus. These findings reveal yet another way through which DNA-PK contributes to HIV transcription.

### DNA-PK inhibitors severely impair HIV-1 replication and transmission in MT-4 cells

The human T cell line MT-4 carrying Human T cell Lymphotropic Virus-1 (HTLV-1) is highly susceptible to HIV-1 infection and induced toxicity. We used these cells to examine the ability of Nu7026 and Nu7441 inhibitors to suppress spreading of HIV-1 infection and evaluate the cytotoxicity of these compounds. The cells were pre-treated for 24 hours with various concentrations of the two compounds (2, 4, 8, 16, 32 μM) or the highest concentration of DMSO used to dilute the compounds as control. The next day the cells were infected with X4_LAI.04_, washed two times in PBS and plated in 24-well plates (for HIV infection) and 96-well plates (for MTT assay) in medium containing the compounds. After 3 days, the concentration of p24 was evaluated in the medium and MTT assay was performed.

As shown in figures **8A and B**, both DNA-PK inhibitors strongly inhibit HIV-1 replication compared to the DMSO control. The compounds were not cytotoxic even at highest concentrations (**Figs. 8C and D**).

**Fig. 8:**
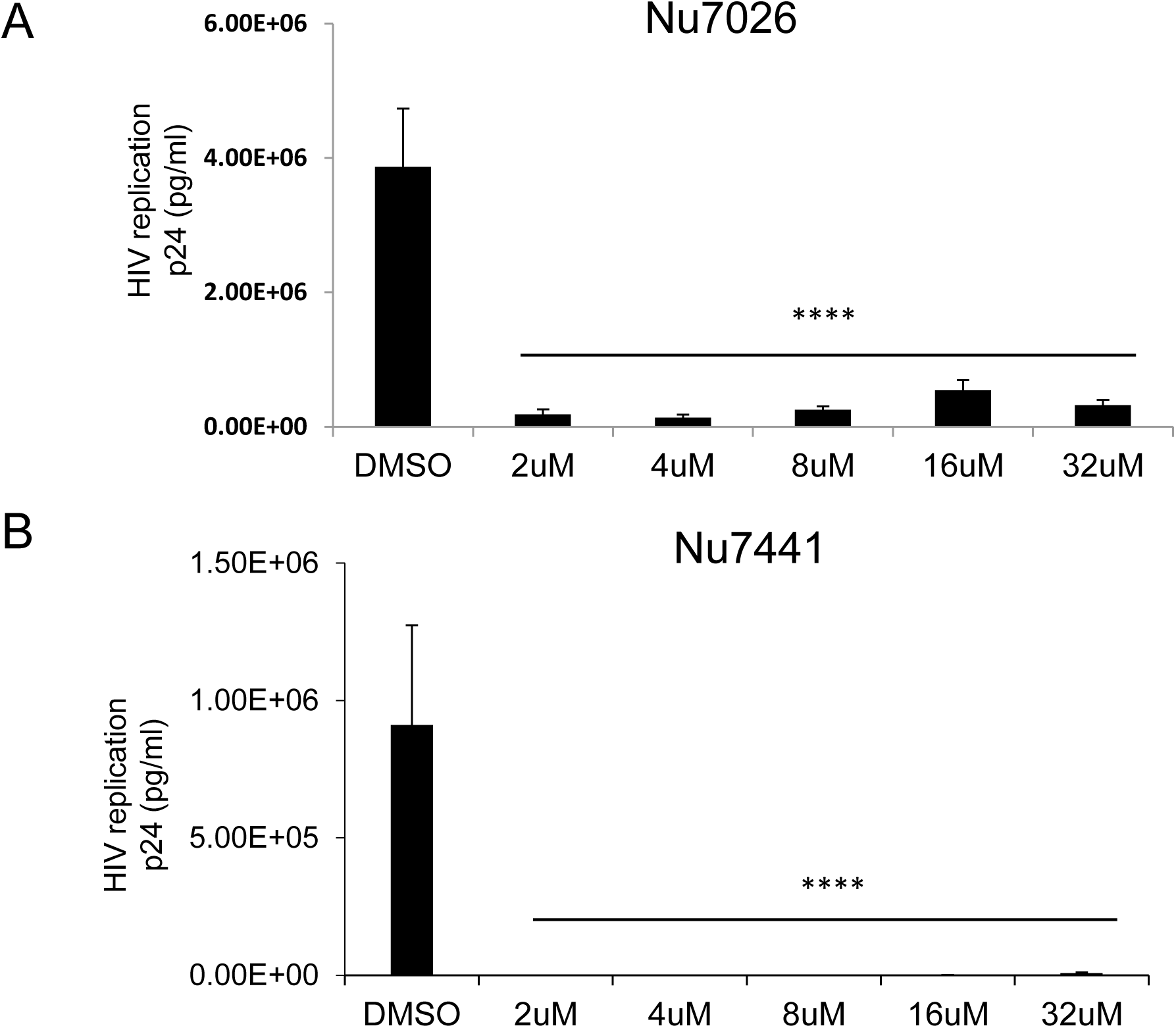
DNA-PK inhibitors completely block HIV replication in MT-4 T cells without showing cytotoxicity. MT-4 T cells were treated overnight with increasing concentrations of the DNA-PK inhibitors (**A**) Nu7026 and (**B**) Nu7441; DMSO was used as solvent control. Next day, the cells were infected with a replication competent X4 virus (HIV_LAI.04_). After 3 days of culture, cells supernatants were evaluated for p24 with Luminex (Luminexcorp). All the results represent the mean ± SD of four independent measurements. The p value of statistical significance was set at either; p < 0.0001 (****).

**Fig. 8:**
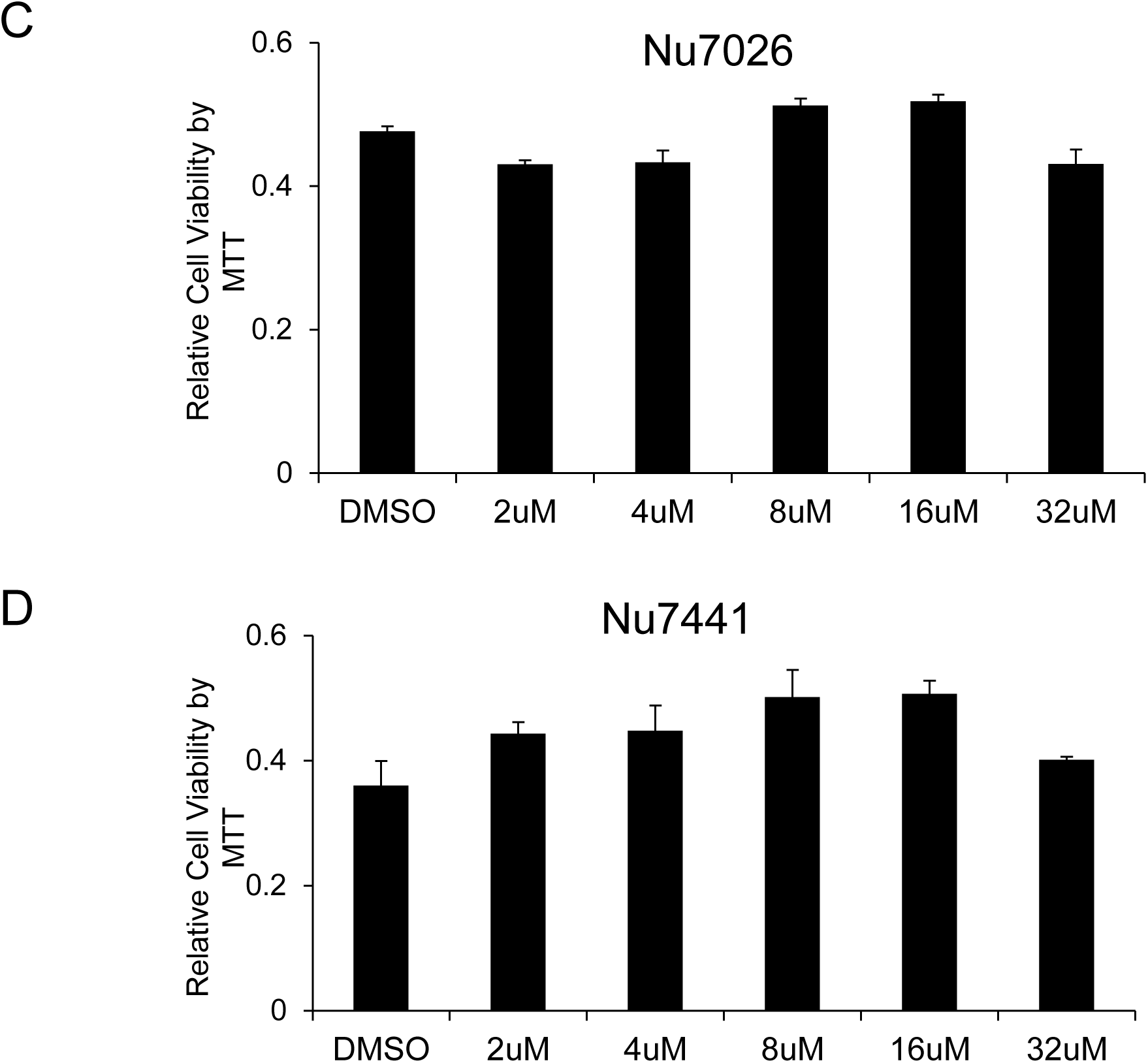
DNA-PK inhibitors completely block HIV replication in MT-4 T cells without showing cytotoxicity. The cytotoxicity of the inhibitors in MT-4 cells was evaluated with MTT assay. No cytotoxicity was detected in cells treated with (**C**) Nu7026 or (**D**) Nu7441, compared to the DMSO control. All the results represent the mean ± SD of four independent measurements.

### DNA-PK inhibitor restricts the reactivation of latent proviruses in peripheral blood mononuclear cells (PBMCs) from HIV patients

Overall, the results documented so far confirm an important role of DNA-PK in supporting HIV transcription and replication. We have shown that the small molecule inhibitors of DNA-PK efficiently restrict HIV gene expression, replication and the reactivation of latent proviruses in different cell types. To provide the clinical relevance to these findings, we evaluated the ability of DNA-PK inhibitors to restrict the reactivation of latent HIV proviruses present in the PBMCs of HIV patients on combination anti-retroviral treatment (cART). Ten million PBMCs from 4 HIV-infected patients undergoing successful cART treatment (undetectable viral load) were mixed and used in each sample. The cells were treated with either cART (2 NRTIs+1 PI, Zidovudine/Abacavir/Fosamprenavir) alone, with DNA-PK inhibitor, Nu7441 (12 μM) alone, or together with both Nu7441+cART, overnight. The next day, culture medium was replaced with fresh medium containing cART and/or Nu7441, and the cells were stimulated with PMA/ionomycin/IL-2 for 3 days to reactivate latent HIV provirus. Cellular RNA was isolated, cDNA was prepared and equal amounts of cDNA (5 ng) were analyzed by qPCR, with a primer set specific for the Nuc-2 region of LTR. The results were normalized with GAPDH gene amplification as loading control. We found that Nu7441, with or without cART, was able to restrict the reactivation of latent provirus, which occurs following the stimulation with PMA/ionomycin/IL-2. Notably, cART alone was not effective in controlling latent proviral reactivation and HIV gene expression (**Fig. 9A)**. Notably, Nu7441 also restricted the reactivation of latent provirus which was partially reactivated while thawing the PBMCs (compare lanes 1 and 3, 5). Viability of PBMCs was not compromised by the treatments (**Fig. 9B**). Instead, drug treatments, by virtue of repressing HIV replication, enhanced cell survival. This result validates our hypothesis that transcriptional inhibitors, such as DNA-PK inhibitors, can be used to suppress the ectopic reactivation of latent HIV provirus, a main contributor to viral blips, even in HIV patients undergoing successful cART.

**Fig. 9:**
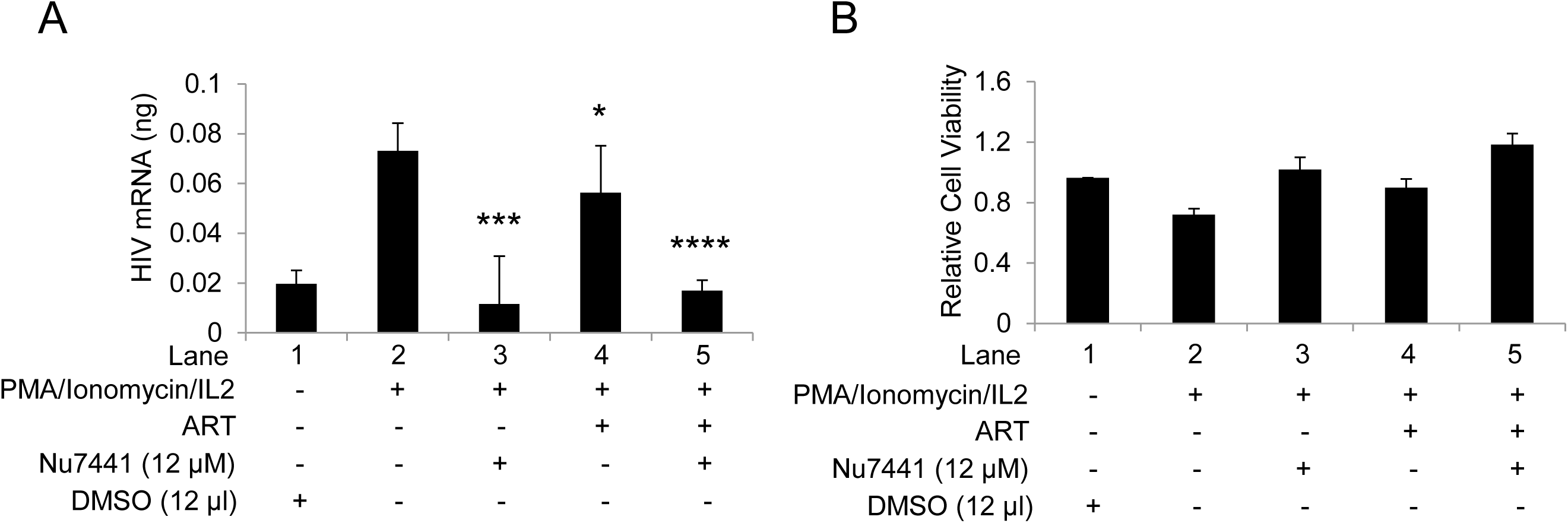
DNA-PK inhibitors restrict the reactivation of latent provirus in PBMCs of HIV-infected patients. PBMCs from HIV-infected patients were treated with either ART alone, Nu7441 alone or both together overnight. The day after the medium was replaced with fresh medium containing the drugs and the cells were stimulated for 3 days with PMA/ionomycin/IL-2 to reactivate latent HIV provirus present in PBMCs. Cellular RNAs were isolated and evaluated via RT-qPCR (**A**). Cell viability was examined by MTS assays (**B**). The results were reproduced 3 more times, again by mixing PBMCs of 4 different patients each time. Presented data are the Mean ± SD for 4 experiments.

### DNA-PK promotes HIV transcription by supporting different aspects of HIV transcription

In order to summarize our results, we present the following model (**Fig. 10**). Based on our previous (61) and present studies, we propose that DNA-PK promotes the initiation phase of HIV transcription mainly by directly catalyzing the phosphorylation of RNAP II CTD at Ser5. It also facilitates the elongation phase of transcription by both directly catalyzing the phosphorylation of Ser2 of CTD and through the recruitment of P-TEFb at LTR. We also found that DNA-PK, by phosphorylating TRIM28 at the specific serine residue (S824), converts it from a RNAP II pausing factor into an elongation-supporting factor. Thus, DNA-PK also augments HIV transcription by relieving the RNAP II pausing. Altogether, DNA-PK enhances the HIV transcription by supporting the multiple phases of transcription, namely initiation, RNAP II pause release and elongation. Therefore, besides controlling replication, DNA-PK inhibition may be used to prevent the reactivation of the latent HIV proviruses.

**Fig. 10:**
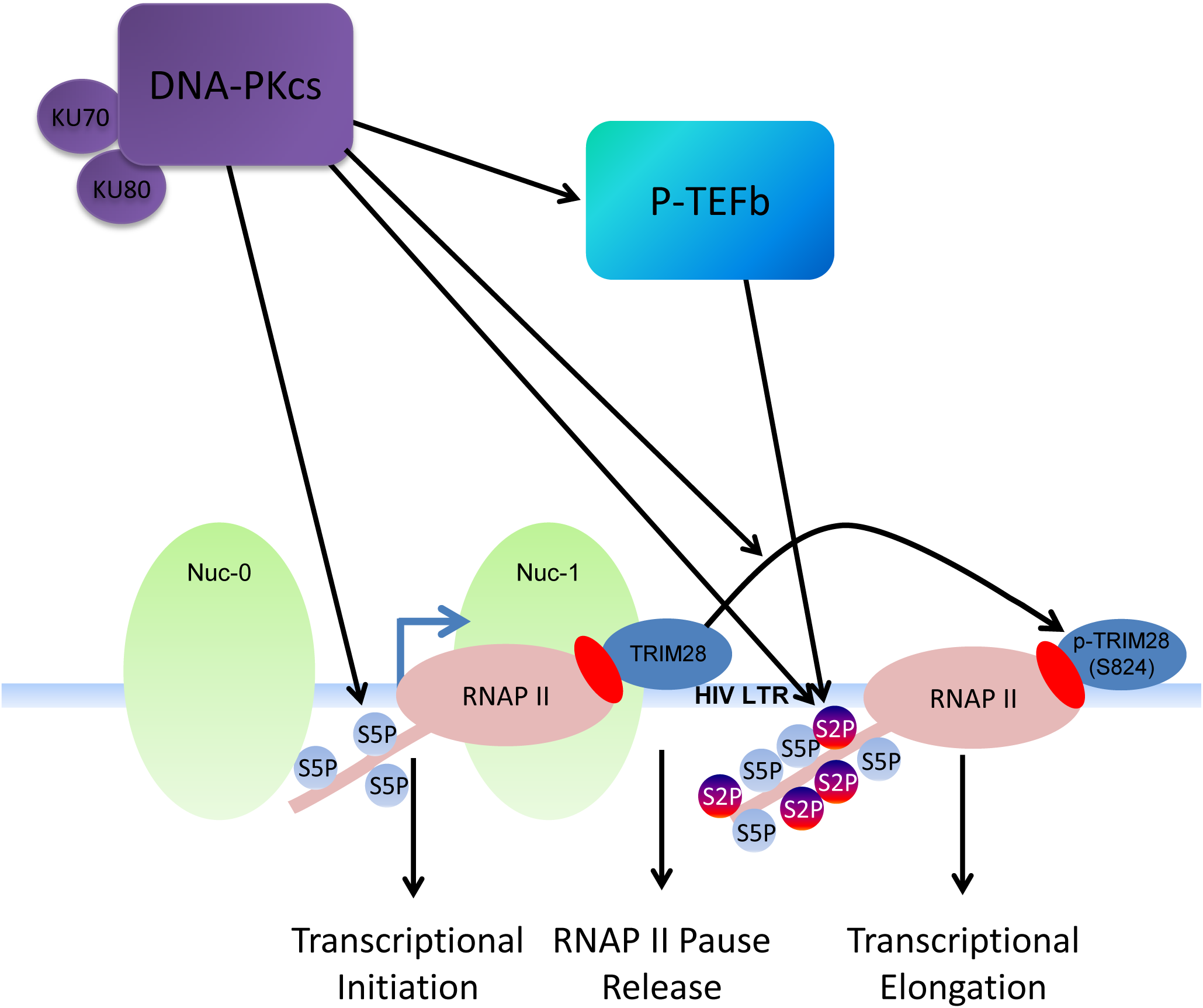
DNA-PK facilitates HIV transcription by targeting multiple mechanisms. In our model presented here we suggest that DNA-PK supports the initiation phase of transcription by catalyzing the phosphorylation of RNAP II CTD at Ser5. DNA-PK facilitates transcriptional elongation by enhancing phosphorylation of RNAP II CTD at Ser2, both via directly catalyzing and through the recruitment of P-TEFb at HIV LTR. DNA-PK relieves the RNAP II pausing by catalyzing the phosphorylation of TRIM28.

## Discussion

Unraveling the molecular mechanisms that control HIV life cycle, especially transcription, replication and reactivation of the latent provirus, is crucial for cure and eradication of HIV. The studies presented herein provide strong and compelling evidence for the important role of DNA-PK in supporting various stages of HIV transcription, which subsequently promote HIV replication and reactivation from latency. We discovered that DNA-PK promotes transcriptional initiation by catalyzing the phosphorylation of CTD at Ser5; relieves RNAP II pausing by phosphorylating TRIM28 at S824; facilitates transcriptional elongation by both catalyzing the phosphorylation of CTD at Ser2 and promoting of P-TEFb recruitment at HIV LTR. Accordingly, we noted that DNA-PK inhibition or depletion severely impairs HIV transcription, replication and reactivation of latent provirus. Overall, these results reveal the underlying molecular mechanism through which DNA-PK facilitates HIV gene expression, and suggest that DNA-PK inhibitors may have a potential role as adjunctive agents in the treatment of HIV infection.

The important role of DNA-PK, especially during non-homologous end joining (NHEJ) DNA-repair pathway, is well-established (17, 32, 55). Our earlier work and studies by others have clearly documented the interaction between DNA-PK and RNAP II (16, 37, 57, 61). Moreover, for the first time, we have shown the importance of DNA-PK during transcription, its presence at HIV LTR, and confirmed DNA-PK as an integral part of the RNAP II holoenzyme (61). Recently, our findings have been confirmed, and a similar mechanism involving DNA-PK has been proposed for the transcription of inducible cellular genes (7, 9, 24). To extend these findings further, in this investigation we have demonstrated that DNA-PK enhances HIV transcription by facilitating the various stages of transcription. We also noted reduced nuclear level of DNA-PK in unstimulated cells, which upregulates upon cell stimulation (**Figs. 2, 4, and 7**), suggesting a clear role of reduced DNA-PK levels in supporting HIV latency in quiescent cells.

The observation of enhanced recruitment of DNA-PK at HIV LTR during HIV gene expression and the synergistic recruitment kinetics of DNA-PK with RNAP II along HIV genome (61) prompted us to explore further the role of DNA-PK during HIV transcription, replication and latent proviral reactivation. In this investigation we utilized highly specific DNA-PK inhibitors that are known to induce phenotypes similar to those seen in DNA-PK deficient cell lines and DNA-PK knockout mice (12, 15). The knockdown studies reaffirmed the specificity of inhibitors towards DNA-PK. Analysis of the reactivation of latent HIV provirus expressing *luciferase* reporter gene under the control of HIV LTR promoter unequivocally demonstrated that DNA-PK inhibitors are very strong repressors of HIV transcription and replication (**Fig. 1**), since TNF-α, a strong inducer of latent provirus, was unable to increase HIV gene expression in the presence of optimal concentrations of specific DNA-PK inhibitors. The dose-dependent nature of this repression was also confirmed in experiments with two different cell lines, representing CD4+ lymphoid and myeloid cells, the major targets of HIV infection in patients. The physiological relevance of these results was established by confirming the inhibitory effect of DNA-PK inhibitors in suppressing the reactivation of latent HIV provirus in primary CD4+ T cells, infected either *ex vivo* (**Fig. 1**) or obtained from HIV patients (**Fig. 9**). We noted some cytotoxicity of DNA-PK inhibitors in primary CD4+ T cells at concentrations over 16 μM; therefore, the inhibitors were always used within physiologically non-toxic range (**Fig. 1**). Under these conditions, DNA-PK inhibitors even supported the survival of primary cells isolated form HIV patients (**Fig. 9**), and also in MT-4 cells (**Fig. 8**). This observation confirms that DNA-PK-mediated inhibition of HIV replication averts HIV-mediated cytotoxicity.

In our prior study, we have confirmed the presence of DNA-PK at HIV LTR and, by performing *in vitro* kinase assays, demonstrated that DNA-PK was able to catalyze RNAP II CTD phosphorylation *in vitro*, using purified DNA-PK protein (61). To extend those *in vitro* findings, here we demonstrated that DNA-PK does, in fact, phosphorylate RNAP II CTD *in vivo*. We have shown that following the treatment of cells with either specific DNA-PK inhibitor or upon DNA-PK depletion (**Figs. 2 and 3**), RNAP II CTD phosphorylation (Ser2 and Ser5) becomes highly impaired in both Jurkat and THP-1 cells. Therefore, our results demonstrate that DNA-PK enhances HIV transcription by promoting both the initiation and elongation phases of HIV transcription.

The interaction between DNA-PK and P-TEFb subunits has also been documented (1, 36). These findings impelled us to determine if DNA-PK also increases the RNAP II CTD phosphorylation by bringing P-TEFb along with it to HIV LTR. In fact, upon DNA-PK depletion, in parallel to impaired DNA-PK recruitment, we noted severely decreased recruitment of both P-TEFb subunits at HIV LTR (**Fig. 4**). Even after proviral reactivation with TNF-α, we observed highly restricted recruitment of P-TEFb, accordingly we noted that DNA-PK depletion impacted the cellular Tat levels, the main recruiter of P-TEFb at LTR (**Figs. 4 and 5C**).

In DNA-PK knockdown cells, we discovered that HIV, which expresses functionally impaired Tat (H13L), a mutation prevalent in latent proviral isolates (43, 59, 70), was almost ineffective in supporting HIV transcription. Moreover, HIV carrying wild-type Tat was also highly defective in expressing the reporter *GFP* gene (**Figs. 5A and B**). The finding that DNA-PK depletion results reduced HIV protein expression, such as Tat (**Fig. 5C**), was further supported by the impaired latent proviral reactivation and replication, in DNA-PK inhibited and depleted cells (**Figs. 1 and 2**). Moreover, previously we and others have revealed that Tat phosphorylation catalyzed by DNA-PK is essential for its optimal activity (61, 71). Hence, besides Tat levels, the deficient phosphorylation of Tat, due to DNA-PK inhibition or depletion, may contribute to reduced Tat activity and P-TEFb recruitment; this is the subject for future investigations. Altogether, our results confirm the important role of DNA-PK in supporting HIV transcription.

Given the fact that chromatin structure around the promoter of a gene controls its expression, we investigated which specific epigenetic changes are induced by DNA-PK that promote HIV transcription, and the reversion of those changes upon DNA-PK inhibition (**Fig. 6).** Similar to our prior findings, we observed greater recruitment of DNA-PK at LTR, upon latent proviral activation. In parallel to DNA-PK, we also noticed higher recruitment of RNAP II and co-recruitment of other transcription factors, such as NF-κB and P-TEFb at LTR, which clearly indicates enhanced ongoing transcription from LTR. Simultaneously, we found accumulation of euchromatic marks, such as acetylation of core histones (H3-AC), but removal of heterochromatic marks and enzymes involved, such as H3K27me3 and of HDAC3. All these changes facilitate the establishment of transcriptionally-active euchromatin structures at HIV LTR, which support HIV transcription. However, upon inhibition of DNA-PK, the whole process just got reversed, and we noted the loss of euchromatic marks and gathering of heterochromatic marks at HIV LTR, which leads to the generation of transcriptionally-repressive heterochromatin structures at HIV LTR. Heterochromatin structures suppress transcription by impeding the access of transcription machinery as confirmed by reduced levels of RNAP II and other transcription factors at LTR.

Next, we investigated how DNA-PK promotes HIV transcription by regulating TRIM28 activity (**Fig. 8**). Recently, TRIM28 was confirmed as one of the factors that facilitate RNAP II pausing at several cellular genes, including HIV (8, 40). DNA-PK has been shown to interact directly with TRIM28 and catalyze the TRIM28 phosphorylation at serine 824 residue, p-TRIM28-(S824) (6, 8). This modification makes TRIM28 functionally active and converts it from a transcriptionally pausing factor to a transcriptionally active factor, which allows the release of paused RNAP II and promotes transcriptional elongation (6-8). Thus, like SPT5 component of another pausing factor DSIF complex, TRIM28 first promotes RNAP II pausing. However after phosphorylation at S824, it becomes activator of transcription and facilitates the release of paused RNAP II (3, 8). We showed that TRIM28 is indeed the substrate of DNA-PK, as specific inhibition or depletion of DNA-PK resulted in reduced phosphorylation of TRIM28 at S824. However, during this study we noted that the P-TEFb inhibitor, 5,6-Dichloro-1-*beta*-D-ribofuranosylbenzimidazole (DRB) does not affect TRIM28 phosphorylation, suggesting that TRIM28 S824 phosphorylation is not mediated by P-TEFb (not shown). The data suggest that limited phosphorylation of TRIM28, upon DNA-PK inhibition or depletion stabilizes the pausing state of RNAP II and contributes to decreased HIV transcription. Subsequently, in ChIP assays, in parallel to DNA-PK recruitment at LTR, we found the enhanced recruitment of TRIM28 and phosphorylated TRIM28 (S824) at HIV LTR. Especially, at Nuc-2 region of LTR, we noticed relatively higher fold enrichment of p-TRIM28(S824) after stimulation with TNF-α. The accumulation of ρ-TRIM28(S824) marks the presence of the transcription-supporting form of TRIM28 and thus indicates the transformation of paused RNAP II into a processive elongating RNAP II. However, after DNA-PK depletion we found very limited levels of p-TRIM28(S824) at LTR, even after activation, demonstrating that DNA-PK is the main kinase that catalyzes the phosphorylation of TRIM28. However, besides DNA-PK, ATM has also been shown to catalyze the phosphorylation of TRIM28 at S824 (7, 19, 56). It should be noted that following DNA-PK knockdown, we found reduced presence of both TRIM28 and p-TRIM28(S824) at LTR, clearly suggesting a primary role of DNA-PK in the recruitment of TRIM28 at HIV LTR. It appears that ATM cannot compensate for the lack of this activity of DNA-PK. The detailed investigation of this important mechanism is a subject for future investigations utilizing a combination of biochemical, genetic and genomics approaches.

We also evaluated the effect of the DNA-PK inhibitors Nu7026 and Nu7441 on HIV-1 replication in MT-4 cells, using a prototypical replication competent X4 variant of HIV-1 (HIV-1 X4_LAI.04_). Both inhibitors were able to completely inhibit HIV replication at low doses (**Figs. 8A and B**) and well tolerated by the cells (**Figs. 8C and D**).

Finally, we evaluated the clinical relevance of the DNA-PK inhibitor Nu7441 by assessing its impact in restricting the reactivation of latent provirus using PBMCs from HIV-infected patients (**Fig. 9**). We found an almost complete block of HIV gene expression in the cells treated with Nu7441, even after stimulation of latently infected cells with α-CD3/CD28 antibodies. Notably, HAART treatment, which is very effective in controlling HIV replication, was not effective in controlling HIV gene expression of the reactivated latent provirus. These results strongly argue for the inclusion of transcriptional inhibitors, such as DNA-PK inhibitors, as a supplement to HAART regimens for suppressing transient reactivation of latent proviruses. While continuous treatment with HAART prevents the infection of neighboring cells, in sanctuary sites, including gut-associated lymphoid tissue (GALT) and central nervous system (CNS), these transient episodes can lead to spread of viral reservoir and contribute to immune activation.

Thus, our findings unraveled the molecular mechanisms utilized by DNA-PK to enhance HIV transcription and confirmed the physiological relevance of DNA-PK inhibitors in restricting HIV gene expression, replication and reactivation of latent provirus.

We propose a model to depict these multiple activities of DNA-PK (**Fig. 10**). Briefly, our data suggest that DNA-PK enhances HIV transcription by supporting the initiation phase of transcription by catalyzing the phosphorylation of RNAP II CTD at Ser5. By catalyzing the phosphorylation of TRIM28, specifically at residue S824, DNA-PK converts TRIM28 from a pausing factor to an elongating factor, thus relieving RNAP II pausing. Additionally, DNA-PK supports the elongation phase of HIV transcription by enhancing the processivity of RNAP II through augmenting the CTD phosphorylation at Ser2, both directly and via facilitating the recruitment of P-TEFb at HIV LTR. Consequently, by repressing all these pathways, DNA-PK inhibitors show such a profound effect on HIV replication and latent proviral reactivation.

## Conclusions

Our results confirm the important role of DNA-PK in both HIV transcription and replication. By performing luciferase and reverse transcriptase assays, we demonstrated that the restriction of DNA-PK activity via specific inhibitors strongly represses HIV transcription and viral replication, and confirmed these results not only in lymphoid and myeloid cell lines, but also in primary CD4+ T cells. Subsequently, by chromatin immunoprecipitation assays we showed that specific DNA-PK inhibitors repress HIV transcription by inducing transcriptionally repressive chromatin structures at HIV LTR, and inhibit both the initiation and elongation phases of HIV transcription by severely restricting the phosphorylation of the carboxyl terminal domain (CTD) of RNAP II.

Altogether, these results confirm the importance of DNA-PK in HIV replication and define the underlying molecular mechanism through which DNA-PK facilitates HIV gene expression. Moreover, these results support a potential therapeutic use of specific DNA-PK inhibitors as supplements in HAART/cART regimens to further enhance the effectiveness of anti-HIV therapy and might even control the rate of HIV-associated cancers, as DNA-PK inhibitors are under trial for cancer therapy.

## Materials and Methods

### Reagents

DNA-PK Inhibitors: Nu7026 (Cat. # S2893) and Nu7441 (Cat. # S2638) from Selleckchem, while IC86621 (Cat. # 404009-40-1) from Sigma-Aldrich, was purchased. Stock solutions were made in DMSO at 5 mM concentration. DNA-PKcs shRNA vectors (Cat # sc-35200-SH) were purchased from Santa Cruz Biotech. In some experiments, we also used the lentiviral vector pHR’Sin-Puro with shRNA, which carry puromycin resistance gene and expresses shRNA through H1 Pol-III promoter. The shRNAs to DNA-PKcs (5′-GAACACTTGTACCAGTGTT) or (5′-GATCGCACCTTACTCTGTT) and scrambled control (5’-TTGATGCACTTACTAGATTAC), were cloned. A total of 4 to 6 μg of VSV-G pseudotyped lentiviral vectors carrying specific or control shRNA were used for knockdown.

### Cell culture conditions

The J1.1, U1, Jurkat, THP-1, U937, MT-4, primary T cells and PBMCs were cultured in RPMI medium with L-glutamine(Invitrogen), 10% fetal bovine serum (FBS), penicillin (100 IU/ml), Ciprofloxacin (10 μM) and streptomycin (100 μg/ml). Primary cells were also supplied with primocin (invivoGen), 25 mM HEPES (pH 7.2) and IL-2 (20 U/ml of recombinant human IL-2 [R&D Systems, Inc. or Roche]). The cells were grown in incubator maintaining CO_2_ at 5% and temperature at 37°C.

### Cell lines and Viruses

THP-1, Jurkat and primary cells were infected with replication-incompetent HIV (pHR’P-Luc, pHR’-PNL-wildTat-d2GFP or pHR’-PNL-H13LTat-d2GFP, carrying reporter genes either *luciferase* or *GFP* under the control of the HIV LTR promoter. VSV-G-pseudotyped HIV particles were packaged as previously described (59). The latently infected promonocytic U1 and T-lymphocytic J1.1 lines were kindly provided by NIH AIDS reagent program. These cell lines harbor integrated replication-competent latent provirus and constitutively express very low amounts of HIV proteins. The cells were infected with replication-competent HIV-1 NL4-3 and X4_LAI.04_ using standard protocol (35). The cell viability was regularly evaluated by propidium iodide staining.

### Luciferase Assay

THP-1-pHR’-P-Luc, Jurkat-pHR’-P-Luc and primary CD4+ T cells were plated in 6-well plates at a concentration 10^6^ cells/ml, in complete RPMI medium. The cells were pre-incubated with different concentrations of DNA-PK inhibitors or with dimethyl sulfoxide (DMSO) as control, overnight. Since the inhibitors were dissolved in DMSO, a set of cells was treated with the highest used concentration of DMSO as negative/background control. Next day, cells were supplied with fresh media containing inhibitors. The latent HIV provirus in the Jurkat and THP-1 cell lines was reactivated by treatment with Tumor Necrosis Factor alpha (TNF-α, 10 ng/ml), whereas in primary CD4+ T cells by stimulation through T cell receptor (TCR) with α-CD3/CD28 antibodies-coated beads (25 μl/10^6^ cells). After 48 hours, the cells were harvested and washed three times with phosphate-buffered saline (PBS). Subsequently, Bradford assays (Bio-Rad) were performed and equal amount of cell extracts were used in each luciferase reaction. Luciferase levels in the cells were assessed using a Promega commercial kit (Cat# E4530; Madison, WI). Briefly, the cells were lysed for 30 minutes at room temperature with passive lysis buffer and centrifuged at 14000 rpm for 2 minutes. 5 μl of samples were added to individual wells of a 96-well plate, followed by 70 μl of luciferase substrate/assay buffer. Samples were tested in triplicate. Luminescence was read in a Microplate Luminometer (Bio-Rad or Turner Biosystems).

### Cell cytotoxicity: MTS assay and MTT assay

The cytotoxicity of DNA-PK inhibitors in Jurkat, THP-1 and primary CD4+ T cells carrying pHR’P-Luc, was assessed using 3-(4,5-dimethylthiazol-2-yl)-5-(3-carboxymethoxyphenyl)-2-(4-sulfophenyl)-2H-tetrazolium (MTS) reagent (Promega) following the supplier’s protocol. Briefly, the cells were seeded (3x10^3^ cells/well) in 96-well plates and incubated with increasing concentrations of Nu7441 or Nu7026 for 5 days. Then the cells were incubated for 4h with the MTS reagent directly added in the culture wells. Subsequently, the optical density was measured at 490 nm, using a visible light 96-well plate reader.

The cytotoxicity of DNA-PK inhibitors in MT-4 cells was evaluated with 3-(4, 5-dimethylthiazolyl-2)-2, 5-diphenyltetrazolium bromide (MTT) reagent (Sigma-Aldrich). The cells were pre-treated for 24 h with the DNA-PK inhibitors Nu7026 and Nu7441 at different concentrations and DMSO as control, and seeded (2x10^4^ cells/well) in 96-well plate in 100 μl of RPMI 10% FBS. The day after the cells were infected with X4_LAI.04,_ washed two times in PBS and plated again in a 96 well-plate. After 3 days, the MT-4 were incubated for 2 h at 37 °C with MTT reagent, then treated with 100 μl of DMSO per well for 30 minutes and the optical density was measured at 570 nm in a plate reader.

### HIV Replication Assay

U1, J1.1 and CD4^+^T cells were pre-treated overnight with multiple concentrations of DNA-PK inhibitors, prior to stimulation with 10 ng/ml of TNF-α for 56 hours more. HIV replication was determined by the reverse transcriptase (RT) assay as previously described in (21). Briefly, the cell supernatants were incubated overnight at 37° C with RT buffer (1 mM DTT, 5 mM MgCl_2_, 50 mM Tris-HCL, 20 mM of KCl, poly A, poly(dT), 0.1% Triton and [3H]TTP). The samples were then spotted on DEAE Filter mat paper and washed with 5% disodium phosphate (Na_2_HPO_4_) and water. The samples were dried completely and the activity was read in a Betaplate counter (Wallac, Gaithersburg, MA). For replication assay in MT-4 cells, the cells were pre-treated for 24 hours with the DNA-PK inhibitors Nu7026 or Nu7441 at different concentrations and DMSO as control, and seeded (10^5^ cells/well) in 24 well plate in 1ml of RPMI 10% FBS. Next day, the cells were infected with 10 μL of viral stock X4_LAI.04_ (0.7-ng/mL p24gag antigen) with X4_LAI.04_ and culture medium was replaced with fresh RPMI with DNA-PK inhibitors. Thus, inhibitors were present during the entire culture period. After 3 days, HIV-1 replication in MT-4 was evaluated by measuring the levels of p24gag in cell culture medium using a dynamic immunofluorescent cytometric bead assay (Luminex).

### Western Blot

Jurkat and U937 cells (wild type or DNA-PK knockdown) were plated at 5 x 10^6^/ml in Petri dishes. The wild type cells were incubated with 8μM, 12 μM and 16 μM of Nu7441 overnight. Cells were lysed with 0.5% NP-40 (supplemented with 10 mM HEPES-KOH pH 7.9, 60 mM KCL, 1 mM EDTA, 1 mM DTT, 1 mM phenylmethylsulfonylfluoride (PMSF), 10 μg/mL leupeptin, 10 μg/mL aprotinin. The isolated nuclei were then lysed in a buffer comprising 250 mM Tris pH7.8, 60 mM KCL, 1 mM EDTA, 1 mM DTT, 1 mM PMSF, 10 μg/mL leupeptin, 10 μg/mL aprotinin, 100 mM NaF and 200 μM sodium orthovanadate with three freeze-thaw cycles. Protein samples were separated by electrophoresis in SDS-PAGE, transferred to a nitrocellulose membrane and detected with specific antibodies. The immunoreactive proteins, after incubation with appropriately labeled secondary antibodies, were detected with an enhanced chemiluminescence detection kit. Densitometric analysis was done with ImageJ software.

### Reverse transcriptase-(RT)-qPCR

Total RNA was isolated from cells using a High Pure RNA Isolation Kit (Roche) or Trizol (ThermoFisher). The mRNA was reverse transcribed into complementary DNA (cDNA) using OneTaq RT-PCR Kit (New England Biolabs) or oligo dT primer. cDNA was quantified via Real-time quantitative PCR (RT-qPCR), which was performed in CFX96 Real-time PCR detection system (Bio-Rad) using SYBR green reaction mix (Bio-Rad). The data were normalized to the corresponding values for glyceraldehyde-3-phosphate dehydrogenase (GAPDH). The Nuc-2 primer sequences are described in the ChIP section.

### Chromatin immunoprecipitation (ChIP) Assays and q-PCR

ChIP assays were performed using a previously described protocol (59, 60). For cell stimulation, we either used 10 ng/ml TNF-α (for cell lines) or 25 μl per 10^6^ cells of α-CD3/CD28 antibodies bound Dynal beads along with 20 U/ml of IL-2 (for primary T cells). The chromatin was immunoprecipitated using different antibodies, detailed in the antibodies section. Each sample (5%) was analyzed by quantitative real-time PCR (qPCR) to assess the amount of sample immunoprecipitated by an individual antibody. Control preimmune IgG value was subtracted from each sample value to remove the background counts. SYBR green PCR master mix (12.5 μl/sample; Bio-Rad) combined with 1 μl of each primer, 5 μl of ChIPed DNA and water to a final volume 25 μl was analyzed by q-PCR. The primers used were the following (numbered with respect to the transcription start site): Promoter region of HIV-1 LTR (promoter) forward,-116, AGC TTG CTA CAA GGG ACT TTC C and reverse +4, ACC CAG TAC AGG CAA AAA GCA G; Nucleosome-1 region HIV-1 LTR (Nuc-1) forward +30, CTG GGA GCT CTC TGG CTA ACT A and reverse +134, TTA CCA GAG TCA CAC AAC AGA CG; Nucleosome-2 region HIV-1 LTR (Nuc-2) forward +283F, GACTGGTGAGTACGCCAAAAAT and reverse +390R, TTTCCCATCGCGATCTAATTC glyceraldehyde 3-phosphate dehydrogenase (GAPDH) promoter forward, –125, CAC GTA GCT CAG GCC TCA AGA C and reverse, –10, AGG CTG CGG GCT CAA TTT ATA G; GAPDH was also assessed by forward, –145, TAC TAG CGG TTT TAC GGG CG and reverse, +21 TCG AAC AGG AGG AGC AGA GAG CGA.

### Antibodies

The following antibodies where used in this study: RNAP II (17-672, Millipore; 61667, Active Motif; or sc-899, Santa Cruz), p-RNAP II (Ser2) (13499, Cell Signaling; 3E10, Active Motif, ab5095, Abcam), p-RNAP II (Ser5) (61085, Active Motif; 13523, Cell Signaling), H3-Ac (Upstate 07-593), H3K27me3 (07-449; Upstate), DNA-PKcs (Sc-9051, Santa Cruz), p-DNA-PKcs (S2056) (ab18192, Abcam), TRIM28 (A300-274A, Bethyl), p-TRIM28 (S824) (A300-767A, Bethyl), Tat (ab6539, Abcam), Cyclin T1 (sc-8128, Santa Cruz), p65 (sc-372 and sc-514451 Santa Cruz), CDK9 (sc-13130, sc-484), CDK7 (A300-405A, Bethyl), HDAC1 (sc-7872 and sc-6298, Santa Cruz), HDAC3 (sc-11417 and sc-376957 Santa Cruz), β-Actin (sc-47778 Santa Cruz).

### Flow cytometric analysis

Jurkat and U937 cells and corresponding DNA-PK knockdown clones were infected with VSV-G pseudotyped replication-incompetent HIV carrying *GFP* gene. After 48 to 72 hours, the expression of the fluorescent reporter, GFP, was assessed through fluorescence-activated cell sorting (FACS) using a FACSCalibur flow cytometer. Data were analyzed using Flowjo software.

### Statistical analysis

Data were analyzed using Microsoft Excel or GraphPad Prism and ImageJ softwares. For paired samples, statistical analyses were performed using Student’s *t* test. Statistical comparisons between the control and tested groups were analyzed using the one-way ANOVA. The p value of statistical significance was set at either; p < 0.05 (*), 0.01 (**), 0.001 (***), or 0.0001 (****). All experiments were independently repeated at least three times.

## Authors’ contributions

Research designed: MT; Research performed: MT, SZ, LD, GSahu, LS, TJ, and HY; Data analysed: MT, SZ, AO, LS, GSahu, LD, GSimon and MB; Manuscript written: MT, SZ, GSimon, AO and MB. All authors read and approved the final manuscript.

## Acknowledgements

We thank the AIDS Research and Reference Reagent Program, Division of AIDS, National Institute of Allergy and Infectious Diseases, US National Institutes of Health; M. Gately (Hoffmann La Roche) for human recombinant interleukin 2; We are also thankful to the Flow Cytometry core facility of George Washington University. We also want to thank the reviewers for providing constructive criticisms prior to publication. The following reagents were obtained through the AIDS Research and Reference Reagent Program, Division of AIDS, NIAID, National Institutes of Health: U1, J1.1. We are highly thankful to Dr. Kalamo Farley for his efforts in standardizing some of the assay conditions.

## References

1. An, J., T. Yang, Y. Huang, F. Liu, J. Sun, Y. Wang, Q. Xu, D. Wu, and P. Zhou. 2011. Strand-specific PCR of UV radiation-damaged genomic DNA revealed an essential role of DNA-PKcs in the transcription-coupled repair. BMC Biochem 12:2.

2. Anderson, C. W., and T. H. Carter. 1996. The DNA-activated protein kinase - DNA-PK. Curr Top Microbiol Immunol 217:91–111.

3. Bourgeois, C. F., Y. K. Kim, M. J. Churcher, M. J. West, and J. Karn. 2002. Spt5 cooperates with Tat by preventing premature RNA release at terminator sequences. Mol. Cell. Biol. 22:1079–1093.

4. Bruner, K. M., A. J. Murray, R. A. Pollack, M. G. Soliman, S. B. Laskey, A. A. Capoferri, J. Lai, M. C. Strain, S. M. Lada, R. Hoh, Y. C. Ho, D. D. Richman, S. G. Deeks, J. D. Siliciano, and R. F. Siliciano. 2016. Defective proviruses rapidly accumulate during acute HIV-1 infection. Nat Med 22:1043–1049.

5. Budhiraja, S., and A. P. Rice. 2013. Reactivation of latent HIV: do all roads go through P-TEFb Future Virol 8:649–659.

6. Bunch, H., and S. K. Calderwood. 2015. TRIM28 as a novel transcriptional elongation factor. BMC molecular biology 16.

7. Bunch, H., B. P. Lawney, Y. F. Lin, A. Asaithamby, A. Murshid, Y. E. Wang, B. P. Chen, and S. K. Calderwood. 2015. Transcriptional elongation requires DNA break-induced signalling. Nature communications 6:10191.

8. Bunch, H., X. F. Zheng, A. Burkholder, S. T. Dillon, S. Motola, G. Birrane, C. C. Ebmeier, S. Levine, D. Fargo, G. Hu, D. J. Taatjes, and S. K. Calderwood. 2014. TRIM28 regulates RNA polymerase II promoter-proximal pausing and pause release. Nat Struct Mol Biol 21:876–883.

9. Calderwood, S. K. 2016. Creative damage unleashes transcription. Cell cycle 15:1021–1022.

10. Carter, T., I. Vancurova, I. Sun, W. Lou, and S. DeLeon. 1990. A DNA-activated protein kinase from HeLa cell nuclei. Molecular and cellular biology 10:6460–6471.

11. Chun, T. W., R. T. Davey, Jr., D. Engel, H. C. Lane, and A. S. Fauci. 1999. Re-emergence of HIV after stopping therapy. Nature 401:874–875.

12. Cornell, L., J. Munck, N. Curtin, and H. Reeves. 2012. DNA-Pk or Atm Inhibition Inhibits Non-Homologous End Joining and Enhances Chemo- and Radio Sensitivity in Hepatocellular Cancer Cell Lines. Gut 61:A201–A201.

13. Dahmus, M. E. 1995. Phosphorylation of the C-terminal domain of RNA polymerase II. Biochimica et biophysica acta 1261:171–182.

14. Davey, R. T., Jr., N. Bhat, C. Yoder, T. W. Chun, J. A. Metcalf, R. Dewar, V. Natarajan, R. A. Lempicki, J. W. Adelsberger, K. D. Miller, J. A. Kovacs, M. A. Polis, R. E. Walker, J. Falloon, H. Masur, D. Gee, M. Baseler, D. S. Dimitrov, A. S. Fauci, and H. C. Lane. 1999. HIV-1 and T cell dynamics after interruption of highly active antiretroviral therapy (HAART) in patients with a history of sustained viral suppression. Proc. Natl. Acad. Sci. U S A 96:15109–15114.

15. Davidson, D., L. Amrein, L. Panasci, and R. Aloyz. 2013. Small Molecules, Inhibitors of DNA-PK, Targeting DNA Repair, and Beyond. Front Pharmacol 4:5.

16. Dvir, A., L. Y. Stein, B. L. Calore, and W. S. Dynan. 1993. Purification and characterization of a template-associated protein kinase that phosphorylates RNA polymerase II. The Journal of biological chemistry 268:10440–10447.

17. Falck, J., J. Coates, and S. P. Jackson. 2005. Conserved modes of recruitment of ATM, ATR and DNA-PKcs to sites of DNA damage. Nature 434:605–611.

18. Garber, M. E., T. P. Mayall, E. M. Suess, J. Meisenhelder, N. E. Thompson, and K. A. Jones. 2000. CDK9 autophosphorylation regulates high-affinity binding of the human immunodeficiency virus type 1 Tat-P-TEFb comples to TAR RNA. Mol. Cell. Biol. 20:6958–6969.

19. Geuting, V., C. Reul, and M. Lobrich. 2013. ATM Release at Resected Double-Strand Breaks Provides Heterochromatin Reconstitution to Facilitate Homologous Recombination. Plos Genet 9.

20. Gottlieb, T. M., and S. P. Jackson. 1993. The DNA-dependent protein kinase: requirement for DNA ends and association with Ku antigen. Cell 72:131–142.

21. Guendel, I., S. Iordanskiy, R. Van Duyne, K. Kehn-Hall, M. Saifuddin, R. Das, E. Jaworski, G. C. Sampey, S. Senina, L. Shultz, A. Narayanan, H. Chen, B. Lepene, C. Zeng, and F. Kashanchi. 2014. Novel neuroprotective GSK-3beta inhibitor restricts Tat-mediated HIV-1 replication. J Virol 88:1189-1208.

22. Hardcastle, I. R., X. Cockcroft, N. J. Curtin, M. D. El-Murr, J. J. Leahy, M. Stockley, B. T. Golding, L. Rigoreau, C. Richardson, G. C. Smith, and R. J. Griffin. 2005. Discovery of potent chromen-4-one inhibitors of the DNA-dependent protein kinase (DNA-PK) using a small-molecule library approach. J Med Chem 48:7829–7846.

23. Hartley, K. O., D. Gell, G. C. Smith, H. Zhang, N. Divecha, M. A. Connelly, A. Admon, S. P. Lees-Miller, C. W. Anderson, and S. P. Jackson. 1995. DNA-dependent protein kinase catalytic subunit: a relative of phosphatidylinositol 3-kinase and the ataxia telangiectasia gene product. Cell 82:849–856.

24. Hazra, J., P. Mukherjee, A. Ali, S. Poddar, and M. Pal. 2017. Engagement of Components of DNA-Break Repair Complex and NF kappa B in H5p70A1A Transcription Upregulation by Heat Shock. PloS one 12.

25. Ho, Y. C., L. Shan, N. N. Hosmane, J. Wang, S. B. Laskey, D. I. Rosenbloom, J. Lai, J. N. Blankson, J. D. Siliciano, and R. F. Siliciano. 2013. Replication-competent noninduced proviruses in the latent reservoir increase barrier to HIV-1 cure. Cell 155:540–551.

26. Isel, C., and J. Karn. 1999. Direct evidence that HIV-1 Tat stimulates RNA polymerase II carboxyl-terminal domain hyperphosphorylation during transcriptional elongation. J Mol Biol 290:929–941.

27. Karn, J. 2011. The molecular biology of HIV latency: breaking and restoring the Tat-dependent transcriptional circuit. Current opinion in HIV and AIDS 6:4–11.

28. Kim, Y. K., C. F. Bourgeois, C. Isel, M. J. Churcher, and J. Karn. 2002. Phosphorylation of the RNA polymerase II carboxyl-terminal domain by CDK9 is directly responsible for human immunodeficiency virus type 1 Tat-activated transcriptional elongation. Mol. Cell. Biol. 22:4622–4637.

29. Kim, Y. K., C. F. Bourgeois, R. Pearson, M. Tyagi, M. J. West, J. Wong, S. Y. Wu, C. M. Chiang, and J. Karn. 2006. Recruitment of TFIIH to the HIV LTR is a rate-limiting step in the emergence of HIV from latency. EMBO J. 25:3596–3604.

30. Leahy, J. J. J., B. T. Golding, R. J. Griffin, I. R. Hardcastle, C. Richardson, L. Rigoreau, and G. C. M. Smith. 2004. Identification of a highly potent and selective DNA-dependent protein kinase (DNA-PK) inhibitor (NU7441) by screening of chromenone libraries. Bioorganic & Medicinal Chemistry Letters 14:6083–6087.

31. Lees-Miller, S. P., Y. R. Chen, and C. W. Anderson. 1990. Human cells contain a DNA-activated protein kinase that phosphorylates simian virus 40 T antigen, mouse p53, and the human Ku autoantigen. Molecular and cellular biology 10:6472–6481.

32. Lees-Miller, S. P., R. Godbout, D. W. Chan, M. Weinfeld, R. S. Day, 3rd, G. M. Barron, and J. Allalunis-Turner. 1995. Absence of p350 subunit of DNA-activated protein kinase from a radiosensitive human cell line. Science 267:1183–1185.

33. Li, X., Y. K. Lee, J. C. Jeng, Y. Yen, D. C. Schultz, H. M. Shih, and D. K. Ann. 2007. Role for KAP1 serine 824 phosphorylation and sumoylation/desumoylation switch in regulating KAP1-mediated transcriptional repression. The Journal of biological chemistry 282:36177–36189.

34. Li, Y., X. Wang, P. Yue, H. Tao, S. S. Ramalingam, T. K. Owonikoko, X. Deng, Y. Wang, H. Fu, F. R. Khuri, and S. Y. Sun. 2013. Protein phosphatase 2A and DNA-dependent protein kinase are involved in mediating rapamycin-induced Akt phosphorylation. J Biol Chem 288:13215–13224.

35. Lisco, A., J. C. Grivel, A. Biancotto, C. Vanpouille, F. Origgi, M. S. Malnati, D. Schols, P. Lusso, and L. B. Margolis. 2007. Viral interactions in human lymphoid tissue: Human herpesvirus 7 suppresses the replication of CCR5-tropic human immunodeficiency virus type 1 via CD4 modulation. Journal of virology 81:708–717.

36. Liu, H., C. H. Herrmann, K. Chiang, T. L. Sung, S. H. Moon, L. A. Donehower, and A. P. Rice. 2010. 55K isoform of CDK9 associates with Ku70 and is involved in DNA repair. Biochem Biophys Res Commun 397:245–250.

37. Maldonado, E., R. Shiekhattar, M. Sheldon, H. Cho, R. Drapkin, P. Rickert, E. Lees, C. W. Anderson, S. Linn, and D. Reinberg. 1996. A human RNA polymerase II complex associated with SRB and DNA-repair proteins. Nature 381:86–89.

38. Margolis, D. M. 2010. Mechanisms of HIV latency: an emerging picture of complexity. Current HIV/AIDS reports 7:37–43.

39. Mbonye, U., and J. Karn. 2011. Control of HIV latency by epigenetic and non-epigenetic mechanisms. Current HIV research.

40. McNamara, R. P., J. E. Reeder, E. A. McMillan, C. W. Bacon, J. L. McCann, and I. D'Orso. 2016. KAP1 Recruitment of the 7SK snRNP Complex to Promoters Enables Transcription Elongation by RNA Polymerase II. Molecular cell 61:39–53.

41. Nagasawa, M., F. Watanabe, A. Suwa, K. Yamamoto, K. Tsukada, and H. Teraoka. 1997. Nuclear translocation of the catalytic component of DNA-dependent protein kinase upon growth stimulation in normal human T lymphocytes. Cell Structure and Function 22:585–594.

42. Orphanides, G., T. Lagrange, and D. Reinberg. 1996. The general transcription factors of RNA polymerase II. Genes Dev. 10:2657–2683.

43. Pearson, R., Y. K. Kim, J. Hokello, K. Lassen, J. Friedman, M. Tyagi, and J. Karn. 2008. Epigenetic silencing of human immunodeficiency virus (HIV) transcription by formation of restrictive chromatin structures at the viral long terminal repeat drives the progressive entry of HIV into latency. J. Virol. 82:12291–12303.

44. Peddi, P., C. W. Loftin, J. S. Dickey, J. M. Hair, K. J. Burns, K. Aziz, D. C. Francisco, M. I. Panayiotidis, O. A. Sedelnikova, W. M. Bonner, T. A. Winters, and A. G. Georgakilas. 2010. DNA-PKcs deficiency leads to persistence of oxidatively induced clustered DNA lesions in human tumor cells. Free Radic Biol Med 48:1435–1443.

45. Peterlin, B. M., and D. H. Price. 2006. Controlling the elongation phase of transcription with P-TEFb. Mol. Cell 23:297–305.

46. Pollack, R. A., R. B. Jones, M. Pertea, K. M. Bruner, A. R. Martin, A. S. Thomas, A. A. Capoferri, S. A. Beg, S. H. Huang, S. Karandish, H. P. Hao, E. Halper-Stromberg, P. C. Yong, C. Kovacs, E. Benko, R. F. Siliciano, and Y. C. Ho. 2017. Defective HIV-1 Proviruses Are Expressed and Can Be Recognized by Cytotoxic T Lymphocytes, which Shape the Proviral Landscape. Cell Host Microbe 21:494-+.

47. Price, D. H. 2000. P-TEFb, a cyclin-dependent kinase controlling elongation by RNA polymerase II. Mol. Cell. Biol. 20:2629–2634.

48. Reeves, W. H., and Z. M. Sthoeger. 1989. Molecular cloning of cDNA encoding the p70 (Ku) lupus autoantigen. The Journal of biological chemistry 264:5047–5052.

49. Rice, A. P., and M. B. Mathews. 1988. Transcriptional but not translational regulation of HIV-1 by the *tat* gene product. Nature 332:551–553.

50. Riedl, T., and J.-M. Egly. 2000. Phosphorylation in transcription: the CTD and more. Gene Expr. 9:3–13.

51. Sahu, G., K. Farley, N. El-Hage, B. Aiamkitsumrit, R. Fassnacht, F. Kashanchi, A. Ochem, G. L. Simon, J. Karn, K. F. Hauser, and M. Tyagi. 2015. Cocaine promotes both initiation and elongation phase of HIV-1 transcription by activating NF-kappaB and MSK1 and inducing selective epigenetic modifications at HIV-1 LTR. Virology 483:185–202.

52. Saunders, A., L. J. Core, and J. T. Lis. 2006. Breaking barriers to transcription elongation. Nat. Rev. Mol. Cell Biol. 7:557–567.

53. Siliciano, R. F., and W. C. Greene. 2011. HIV Latency. Cold Spring Harb Perspect Med 1:a007096.

54. Stein, R. C. 2001. Prospects for phosphoinositide 3-kinase inhibition as a cancer treatment. Endocr Relat Cancer 8:237–248.

55. Taccioli, G. E., T. M. Gottlieb, T. Blunt, A. Priestley, J. Demengeot, R. Mizuta, A. R. Lehmann, F. W. Alt, S. P. Jackson, and P. A. Jeggo. 1994. Ku80: product of the XRCC5 gene and its role in DNA repair and V(D)J recombination. Science 265:1442–1445.

56. Tomimatsu, N., B. Mukherjee, and S. Burma. 2009. Distinct roles of ATR and DNA-PKcs in triggering DNA damage responses in ATM-deficient cells. EMBO reports 10:629–635.

57. Trigon, S., H. Serizawa, J. W. Conaway, R. C. Conaway, S. P. Jackson, and M. Morange. 1998. Characterization of the residues phosphorylated in vitro by different C-terminal domain kinases. The Journal of biological chemistry 273:6769–6775.

58. Tyagi, M., and M. Bukrinsky. 2012. Human immunodeficiency virus (HIV) latency: the major hurdle in HIV eradication. Mol Med 18:1096–1108.

59. Tyagi, M., and J. Karn. 2007. CBF-1 promotes transcriptional silencing during the establishment of HIV-1 latency. EMBO J. 26:4985–4995.

60. Tyagi, M., R. J. Pearson, and J. Karn. 2010. Establishment of HIV latency in primary CD4+ cells is due to epigenetic transcriptional silencing and P-TEFb restriction. Journal of virology 84:6425–6437.

61. Tyagi, S., A. Ochem, and M. Tyagi. 2011. DNA-PK functionally interacts with RNA polymerase II complex recruited at HIV LTR and play important role in HIV gene expression. J Gen Virol. 35

62. Van Lint, C., A. Burny, and E. Verdin. 1991. The intragenic enhancer of human immunodeficiency virus type 1 contains functional AP-1 binding sites. Journal of virology 65:7066–7072.

63. Van Lint, C., S. Emiliani, M. Ott, and E. Verdin. 1996. Transcriptional activation and chromatin remodeling of the HIV-1 promoter in response to histone acetylation. EMBO J. 15:1112–1120.

64. Verdin, E. 1991. DNase I-hypersensitive sites are associated with both long terminal repeats and with the intragenic enhances of integrated human immunodeficiency virus type 1. J. Virol. 65:6790–6799.

65. Ward, J. F. 1988. DNA damage produced by ionizing radiation in mammalian cells: identities, mechanisms of formation, and reparability. Prog Nucleic Acid Res Mol Biol 35:95–125.

66. Willmore, E., S. de Caux, N. J. Sunter, M. J. Tilby, G. H. Jackson, C. A. Austin, and B. W. Durkacz. 2004. A novel DNA-dependent protein kinase inhibitor, NU7026, potentiates the cytotoxicity of topoisomerase II poisons used in the treatment of leukemia. Blood 103:4659–4665.

67. Yamamoto, S., Y. Watanabe, P. J. van der Spek, T. Watanabe, H. Fujimoto, F. Hanaoka, and Y. Ohkuma. 2001. Studies of nematode TFIIE function reveal a link between Ser-5 phosphorylation of RNA polymerase II and the transition from transcription initiation to elongation. Molecular and cellular biology 21:1–15.

68. Yaneva, M., J. Wen, A. Ayala, and R. Cook. 1989. cDNA-derived amino acid sequence of the 86-kDa subunit of the Ku antigen. The Journal of biological chemistry 264:13407–13411.

69. You, H., M. M. Kong, L. P. Wang, X. Xiao, H. L. Liao, Z. Y. Bi, H. Yan, H. Wang, C. H. Wang, Q. Ma, Y. Q. Liu, and Y. Y. Bi. 2013. Inhibition of DNA-dependent protein kinase catalytic subunit by small molecule inhibitor NU7026 sensitizes human leukemic K562 cells to benzene metabolite-induced apoptosis. J Huazhong Univ Sci Technolog Med Sci 33:43–50.

70. Yukl, S., S. Pillai, P. Li, K. Chang, W. Pasutti, C. Ahlgren, D. Havlir, M. Strain, H. Gunthard, D. Richman, A. P. Rice, E. Daar, S. Little, and J. K. Wong. 2009. Latently-infected CD4+ T cells are enriched for HIV-1 Tat variants with impaired transactivation activity. Virology 387:98–108.

71. Zhang, S. M., H. Zhang, T. Y. Yang, T. Y. Ying, P. X. Yang, X. D. Liu, S. J. Tang, and P. K. Zhou. 2014. Interaction between HIV-I Tat and DNA-PKcs modulates HIV transcription and class switch recombination. Int J Biol Sci 10:1138–1149.

72. Zhou, X., X. Zhang, Y. Xie, K. Tanaka, B. Wang, and H. Zhang. 2013. DNA-PKcs inhibition sensitizes cancer cells to carbon-ion irradiation via telomere capping disruption. PLoS One 8:e72641.

